# *Trypanosoma congolense* Variant Surface Glycoprotein gene expression occurs in the absence of monoallelic transcription control

**DOI:** 10.64898/2026.07.31.741978

**Authors:** Jane C Munday, Marija Krasiļņikova, Catarina A Marques, Stephen D Larcombe, Pieter Monsieurs, Guy Oldrieve, Kevin O Kidambasi, Emma M Briggs, Craig Lapsley, Andrew P Jackson, Jan Van Den Abbeele, Liam J Morrison, Keith R Matthews, Richard McCulloch

## Abstract

Antigenic variation is a very widespread process for pathogen evasion of mammalian adaptive immunity, involving the continuous change of exposed antigens. In many single-celled pathogens, antigen expression during antigenic variation is monoallelic: just a single gene from a large family is expressed in one cell at a time. In African trypanosomes, antigenic variation relies on expression of Variant Surface Glycoprotein (VSG), and in *Trypanosoma brucei* there is detailed understanding of the machinery that dictates that only one of approximately 15 VSG expression sites is actively transcribed at a time. In the closely related African trypanosome, *T. congolense*, which remains a significant blight on agriculture productivity in sub-Sharan Africa, we have no such understanding of *VSG* gene expression control or dynamics. Here, we have investigated the mechanics of antigenic variation in *T. congolense*, first examining the patterns of VSG expression at the transcript and protein level in small parasite populations *in vitro*. Surprisingly, this analysis revealed much greater VSG diversity than seen in *T. brucei*, with such expression diversity unaltered in mutants that impair homologous recombination. Using single cell transcriptomics, we explain this diversity, since we find no evidence for monoallelic transcription of *T. congolense VSG*s, but show instead that each parasite can dynamically express up to ∼40 different *VSG*s in a single cell both *in vitro* and *in vivo*. *VSG* co-expression occurs from *VSG* genes distributed across the genome, indicating the lack of a dedicated locus for *VSG* transcription. Thus, comparing two trypanosome species that rely on the same class of surface antigen for immune evasion has revealed highly distinct mechanisms for controlling antigen gene expression, challenging the assumed common operation of antigenic variation across African trypanosome species.

## Introduction

All pathogens of mammals must survive immune attack. In single-celled pathogens, including bacteria, fungi and protozoa, a common route to evade adaptive immunity and thereby sustain infections and promote transmission is antigenic variation: the periodic switching of exposed surface antigens^1^. Though the underlying mechanisms of antigenic variation vary across pathogens, a common feature of the operation of antigenic variation is monoallelic antigen expression, or allelic exclusion, where each cell expresses just a single antigen at a time^2–4^. For bacterial pathogens, such as *Neisseria*^5^, monoallelic expression is achieved by their genomes containing a single site for antigen transcription, with antigen variant genes moved to that site from a silent archive by recombination. In contrast, the malaria parasite *Plasmodium falciparum* encodes around 60 *var* genes that can be transcribed *in situ*, with controls having evolved to ensure that just one *var* is transcribed at a time^6,7^. African trypanosomes rely on expression and switching between Variant Surface Glycoproteins (VSGs) for antigenic variation, and our understanding of the operation of this process is dominated by decades of study in *Trypanosoma brucei*. Here, we have investigated the mechanics of antigenic variation in the closely related African trypanosome *Trypanosoma congolense*, focusing on the control of *VSG* gene expression in this important animal pathogen, which remains one of the biggest infectious disease constraints on agricultural productivity across Sub-Saharan Africa^8,9^.

Antigenic variation in *T. brucei* relies on two genomic features: a repertoire of >2000 silent *VSG* genes and pseudogenes^10–12^, and telomeric VSG expression sites (ESs) in which the *VSG* is co-transcribed with a number of Expression Site Associated Genes (*ESAGs*) from an RNA Polymerase (Pol) I promoter^13^. The most common route by which a change in the expressed VSG occurs, allowing evasion of anti-VSG immunity, is through recombination reactions focused on the actively transcribed ES^14^, wherein the ES-resident *VSG* is replaced by an antigenically distinct copy from the large silent *VSG* archive. Such *VSG* recombination is mainly catalysed by DNA break repair through homologous recombination (HR), as evidenced by experiments showing that loss of RAD51^14,15^ or its mediators, including BRCA2^16^, impairs VSG switching and diversification^17^. The output of *VSG* recombination is considerable expressed VSG diversity that arises in a hierarchy during infections^18–20^ (Larcombe et al, BIORXIV/2026/741217). Recombination is not, however, the only route for VSG switching in *T. brucei*, since the genome does not contain a single ES used for *VSG* transcription in the mammal, but approximately 15. Just one ES is selected for active transcription in a cell at a time, with all others inactive, and the rigour of VSG ES monoallelic transcription in wildtype *T. brucei* underlined by the failure of several experimental approaches that sought to isolate cells that naturally and stably co-transcribe more than one ES^21–24^. A growing body of work has now revealed the *T. brucei* machinery that dictates ES monoallelic transcription, with the earliest described factors being the VSG exclusion complex (VEX)^25^. Two key components of VEX are the proteins VEX1^26^ and VEX2^27–29^, which interact with the active ES as part of a subnuclear compartment termed the expression site body (ESB)^30^, which includes an extranucleolar pool of RNA Pol I and several other ESB-specific and -associated proteins^31–33^. Functional analyses point to differing roles of these factors in monoallelic control of ES transcription: RNAi against VEX1 or VEX2 leads to increased transcript levels from normally silent ESs, resulting in expression of multiple antigenically distinct VSG transcripts and proteins in a cell^26,29^; in contrast, RNAi against ESB1 results in loss of transcripts from active and inactive ESs^31^, whilst ESBX RNAi leads to both loss of transcripts from the active ES and increased transcripts from silent ESs^32^.

Recently there has been considerable progress in our ability to grow^34^ and genetically manipulate^35^*T. congolense*^9^, allowing us to begin to assess the operation of antigenic variation in this species, which is a dominant contributor to animal African trypanosomiasis in sub-Saharan Africa. Although*T. congolense* encodes VSGs and is believed to survive in the mammalian host through antigenic variation^36,37^, our understanding of VSG expression dynamics and control is substantially more limited than in *T. brucei*. Expression of new VSGs as a response to immune pressure has been documented, either following infection, drug cure and then challenge^38^, or using monoclonal antibodies against specific VSGs^39^, but the mechanisms behind such change are untested. Duplication of a *T. congolense VSG* between telomeres has been described, with the generation of a duplicated copy associated with expression^40^, but a mechanistic demonstration that such recombination provides for immune evasion is lacking, including a test of whether or not HR acts to direct changes in *T. congolense* VSG expression. In this regard, the *VSG* repertoires of *T. brucei* and *T. congolense* appear distinct: whereas *T. brucei VSG*s are separated into a- and b-types (based on cysteine residue distribution)^41^ that undergo seemingly unconstrained recombination, *T. congolense* contains only b-type *VSG*s that can be separated into discrete phylotypes, with recombination considered to be largely limited to within a phylotype^36,42^. What impact such a constraint might have on antigenic variation has not been assessed. Importantly, transcription of *T. congolense VSG*s remains barely studied: *T. brucei*-like *VSG* ESs, including *ESAG*s, 70 bp repeats and upstream RNA Pol I promoters, have not been detected in the *T. congolense* genome^43^, including in a recent telomere-telomere *de novo* genome assembly (Krasilnikova et al, BioRXiv10.64898/2026.02.19.706783). Intriguingly, and consistent with an earlier report^44^, transcripts could be mapped to *VSG*s present in

*T. congolense* minichromosomes (Krasilnikova et al, BioRXiv10.64898/2026.02.19.706783), a compartment of the genome that in *T. brucei* appears to harbour only silent telomeric *VSG*s^45^. Such expression may be consistent with predictions of minichromosomal *T. congolense* ES facsimiles, comprising telomeric *VSG*s and some upstream conserved sequences^43^. However, no work has tested for monoallelic expression of *VSG*s in *T. congolense*, which might be predicted if *T. brucei-*like ESs are present.

Here, we document that the diversity of *VSG* transcripts expressed in both cultured *T. congolense* and in cells recovered from mice substantially exceeds that seen in *T. brucei*, a phenotype that is not the result of HR between *VSG*s nor extreme antigenic diversity between individuals in the population but instead arises due to the absence of monoallelic *VSG* expression in individual parasites. This work reveals a fundamental and hitherto unappreciated distinction in how two closely related African trypanosomes control the expression of a common variant surface antigen used for immune survival.

## Results

### *T. congolense* displays considerable *VSG* transcript diversity that is not dependent on homologous recombination

To begin to understand VSG expression dynamics in *T. congolense* and the influence of HR, we generated individual null mutants of *RAD51* and *BRCA2* in *T. congolense* IL3000 bloodstream form cells. The entire ORF from both alleles of each HR gene was successfully removed (Krasilnikova et al, BioRXiv10.64898/2026.02.19.706783), arguing that neither gene product is essential for *T. congolense in vitro*, similar to that reported for the corresponding mutants in *T. brucei*^14,16^. To determine the pattern of *VSG* expression after loss of the HR genes, we sub-cloned wild type (WT) parasites and both mutants and examined two clones of each following their outgrowth (approximately 38-46 population doublings) into starting populations (P0), which were then resampled after 11 and 22 serial passages *in vitro* (P11, P22; approximately 80-90 population doublings between each sampling; Fig.1A). Whole transcriptome RNA-seq data from each subclonal population (after limited expansion, in triplicate) was mapped using a Nanopore-Hi-C *de novo* assembly of the *T. congolense* IL3000 genome (Krasilnikova et al, BioRXiv10.64898/2026.02.19.706783), where *VSG* genes were identified using two filters: presence of Pfam domain PF13206, and Companion^46^ annotation as a *VSG*. We approached the expression analysis in two ways: first, the transcriptome-wide quantifier *Salmon*^47^ was used to map all reads to the genome (Fig.1B shows one clone for each cell type; Fig.S1 shows the other clone for each cell type); second, VSGseq2^48^ was used to perform *de novo* transcript assembly prior to mapping (Fig.S2). Irrespective of the approach, a very large number of distinct *VSG* transcripts were detected in each WT population: *Salmon* mapping predicted 404-507 *VSG*s (minimum transcripts per million (TPM) threshold, 5/VSG; Fig.1, Fig.S1), while VSGseq2 predicted 476-501 (Fig.S2). Altering the minimum TPM/VSG in the mapping changed the total number of *VSG* transcripts detected, but even at a very stringent threshold of 1000/VSG, extensive *VSG* transcript diversity was seen in the WT population (Fig.S3A). Similarly, applying a more stringent filter of what constitutes a *VSG*, by additionally requiring any predicted gene to be part of a defined phylotype^42,49^, did not substantially alter WT *VSG* transcript diversity (Fig.S3B). Although the relative abundance of individual *VSG* transcripts varied between the two WT clones and over time, *VSG* diversity did not (Fig.1B, Fig.S1). Moreover, *VSG* transcript diversity was not notably altered by RAD51 or BRCA2 loss (Fig.1B, Fig.S1; 376-437 and 361-440 *VSG*s, respectively), indicating diversity did not arise through recombination between *VSGs* via HR. To attempt to rule out the possibility of lab-specific issues (such as parasite growth conditions) we examined *VSG* expression in whole transcriptome data generated from *T. congolense* IL3000 bloodstream form cells grown independently in a different lab^35^. These data also revealed considerable *VSG* transcript diversity in four parasite populations (Fig.S4A) and a wide range of *VSG* transcript abundance (Fig.S4B)^36,37,49–51^.

**Figure 1.**
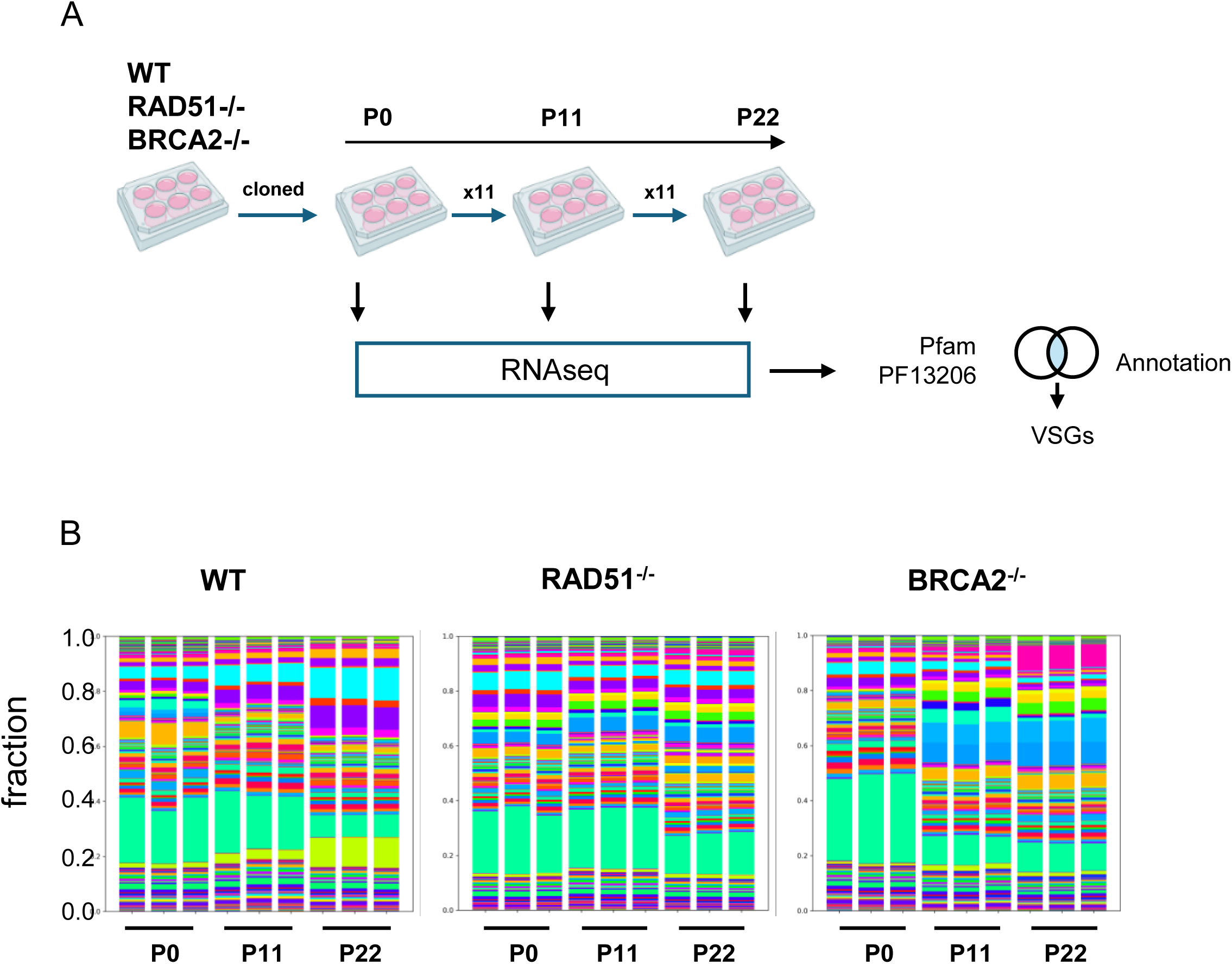
Diversity of *VSG* transcripts expressed over time in culture by wild type *T. congolense* or null mutants of *RAD51* or *BRCA2*. **A.** Outline of growth and sampling for RNA-seq. Wild type cells (WT), RAD51-/- mutants or BRCA2-/- mutants were cloned and then expanded in triplicate for RNA extraction (passage 0, P0), as well as grown for 11 or 22 passages (P11, P22) and RNA sampled in the same way. **B.** Bulk RNA-seq data from WT, RAD51^-/-^ and BRCA2^-/-^ cells mapped to the *T. congolense* transcriptome, showing the relative proportion of all detected *VSG* transcripts per sample, with each colour representing a distinct *VSG* gene. Populations at each of P0, P11 and P22 are shown as three replicates.

### *T. congolense VSG* diversity is present after limited growth *in vitro* and surpasses that seen in cultured *T. brucei*

To explore VSG expression diversity in *T. congolense* further, we used an adapted version of barcoded Nanopore sequencing of splice leader-enriched transcripts (SL-barseq), used previously to measure VSG switching and expression in *T. brucei*^52^. This approach provides a number of tests of the data above: cDNA was sequenced by long-read Nanopore sequencing, testing for potential mapping issues associated with short-read Illumina sequencing; VSG expression was evaluated in a larger number of clones; and the analysis was conducted more rapidly after cloning and from smaller populations (∼3x10^6^ cells, ∼22 population doublings from cloning, compared with ∼42 population doublings in the P0 samples in Fig.1). To approach this, we re-cloned WT cells from the P0 and P22 *T. congolense* populations, as well as from *RAD51* and *BRCA2* mutants at P22 and performed SL-barseq on 10-20 clones (Fig.2A). In every independent recently cloned *T. congolense* population, whether WT or HR mutant, we detected between 44 and 413 *VSG* transcripts (Fig.2B,C). This diversity is in striking contrast to the same analysis of bloodstream form *T. brucei* Lister 427 MITat1.2 parasites, where a single *VSG* (*VSG-2*, found in BES1^13^, which is the predominantly transcribed VSG ES in this *T. brucei* strain^28,53,54^) represented >95% of *VSG* transcripts in all five clones analysed (Fig.2D), consistent with previous independent SL-barseq^52^ analysis and population RNA-seq^13,28,53–55^.

**Figure 2.**
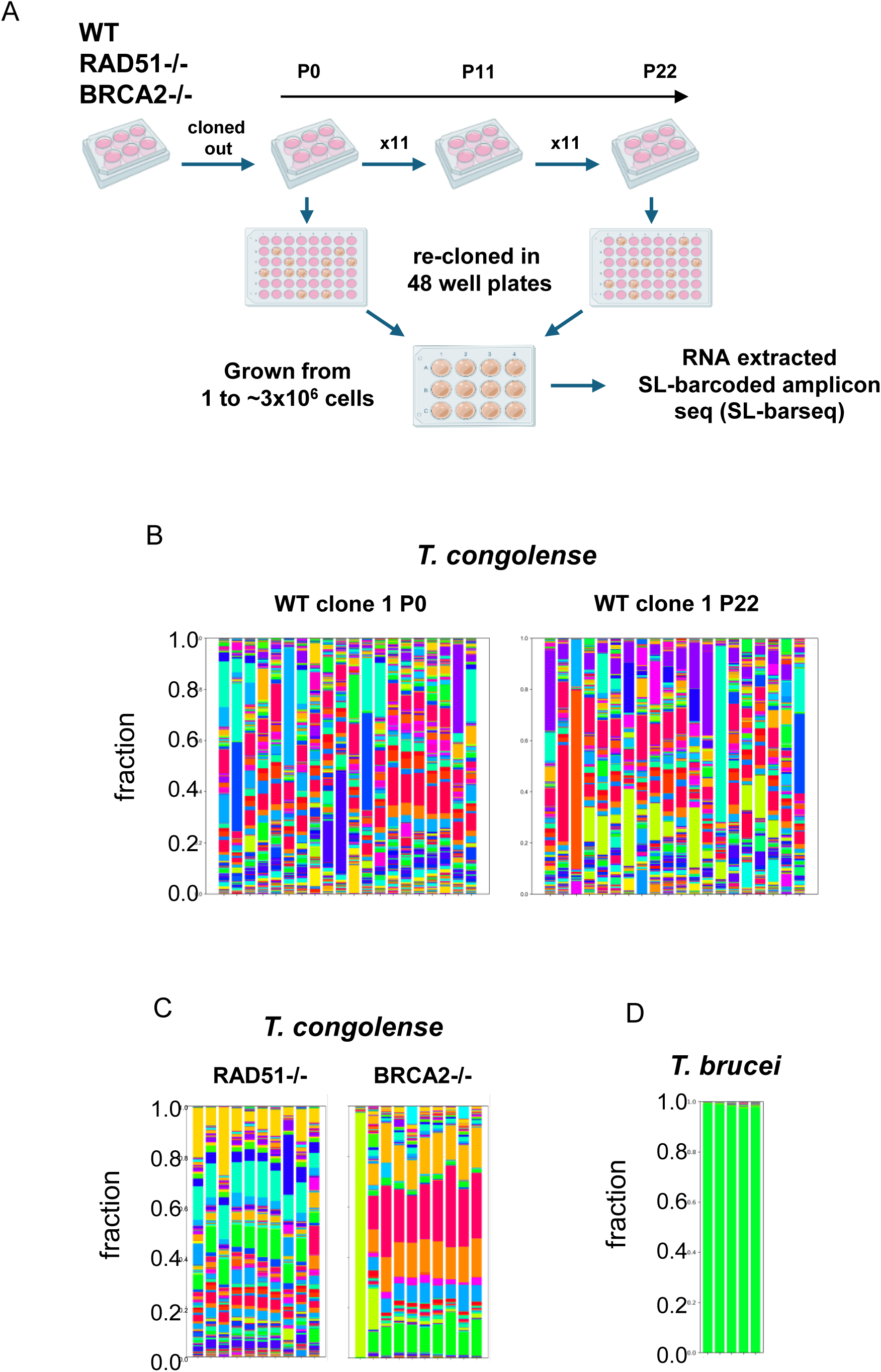
*VSG* transcript diversity is detectable soon after cloning of wild type *T. congolense* and *RAD51* or *BRCA2* null mutants. **A.** Illustration of the cloning of WT, RAD51^-/-^ and BRCA2^-/-^ *T. congolense* cells to give starting clones (P0), followed by a further 11 passages (P11) and then another 11 passages (P22). Two populations of each cell type at each passage number were recloned in 48 -well plates, then multiple sub-clones grown to ∼3x10^6^ cells and RNA extracted for SL-barcoded amplicon PCR analysis (SL-barseq). **B-D.** Long read SL-barseq data from, *T. congolense* WT clones (B), *T. congolense* RAD51^-/-^ and BRCA2^-/-^ clones (C) and Lister 427 *T. brucei* clones (D) mapped to the transcriptome, showing the relative proportion of all detected *VSG* transcripts per sample, with each colour representing a distinct VSG gene; only samples with at least 1000 total *VSG* reads are shown.

### *T. congolense VSG* transcript diversity is reflected in VSG protein diversity

To ask if the diversity of *VSG* transcripts seen in recently cloned populations of *T. congolense* is reflected at the protein level, we examined the profile of VSG proteins expressed in two of the recently cloned WT SL-barseq populations by surface VSG release^56^ (Fig.S5) and identification by Mass Spectrometry (Fig.3A). In both WT clones, we detected multiple distinct VSGs (92 or 70 in clones C10 or G03, respectively, using an Exponentially Modified Protein Abundance Index^57^ (emPAI) >0; 9 or 13 with emPAI >2.0). As seen in comparisons of transcriptomes and proteomes in *T. cruzi*^58^ and *T. brucei*^59^, fewer *T. congolense* VSG proteins were detected than *VSG* transcripts, and such a restriction in proteomic sensitivity relative to transcriptome analysis is consistent with the large majority of different VSG proteins detected also being seen in the repertoire of mapped *VSG* transcripts (Fig.3B). These data are in striking contrast to the near exclusive detection of a single VSG protein in cultured WT Lister 427 MiTat1.2 *T. brucei* reported previously^26,27^. Moreover, the number of *T. congolense* VSGs detected exceeds the VSG numbers detected in *T. brucei* after impairment of monoallelic expression following RNAi of VEX1 and/or VEX2^26,27^.

**Figure 3.**
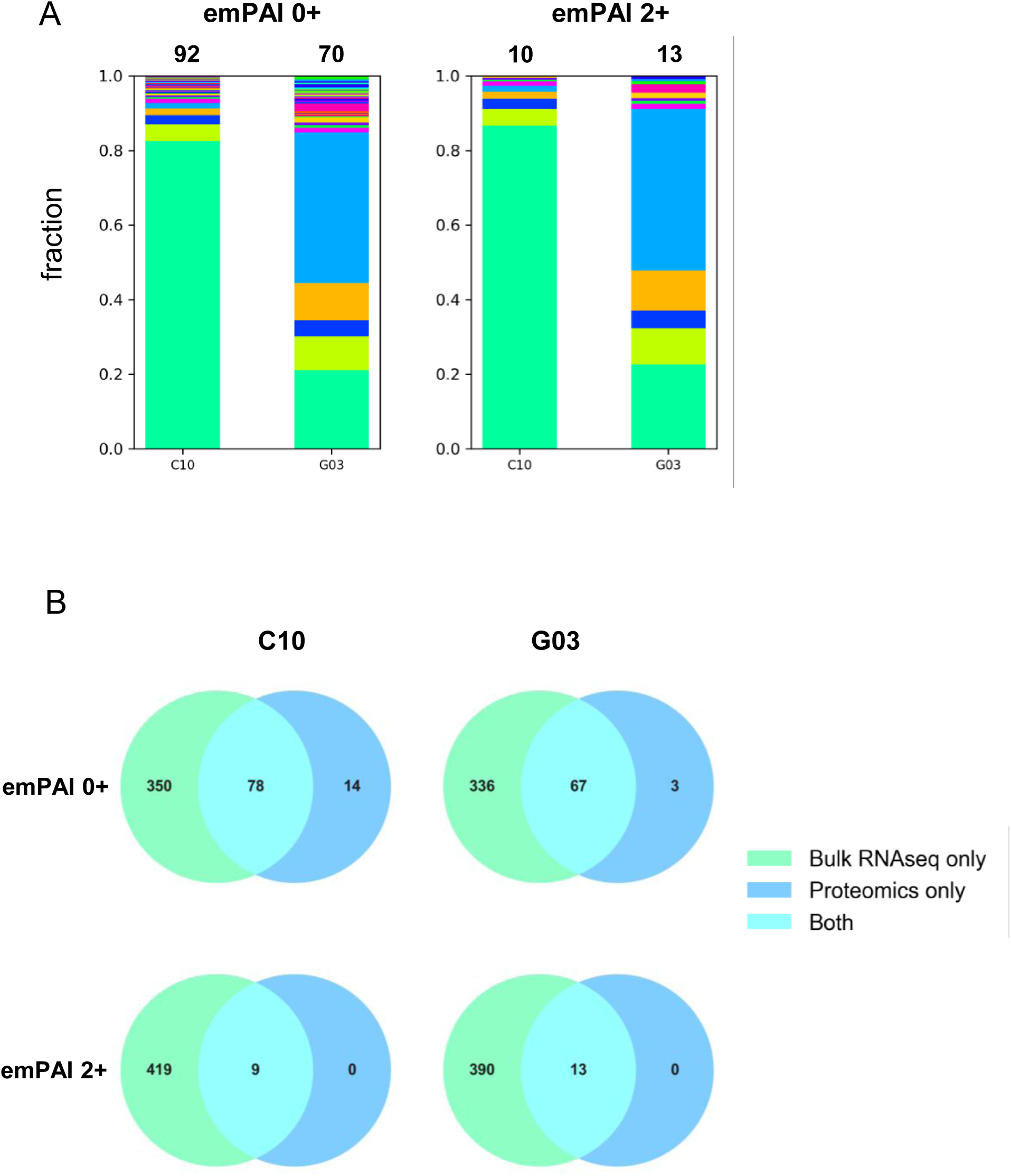
Diversity of VSG proteins expressed in cultured wild type *T. congolense*. Cell surface proteins were enriched and purified from ∼1x10^9^ bloodstream cells from two separate *T. congolense* WT populations (C10 and G03), and the relative proportion of identified VSG proteins in each is shown, with each colour representing a distinct VSG. Plots for two minimum inclusion emPAI thresholds are shown, and the total number of detected VSG proteins is shown above the plots.

### *T. congolense* does not employ monoallelic *VSG* expression *in vitro*

The diversity of *VSG* transcripts and VSG proteins seen in even small populations of *T. congolense* (Figs.2, 3) could be explained by rapid switching between single expressed *VSG*s in each cell, or expression of multiple *VSG*s simultaneously in a single cell. To distinguish between these possibilities, we performed single cell transcriptomics (scRNA-seq) on both *T. congolense* and *T. brucei* grown *in vitro*. For the latter, we chose to examine EATRO1125 AnTat 1.1 90:13 cells, a strain capable of differentiation from replicative slender bloodstream forms to cell cycle-arrested stumpy forms^59–61^, unlike *T. brucei* Lister 427, in which most scRNA-seq analysis of *VSG* expression control has been conducted^29,62,63^ and which does not differentiate naturally to short stumpy forms^64^.

Previous work, using flow cytometry, has shown that EATRO1125 AnTat 1.1 90:13 populations *in vitro* predominantly express a single VSG on their surface^65^, consistent with monoallelic expression, but *VSG* expression analysis by scRNA-seq has not been performed in the same conditions. *T. congolense* IL3000 bloodstream populations also undergo density-dependent cell cycle arrest, despite the absence of morphologically detectable stumpy forms^50,66^ and, unlike in *T. brucei*^65^, it is unclear if such differentiation might be connected to VSG switch rate or mechanism.

Single-cell RNA-seq was conducted for bloodstream forms of each trypanosome species growing *in vitro* using Chromium (10× Genomics) droplet-based single cell isolation and cDNA processing as previously described^59^. For both species, reads were mapped to long-read sequence assemblies of their corresponding genomes, which encompass chromosome telomeres (Krasilnikova et al, BioRXiv10.64898/2026.02.19.706783; Larcombe et al, BIORXIV/2026/741217). Because the focus of this analysis was *VSG*s being expressed in each cell, in addition to canonical quality control filters (see methods), cells with a low percentage of *VSG* transcripts were also removed (Fig.S6A,B), leading to the recovery of a total of 13,160 and 15,127 individual transcriptomes for *T. brucei* and *T. congolense*, respectively. A median of 1,057 and 606 genes was found per cell in *T. brucei* and *T. congolense*, respectively. The two datasets were then normalised, subjected to unsupervised cell clustering and visualised by dimensionality reduction (UMAP, Uniform Manifold Approximation and Projection), as described previously^67^. Six distinct clusters of related transcriptomes were predicted in *T. brucei*, whereas seven clusters were seen in *T. congolense* (Fig.4A, Fig.S7). These clusters are independent of *VSG* expression, since *VSG* genes were excluded from the variable gene list used for principal component and clustering analysis, allowing us to query *VSG* transcript levels across the population.

**Figure 4.**
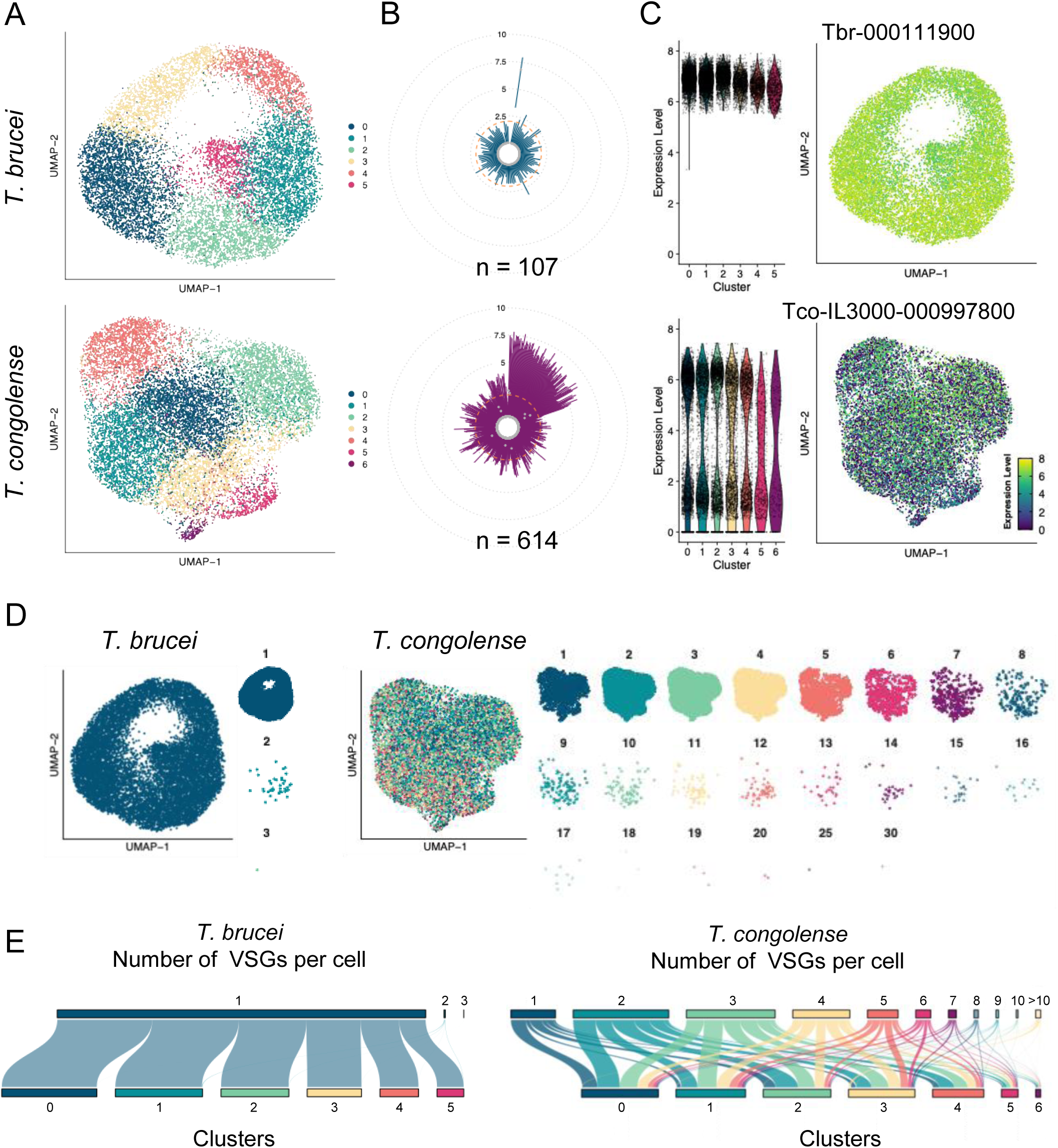
Most *T. congolense* cells do not employ monoallelic *VSG* expression *in vitro*. **A.** UMAP plots of *T. brucei* (top) and *T. congolense* (bottom), where each dot represents a cell coloured by cluster identity (6 clusters in *T. brucei*, 7 clusters in *T. congolense*). **B.** Radial violin plots showing the distribution of the expression of each detected VSG (individual violin plots) in the *T. brucei* (blue, top) and *T. congolense* (purple, bottom) samples. Because of the high number of genes depicted (107 for *T. brucei*, 614 for *T. congolense*), each violin plot appears as a line connecting the minimum and maximum values, with the median shown as a grey dot. The concentric dotted lines mark log2- normalised expression levels of 2.5, 5, 7.5 and 10, while the dashed orange line marks the threshold of 2 applied to count the number of *VSG*s expressed per cell. **C.** Violin and UMAP-feature plots of the Tbr-00111900/AnTat1.1 *VSG* in *T. brucei* (top) and Tco-IL3000-000997800 *VSG* in *T. congolense* (bottom). Each violin plot (left) is coloured by cluster identity, and each dot represents a cell. Each dot in the feature plots (right) represents a cell and is coloured by *VSG* expression level, as shown in the gradient legend. **D.** UMAP plots for either *T. brucei* or *T. congolense* where each dot is coloured by the number of *VSG*s expressed in the cell. Larger plots show all cells in each parasite sample; smaller plots show cells expressing the different numbers of *VSG*s. **E.** Alluvial plots for both *T. brucei* (left) and *T. congolense* (right) showing from which clusters cells expressing different numbers of *VSG*s originate.

To analyse *VSG* expression, we again used *T. congolense* genes identified in the assembled genome by their possession of Pfam domain PF13206 and annotation as a *VSG* by Companion; for *T. brucei*, *VSG* genes in the genome were identified as those that possessed the C-terminal Pfam domain PF10659 and either the VSG-a or VSG-b family Pfam domains PF00913 and PF13206, respectively. 107 distinct *VSG*s to which at least one transcript was mapped were detected in the *T. brucei* sample, while 614 distinct *VSG*s were similarly detected in the *T. congolense* sample (Fig.4B).

Expression of *VSG*s was very different in the populations of the two parasite species. In *T. brucei*, a single *VSG* (AnTat1.1, named Tbr-000111900 in the genome assembly) displayed high and uniform expression in all *T. brucei* clusters and in virtually all cells (Fig.4C; median expression 6.8 (range 5.3- 7.9), with lower expression (3.3) in just one cell). No other *T. brucei VSG* approached this level of expression (Fig.S8A), with only 11 *VSG*s detected at >2 expression in at least one cell (Fig.4B, Fig.S8A,B). In contrast, 96 *T. congolense VSG*s were expressed at a level of >5 in at least one cell in the population (Fig.4B), but with considerable variation in transcript levels between cells. This variation in expression is illustrated by examining Tc-IL3000-000997800 (Fig.4C), the *VSG* with expression >5 in the largest number of cells (5,937): expression was not uniform, with some cells in every cluster displaying either high transcript levels (>5) or substantially lower levels (∼1.5-2), and some cells not detectably expressing the *VSG* (Fig.4C). Many other *T. congolense VSG*s displayed maximum expression that compared with or even exceeded that of Tc-IL3000-000997800 (Fig.S8A), and many showed a similar variable pattern of expression in cells across the population (50 examples are shown in Fig.S8B). These data extend our understanding derived from population-level RNA-seq (Figs.1,2) and proteomics (Fig.3)^26,27^, in that whereas the very large majority of *T. brucei* cells display very abundant expression of a single *VSG* transcript, multiple distinct *VSG*s display variable levels of expression within the cells comprising a *T. congolense* population.

To determine whether or not the increased diversity of *VSG* expression in *T. congolense* is due to a higher rate of *VSG* switching than in *T. brucei*, we next used the barcodes that define each individual cell in the parasite populations to calculate how many *VSG*s were expressed in single cells. To this end, we set a relatively permissive minimal expression cut-off of 2, as this included the 11 *T. brucei VSG*s that show low but detectable expression (Fig.4B). In the *T. brucei* sample, 99.7% of cells expressed a single *VSG* (Figs.4D and 5A,B; Tbr-000111900/AnTat1.1). Only 41 *T. brucei* cells (∼0.3% of the total) showed evidence of expressing more than one *VSG*, with 40 cells expressing two *VSG*s and one cell expressing three (Fig.4D and Fig.5B,C). The huge majority of cells showing evidence for expression of just Tbr-000111900/AnTat1.1 is indicative of an overwhelming predominance of monoallelic *VSG* expression in EATRO1125 *T. brucei*, and consistent with *in vitro* flow cytometry^65^ and scRNA-seq of the same *T. brucei* strain *in vivo*^68^ and of cultured wild type Lister 427^29^. In contrast, only 12.6% of *T. congolense* cells showed evidence for expression of a single *VSG* transcript (Fig.4D, Fig.5A,B), with the large majority (87.4%) expressing at least two *VSG*s (Fig.4D, Fig.5A,B). Co-expression of two or three *VSG* transcripts per cell was most common (∼27% and 25%, respectively), but co-expression of 4-8 *VSG* transcripts was found in 13% of the population (Fig.S9A), and co-expression of up to 30 *VSG* transcripts was seen (Fig.4D, Fig.S9A). These data suggest that unlike in *T. brucei*, monoallelic *VSG* expression is rare in cultured *T. congolense*. Co-expression of *VSG*s in *T. congolense* was not due to distinct behaviour of any of the seven cell clusters we detected in the scRNA-seq data (Fig.4E) but is instead a general feature of most cells in the population.

**Figure 5.**
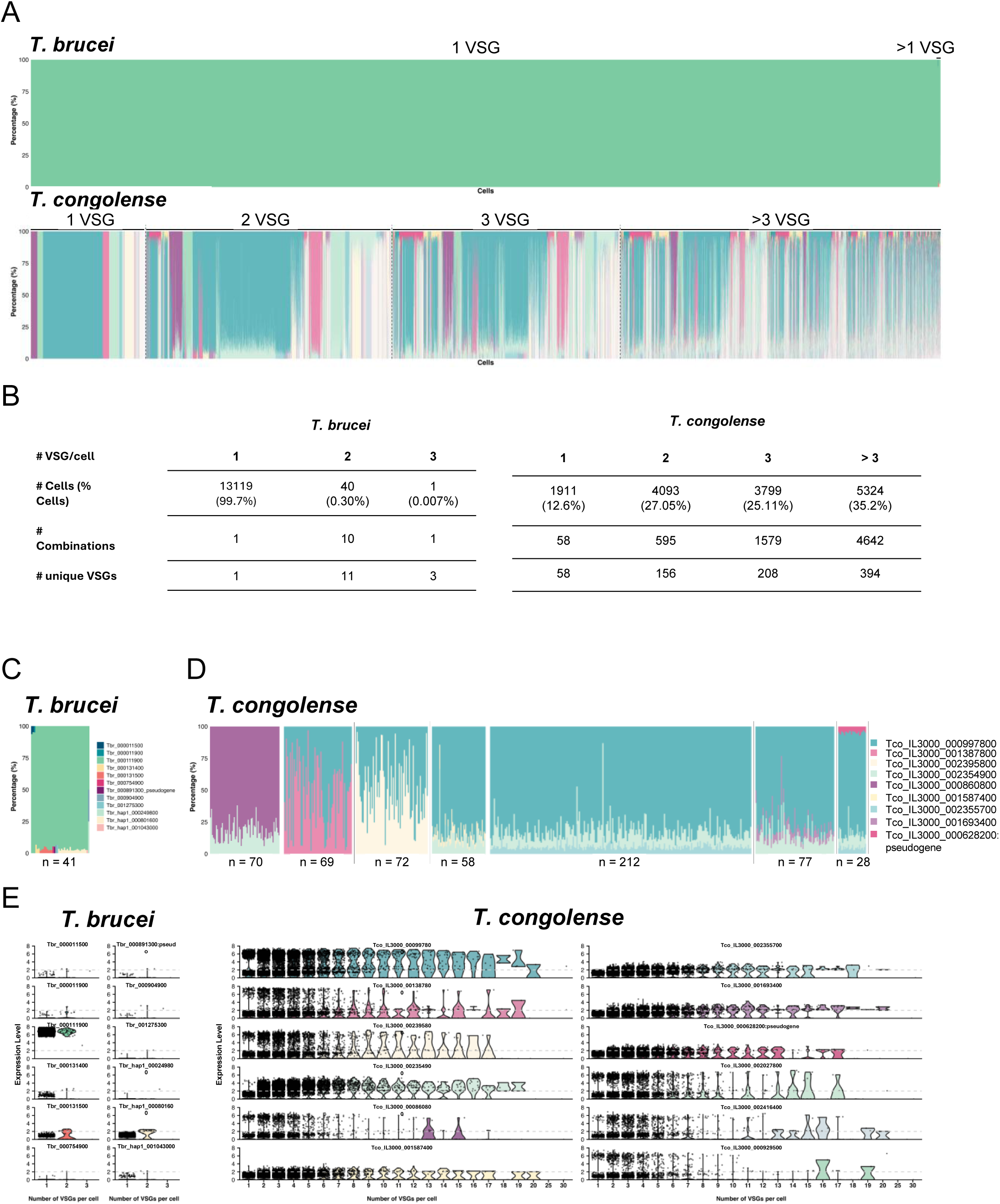
Dynamic co-expression of *VSG*s in *T. congolense* but not in *T. brucei in vitro.* **A.** Bar graphs (*T. brucei* top; *T. congolense* bottom) showing the percentage of each *VSG* detected in each cell (individual bar). Each *VSG* is coloured differently, and the cells are ordered by the number of *VSGs* detected per cell. **B.** Tables summarising results obtained for *T. brucei* (left) and *T. congolense* (right). **C.** Zoomed-in plot from (A), showing the 41 *T. brucei* cells expressing more than one *VSG*. **D.** Zoomed-in plots from (A), showing seven different combinations of *VSG*s detected per *T. congolense* cell. **E.** Violin plots showing the expression levels of the *VSG*s shown in (C) and (D) in cells expressing one or more *VSG*s; each dot represents one cell, and each violin plot is coloured by the corresponding *VSG*.

### *VSG* co-expression in *T. congolense* is dynamic

Population-level RNA-seq detected hundreds of *VSG* transcripts at varying levels within *T. congolense* WT and HR mutant populations (Figs. 1,2), whereas at most ∼30 *VSG*s were co-expressed in a single WT cell, with most cells co-expressing lower numbers (Fig.4). To begin to understand how these data might be resolved, we examined the combinations of co-expressed *VSG* transcripts in the scRNA-seq data (Fig.5). For *T. brucei*, in 13,119 cells (99.7% of the total) AnTat1.1 (Tbr-000111900) accounted for 100% of the detected *VSG* transcripts (Fig.5A,E). Examination of the 41 *T. brucei* cells that expressed more than one VSG revealed 11 other distinct *VSG*s expressed in 10 combinations when two *VSGs* were co-expressed in a cell, and one example of three distinct *VSG*s in a single cell (Fig.5B,C,E). Tbr-000111900/AnTat1.1 remained the predominant *VSG* in each cell that co-expressed two *VSG*s (∼93% of the reads, Fig.5C; >5 expression, Fig.5E). In contrast, in the single cell with three *VSG* transcripts, Tbr-000111900/AnTat1.1 expression was substantially lower (<4, Fig.5E; 50% of reads, Fig.5C), but still higher than the two other *VSG*s.

Similar analysis of *T. congolense* revealed a substantially more complex pattern of *VSG* co-expression (Fig.5). Unlike in *T. brucei*, 58 different *VSG*s were detected amongst the 1,911 *T. congolense* cells that expressed one *VSG* (Fig.5A,B). Examination of the *T. congolense* cells that co-expressed two or three *VSG*s revealed expression of 156 and 208 unique genes, respectively, in 595 and 1,579 combinations (Fig.5B), much surpassing the 11 individual *VSG*s and 10 combinations detected in *T. brucei* cells co-transcribing distinct *VSG*s (Fig.5B). Grouping all *T. congolense* cells in which >3 *VSG*s were co-expressed revealed 4,642 combinations of 394 individual genes, which appears comparable to the total number of *VSG*s detected in population-level RNA-seq (Figs.1,2). Thus, the wide range of *VSG*s that are flexibly co-expressed at a single cell level can explain the *VSG* diversity seen in population data.

To explore *T. congolense* VSG co-expression dynamics further, we focused on a small number of *T. congolense VSG* transcripts found co-expressed along with one (n = 3), two (n = 3) or three (n=1) other *VSG*s (Fig.5D). These combinations were chosen from the top 35 combinations of co-expressed *VSG*s found in the sample (Fig.S9B). When two *VSG* transcripts were co-expressed, there was considerable variation in the proportion of each transcript in individual cells. For example, expression of Tco-IL3000-000860800 ranged from 62-94% of transcripts when co-expressed with Tco-IL3000-002354900 (6-38%; Fig.5D), perhaps indicating preferential expression of the former. Conversely, when *VSG*s Tco-IL3000-000997800 and Tco-IL3000-001387800 were co-expressed, each ranged from ∼2.5-97.5% of total transcripts, indicating no preferential expression (as was also seen with Tco-IL3000-000997800/Tco-IL3000-002395800 co-expression; Fig.5D). Similarly, co- expression of three or four *VSG*s did not reveal uniform levels of each VSG transcript in every cell (Fig.5D). Thus, unlike in *T. brucei*, no single *T. congolense VSG* transcript predominates and, instead, these data suggest considerable dynamism in *VSG* co-expression.

Extending the analysis, using select *VSG*s to determine transcript abundance in cells expressing 2-30 *VSG*s (Fig.5E), suggested two things. First, expression of *VSG*s that were found to be relatively abundant when only small numbers of *VSG*s were co-expressed showed a notably decreased transcript level in cells expressing many *VSG*s, perhaps indicating a maximum total quantity of *VSG* transcript in any given cell. Second, some *VSG*s were expressed at low levels in all cells, suggesting that these *VSG*s may exhibit less of a contribution to *VSG* diversity through their reduced expression dynamism.

### *T. congolense* co-expressed *VSG*s are transcribed from across the genome

To explore how transcription of *VSG*s occurs in *T. congolense*, we next mapped the location of the genes detected by scRNA-seq to the genome. Fig.6A and B summarises the locations of all 399 *VSG*s that were recovered, while Fig.6C details the location of each expressed *VSG* in the 13 largest *T. congolense* chromosomes, as well as in one of two chromosomes that were intermediate in size and sequence content between the 13 largest and 147 minichromosomes found in the available genome assembly (Fig.S10; Krasilnikova et al, BioRXiv10.64898/2026.02.19.706783). The genes encoding the expressed *VSG*s were found on all classes of chromosomes, but locations differed. In the minichromosomes, all expressed *VSG*s corresponded to genes directly adjacent to the telomere tract (Fig.6B, Fig.S10). In contrast, only a minority of expressed *VSG*s were telomere-adjacent in the largest chromosomes (Fig.6B), with the majority at distal locations and often separated from the telomere by non-expressed *VSG*s (Fig.6C, Fig.S11A). Thus, unlike in *T. brucei*, *VSG* expression in *T. congolense* is not limited to transcription sites linked to telomeres, suggesting a lack of dedicated ESs. Consistent with this suggestion, evaluation of genes proximal to the transcribed *VSG*s in *T. congolense* (Fig.S11B) identified a range of predicted functions, not limited to the *ESAG*s that are co-transcribed with *VSG*s in *T. brucei* ESs^13^. One chromosome was notably unusual in this analysis: *VSG*s were found across the length of chromosome 4, with only those closest to the telomere detected by scRNA-seq (Fig.6C), or indeed by bulk RNA-seq (Krasilnikova et al, BioRXiv10.64898/2026.02.19.706783). This finding suggests that chromosome 4 houses a large transcriptionally silent pool of *VSG*s.

**Figure 6.**
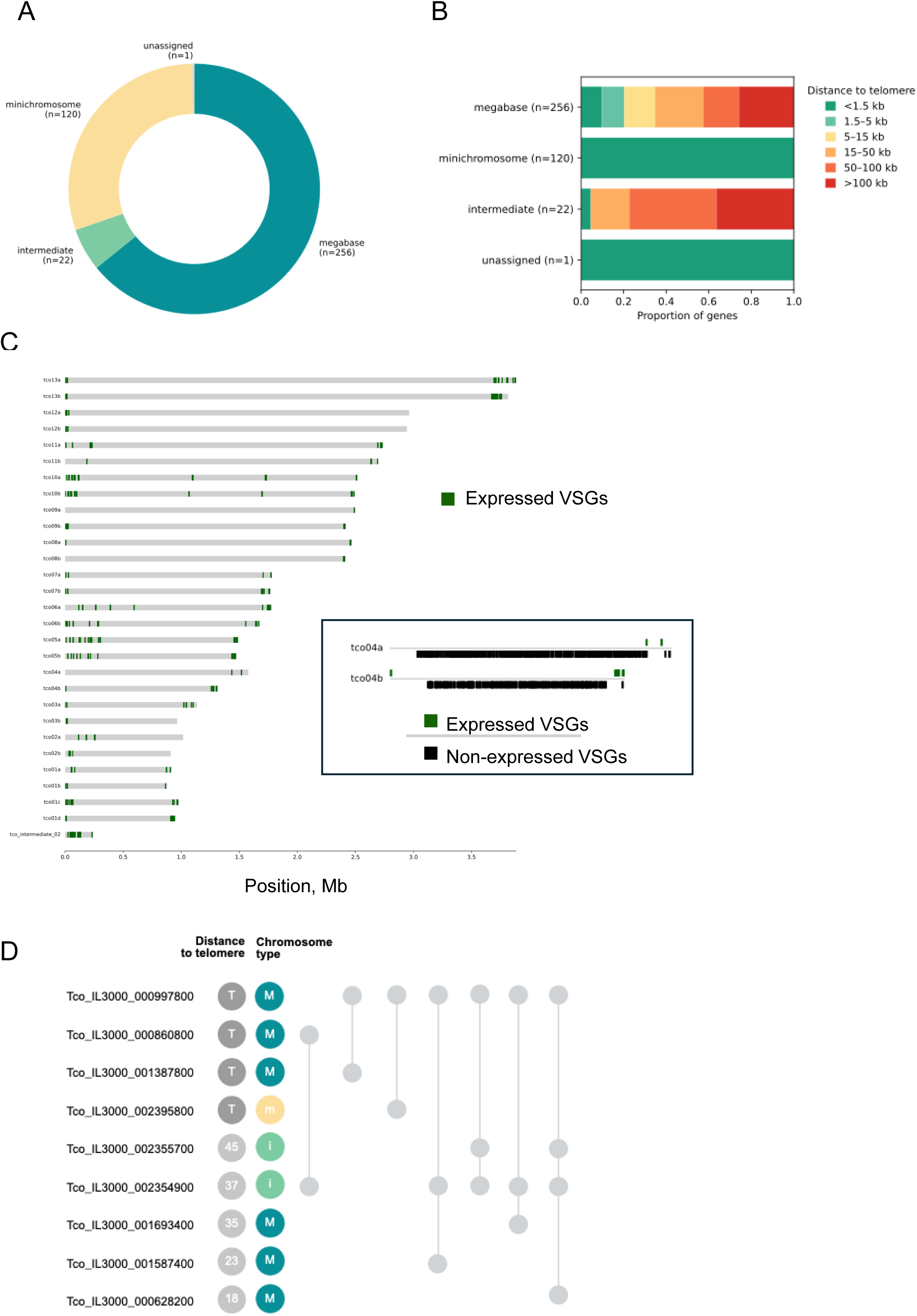
Co-expression of *VSG*s occurs across the *T. congolense* genome. **A.** A breakdown of which contigs/chromosomes contain the expressed *VSG*s detected in scRNA-seq data (expression level >2); ‘megabase’ refers to telomere-to-telomere chromosomes above 800 kb in length; intermediate, 200-500 kb; minichromosomes, <100 kb; ‘unassigned’ refers to a contig which has not been categorised. Number of *VSG*s found on each type of contig is indicated in brackets. **B.** The distance to nearest telomeric repeat (TTAGGG) of the expressed *VSG*s, separated by contig type. **C**. A diagram showing the genomic position of the expressed *VSG*s on the megabase and intermediate chromosomes. A separate, boxed diagram highlights the position of both expressed and non-expressed *VSG*s on chromosome 4 (tco04a and tco04b). **D**. Commonly co-expressed *VSG* combinations in a cell and their genomic localisation. The distance to the nearest telomere repeat is shown in kb (‘T’ indicates less than 1.5 kb from telomere?); ‘M’ refers to megabase chromosomes, ‘m’, minichromosomes, and ‘i’ intermediate chromosomes.

Next, we examined the locations of a selection of the *VSG*s found to be co-expressed by scRNA-seq (Fig.5). When co-expressed in either pairs or groups of three, there was no commonality of location in the genome: expression of *VSG*s was seen from each type of chromosome and regardless of whether the genes were telomere-adjacent or -distal (Fig.6D). These data indicate that the lack of monoallelic *VSG* expression in *T. congolense* reflects a lack of positional restraint on *VSG* transcription, revealing a profound difference in the operation of VSG expression relative to *T. brucei*.

### Absence of *T. congolense* monoallelic *VSG* expression *in vivo*

The above analyses were conducted in parasites grown in culture. To test if *T. congolense VSG* expression also occurs in the absence of monoallelic control in parasites *in vivo*, ∼1000 IL3000 cells were inoculated into mice, and parasites were harvested 5, 8 and 26 days later and subjected to scRNA-seq after purification from blood. Data from each timepoint were merged, revealing that the percentage of *VSG* transcripts detected per cell *in vivo* (∼2% of the transcriptome, Fig.S12A) was substantially lower than that seen in cultured IL3000 (∼10%; Fig.S12A). To understand if this was a consequence of the generation of scRNA-seq data from *in vivo* samples, we assessed the percentage of *VSG* transcripts in published population-level RNA-seq samples from *T. congolense* grown *in vitro*^35^ or *in vivo*^50^. Again, *VSG* transcripts represented a substantially lower and comparable level of the total transcriptome in parasites recovered from mice than in culture (Fig.7A). In *T. brucei*, it has been reported that *VSG* and ES transcription is reduced *in vivo* as parasite populations are enriched for non-replicating, transmission-ready short stumpy forms and denuded of replicating long slender forms^69^. To ask if this might explain the lowered detection of *T. congolense VSG*, we performed two analyses in *T. brucei*. First, we re-examined existing scRNA-seq data that followed EATRO1125 *T. brucei* cells as they undergo induced slender to stumpy differentiation *in vitro*^59^, revealing a substantially reduced percentage of *VSG* transcript in cells belonging to two stumpy populations compared with two slender populations (Fig.7B). Second, we compared the percentage of *VSG* transcript in scRNA-seq data of EATRO1125 *T. brucei* grown *in vitro* (this study) or after 7 days growth in mice^60^. Again, we found substantially lower levels of *VSG* transcript in parasite cells from mice (<1% of the transcriptome; Fig.S12B) than in culture (∼7%; Fig.S12B). As non-replicative *T. brucei* stumpy form cells dominate the infecting population in mice^60,70^, where a related differentiation process occurs in *T. congolense*^50,66,71^, it seems likely that lowered *VSG* expression is a common adaptation of both species in preparing for transmission to the tsetse.

**Figure 7.**
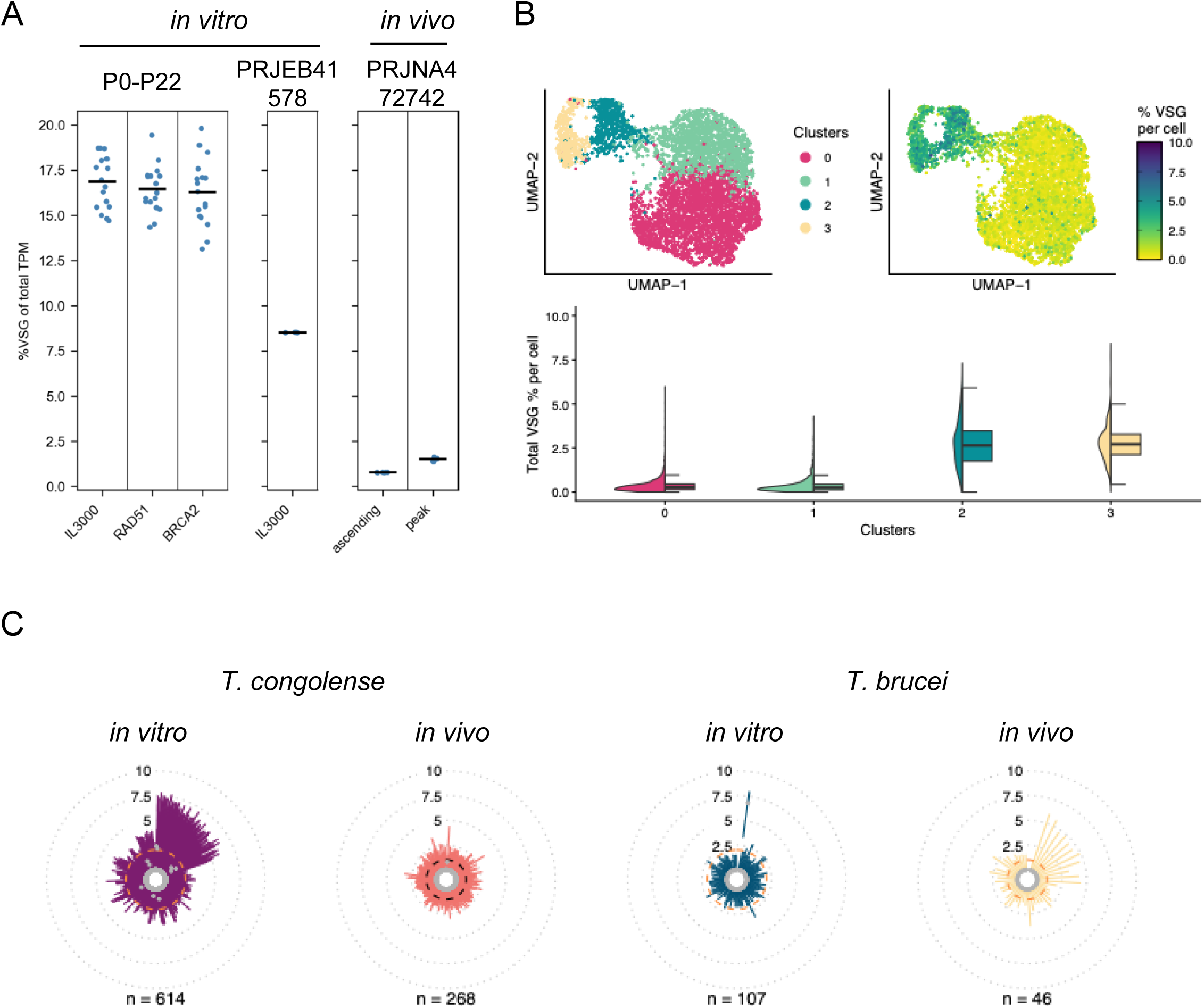
*VSG* expression differs in culture and in mice. **A)** Percentage of *VSG* transcripts (out of total transcripts per million – TPM) in *T. congolense* bulk RNA-seq datasets, both from this study (P0-P22 samples) or previously published studies (PRJEB411478 and PRJNA472742). **B)** Percentage of *VSG* transcripts in the published scRNA-seq data from Briggs *et al*., 2021^59^, where both long slender and short stumpy bloodstream forms of *T. brucei* were analysed. Top left panel, UMAP coloured by cluster (0 and 1, short stumpy forms; 2 and 3, long slender forms). Top right panel, UMAP coloured by percentage of *VSG* transcripts per cell. Bottom panel, violin plots showing the distribution of the percentage of *VSG* transcripts per cluster. **C)** Radial violin plots showing the distribution of the expression of each detected *VSG* (individual violin plots) in the *T. congolense in vitro* (purple, same as in Fig.5B) and *in vivo* (salmon), and in *T. brucei in vitro* (blue, same as in Fig.5B) and *in vivo* (yellow). Dashed lines represents the expression level cutoff used for calling a VSG as expressed: 2 for the *in vitro* samples, and 1 for the *in vivo* samples.

To further explore the implications of this change in *VSG* expression in *in vivo* samples we first examined *VSG* transcript abundance in individual *T. brucei* and *T. congolense* cells across the population (Fig.7C, Fig.S12A,B). In both species the number of *VSG*s detected at even one read/cell in the *in vivo* samples was approximately half that seen *in vitro* (Fig.7C). For *T. brucei*, the predominant and abundant expression of one *VSG* was no longer seen in the *in vivo* sample and instead several *VSG*s were detected, but each at much lower maximal expression levels relative to *in vitro* cells (Fig.7C, FigS12B), with such expression derived from only 1-2 *VSG* reads/cell (Fig.S12B). A similar reduction in *VSG* expression was seen in *T. congolense* taken from mice and, though again each *VSG* displayed lower maximal expression compared with *in vitro* cells (Fig.7C), a wider range of reads/cell (1-15) was seen in *in vivo*-derived *T. congolense* cells than in *T. brucei* (Fig.S12A,B). Given the much-reduced overall levels of detectable *VSG* expression in the *in vivo* scRNA-seq samples for both species, we next determined how many *VSG*s are expressed in individual cells (Fig.8) using an expression cutoff of 1 (Fig.7C, Fig.S12A,B). Strikingly, the same distinct pattern of *VSG* expression between *T. brucei* EATRO1125 and *T. congolense* IL3000 that we described *in vitro* (Fig.4D) was seen *in vivo* (Fig.8): of 45,412 *T. brucei* cells examined after isolation from mice, the vast majority expressed a single *VSG* and only 77 (0.17%) expressed >1 *VSG* (max 3 *VSGs*/cell); in contrast, amongst 26,887 *T. congolense* cells examined, the majority (15,454 cells, 57%) expressed >1 *VSG*, and up to 22 *VSG*s/cell were detected. This distinction between *T. congolense* and *T. brucei VSG* expression was also seen when applying an expression cutoff of 1.5 (Fig.S12C).

**Figure 8.**
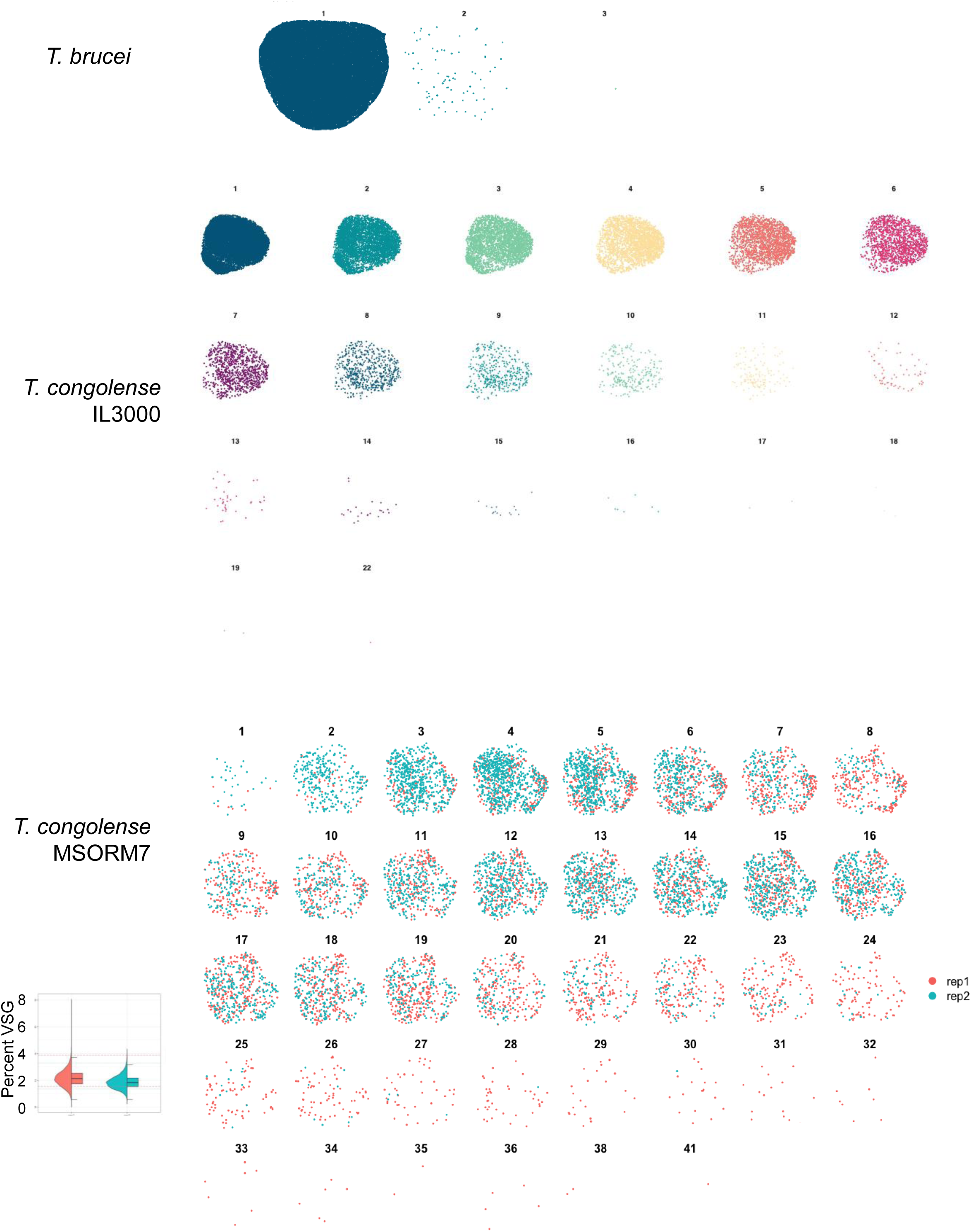
An absence of monoallelic *VSG* expression in *T. congolense* growing in mice. UMAP plots coloured by number of *VSG*s per cell. Top panel, *T. brucei* day 7 after initiation of infection from Larcombe *et al*., 2023^60^; middle panel, *T. congolense* IL3000 days 6, 8 and 26 (merged) after initiation of infection; and bottom panel, *T. congolense* MSORM7, where two repeats are shown 12 days after initiation of infection (the percentage of VSG transcripts per replicate is shown in the insert as violin plots).

The experiments above were conducted with a single strain of *T. congolense*. To test if the lack of monoallelic *VSG* expression we describe might be limited to that strain, we examined *VSG* expression in strain KTT/MSOROM7C1^72^, isolating cells from mice 12 days after they had been introduced by the bite of infected tsetse flies. Samples were purified from the blood of two mice and subjected to scRNA-seq, where *VSG* expression levels (∼2% of the transcriptome) compared with that seen in *T. conoglense* IL3000 cells from mice (Fig.8). Measuring the number of *VSG*s/cell in the two KTT/MSOROM7C1 samples revealed striking concordance with *VSG* expression in IL3000, with only a small minority (35 cells from 11,206; 0.3%) expressing a single *VSG*/cell and the large majority (99.7%) expressing >1 *VSG*/cell, with up to 41 *VSG*s/cell detected (Fig.8). Thus, a lack of monoallelic *VSG* expression appears to be a core feature of *T. congolense* biology.

## Discussion

Decades of research have demonstrated that expression of VSG in *T. brucei* occurs by allelic exclusion^3,25^, with recent work revealing the role of VEX and more recently described ESB-associated proteins in ensuring monoallelic transcription of just one of 15 dedicated telomeric *VSG* expression sites at any one time in a bloodstream form cell^2,29,31–33^. Here, we demonstrate that in the closely related parasite, *T. congolense*, *VSG* expression in bloodstream forms of the parasite is not limited to a single *VSG* per cell. Instead, multiple *VSG*s are co-expressed, with no evidence of expression being from loci related to *T. brucei VSG* ESs, or indeed for any dedicated expression locus. Thus, this work reveals the evolution of very distinct strategies directing the expression of the same surface antigen used for immune evasion in two closely related *Trypanosoma* species.

The single-cell transcriptome analysis we show here for *T. brucei* EATRO1125 is consistent with all related previous reports, *in vitro*^29,62,73^ and *in vivo*^68^ and examining either EATRO1125 or Lister 427 strains, showing that just one *VSG* ES is transcribed in a *T. brucei* cell at one time, unless VEX complex^26,28,29^ or further ESB protein functions^32,33^ are impaired. Both population- and single-cell analyses show how different VSG expression dynamics are in *T. congolense*. Even in very recently cloned *T. congolense* populations, it was not possible to detect a predominantly expressed *VSG*; instead, hundreds of *VSG*s were seen, whose expression pattern is not fixed but varies from clone to clone and over time in culture. This patterning suggests dynamic *T. congolense VSG* expression in the absence of immune pressure. scRNA-seq confirms this dynamism: when comparing thousands of single cells, *in vitro* and *in vivo*, in only a minority is just one *VSG* transcript detected; in most cells, a range of numbers of *VSGs* are co-expressed in thousands of combinations. Mapping the locations of co-expressed *VSG*s across the *T. congolense* genome reinforces the profound difference in *VSG* transcription control relative to *T. brucei*, with *T. congolense VSG*s being expressed from multiple loci across the genome and not limited to telomere-adjacent ESs. These data have several implications for the operation of antigenic variation in these two parasite species.

The simplest explanation for co-expression of multiple *VSG* transcripts in a *T. congolense* cell is that such genes are simultaneously transcribed, and hence there is no monoallelic transcription control of *VSG* expression, unlike in *T. brucei*. It should be noted, however, that this does not preclude the possibility of monoallelic expression of VSG on the cell surface. For instance, it is conceivable that while multiple *VSG* genes are co-transcribed, a downstream selection could operate, such as only one *VSG* mRNA being translated. If so, this would mean that cells of both parasite species display a homogenous VSG coat. To date, we have only performed proteomic analysis of VSG expression on populations of *T. congolense* cells, where it is notable that a restricted number of VSG proteins are seen relative to *VSG* transcripts, perhaps consistent with such control. However, this difference in transcriptomic and proteomic detection of VSG expression may reflect the greater sensitivity of the former approach. Nonetheless, the scRNA-seq data we provide reveal a fundamental mechanistic divergence between *T. brucei* and *T. congolense* in the point of VSG expression control, which suggests so far unexplored differences in the machinery that directs and controls *VSG* transcription between *T. brucei* and *T. congolense*. In this regard, no work has explored the function of VEX or other ESB-associated factors in *T. congolense*, or indeed in any trypanosomatid beyond *T. brucei,* despite the presence of at least some of these genes across this grouping of parasites^25^. A further feature acts to modulate *T. brucei* VSG expression: a highly conserved 16mer in the 3’ UTR of all *VSG* transcripts. Though the 16mer ensures *VSG* mRNA stability^74,75^ through recruitment of an RNA binding protein^76^ and modification of the polyA tail^77^, recent work has suggested it also acts to silence the transcribed *VSG* during switching to a new VSG^78^ and in monoallelic expression^79^. Intriguingly, the same 16mer is not found in *T. congolense VSG* genes, perhaps reflecting their freedom to be co-transcribed.

A more far-reaching potential implication of this work is that whereas each *T. brucei* cell expresses a homogenous VSG coat, *T. congolense* cells may express a coat comprising a heterogenous mix of VSG proteins. Though such a suggestion means a coat protein composition that would be fundamentally different between the two trypanosomes, it may nonetheless serve the same ultimate purpose: expressed VSG diversity. It is increasingly clear that *T. brucei* infections are marked by considerable population diversity in *VSG* transcripts detected at any given time^18–20,80^, and indeed this diversity seems to reflect tissue residence and immune clearance^68^. More limited *in vivo* analysis has been conducted in *T. congolense*, but bloodstream populations express a range of *VSG* phylotypes^51^. Is it possible, then, that the two parasites have evolved different routes to generate VSG coat diversity: in *T. brucei* rapid recombination of *VSG*s into the single active ES, and in *T. congolense* simultaneous expression of distinct VSGs? One reason this may not be the case is that the engineered loss of monoallelic VSG expression in *T. brucei* results in more rapid immune clearance from mice^81^. It will be important to examine patterns of *T. congolense* VSG expression during chronic infections^82,83^, which will test if population-level and single cell-level coat diversity can provide the same function in immune evasion.

This study also suggests a difference in the balance between recombination- and transcription-based *VSG* switching between the two *Trypanosoma* species. Variation in expression of *VSG*s between *T. congolense* cloned populations and individual cells is likely to reflect the expression of only a selection of the available *VSG* repertoire and a capacity for switching between those *VSG*s being expressed at any given time, reflecting a presumed need to change exposed VSGs as anti-VSG immunity arises in infections. RAD51- and BRCA2-dependent HR is an important driver of VSG switching and diversification in *T. brucei*^14–16,84^, but the lack of clear alteration to *VSG* transcript diversity in *T. congolense* after mutation of *RAD51* or *BRCA2* indicates that VSG switching is not mainly driven by HR, as has recently been argued in a further *Trypanosoma* species, *T. vivax*^85^.

Though it is conceivable that non-HR recombination might be used, a simpler explanation for the dynamic *VSG* transcript levels we describe in *T. congolense* single cells is a capacity to activate and deactivate expression of VSGs *in situ*. If so, further study is warranted, since *VSG*s are expressed not merely from telomeres, but more distally and when surrounded by non-expressed *VSG*s. These data indicate that there are no dedicated *T. brucei*-like telomeric *VSG* ESs in *T. congolense*^43^, consistent with an inability to detect the characteristic DNA repeats (50 bp^86^ and 70 bp^87,88^) associated with *T. brucei* ESs (Krasilnikova et al, BioRXiv10.64898/2026.02.19.706783). However, how such *in situ* transcriptional switching in *T. congolense* might operate is unclear: if *VSG*s are found within multigene transcription units, distinct expression of adjacent *VSG*s is perplexing, since the most extensive evidence of transcriptional control in kinetoplastids to date is the RNA Pol I-transcribed *T. brucei* VSG ESs^25^. Recent work has, in fact, revealed RNA Pol II transcriptional control at a gene cluster in *T. brucei*^89^, but no work has detailed the machinery that guides *VSG* transcription in *T. congolense*, and therefore so-far unrecognised non-telomeric *T. congolense VSG* ESs, containing undetected and controllable promoters upstream of individual *VSG*s or groups of *VSG*s, may be present. In this regard, perhaps the simpler *VSG* ESs used in *T. brucei* metacyclic cells found in the tsetse provide a blueprint^90^, given that *T. congolense* shares with *T. brucei* the expression of a restricted set of VSGs in this life cycle stage^39,91,92^. However, the location(s) where expression of *T. congolense* metacyclic *VSG*s occurs is unknown.

The difference in location and organisation of putative bloodstream *VSG* transcription control between *T. brucei* and *T. congolense* raises an evolutionary question: are gene-rich telomeric *VSG* ESs an innovation that arose only in *T. brucei*, or were they present in an ancestor but discarded in *T. congolense*? This distinction could reflect VSG coat differences between the species: whereas a homogeneous VSG coat in *T. brucei* would require transcription of a single *VSG* gene from a strong promoter, a heterogeneous VSG coat of the same density on *T. congolense* could arise from co- transcription of multiple *VSG*s from weaker promoters. Answering these questions will require experiments that rule out the suggestion (above) that monoallelic control of VSG expression does indeed occur in *T. congolense*, but by post-transcriptional, translational or post-translational means. Our data do, however, reveal a commonality in *VSG* expression between *T. brucei* and *T. congolense*, in that in both species *VSG*s represent a reduced fraction of the cell’s transcriptome *in vivo* compared with *in vitro*. Analysis of scRNA-seq data^59^ showed that non-replicating stumpy *T. brucei* cells express lower levels of *VSG* transcripts than replicating slender cells, consistent with previous suggestions that the process of shutting down transcription from the single active *VSG* ES begins during differentiation of *T. brucei* to tsetse transmission form cells^69^. No scRNA-seq data is currently available to distinguish replicating and non-replicating bloodstream form *T. congolense* cells, but such differentiation occurs *in vivo*^50,66,71^. Thus, the lowered levels of *VSG* transcripts in *in vivo* samples of *T. congolense*, where monoallelic expression remain absent, is consistent with similar *VSG* expression downregulation. However, as we do not yet know how *T. congolense VSGs* are transcribed, it is unclear if a similar or distinct *VSG* control mechanism operates in the two trypanosome species.

Though our data suggest a preference for transcriptional switching in *T. congolense*, the identification of a single *T. congolense* chromosome that is mainly composed of silent *VSG*s, accounting for ∼40% of the total *VSG* repertoire (Krasilnikova et al, BioRXiv10.64898/2026.02.19.706783), suggests recombination may yet play a role. In the scRNA-seq data provided here, there is no evidence for these *VSG*s being expressed, but it is possible that this chromosome provides a silent archive that contributes to VSG expression and immune evasion *in vivo*. Such putative *VSG* recombination may yet reveal a role for RAD51 and/or BRCA2 in *T. congolense* VSG switching. Nonetheless, the greater putative prominence of *VSG* transcription switching may be consistent with reduced levels of *VSG* pseudogenes in the *T. congolense* genome^36^ and may suggest a reduced capacity to generate mosaic VSGs, which are seen as important for long-term survival of *T brucei*^18,93,94^. How these features of *T. congolense* VSG expression and switching impact on transmission remains to be seen.

An assumption in the above discussion is that *T. congolense* VSG expression dynamics reflect a role in antigenic variation, which has been inferred to be a common mechanism of immune evasion across African trypanosomes^95^, given their shared possession of *VSG*s^36^. The lack of monoallelic *VSG* transcription need not rule this out, since recent work has documented conditions in which *P. falciparum* displays co-transcription of *var* genes^96^. Moreover, in *T. brucei* the establishment of monoallelic *VSG* expression from a single metacyclic *VSG* ES in tsetse salivary glands follows the simultaneous transcription of many *VSG* ESs^97^. Thus, the lack of monoallelic antigen transcription in *T. congolense* may represent one end of a spectrum of expression control during antigenic variation. Nonetheless, the pattern and dynamics of *T. congolense VSG* transcription we have described also appear reminiscent of the co-expression of surface proteins in many pathogen cells, where an explicit use of antigenic variation has not been demonstrated. In *Babesia*^98^, for instance, a distinction has been drawn between variant VESA1 proteins, which are expressed monoallelically and undergo switches by recombination, and further variable antigens encoded in the genome that are co-expressed and change less frequently during an infection. In *P. falciparum*^99^ and *Pneumocystis*^100^, several proteins are found to be expressed in addition to canonical variant antigens and are similarly exposed to immunity, but differ in not being expressed monoallelically. Closer to *T. congolense*, the genome of *Trypanosoma cruzi* contains an enormous repertoire of surface antigen genes, with recent analyses revealing heterogenous transcription in the mammal^101,102^, whose purpose remains a subject of debate^103^. Similarly, the extracellular trypanosome *T. theileri,* which infects *Bovinae*, encodes a range of multigene families that transcriptome analysis predicts to encode a heterogenous surface coat^104^. In this context, it is reasonable to ask if the absence of monoallelic *VSG* transcription in *T. congolense* means the parasite no longer deploys VSG on the cell surface for antigenic variation, but for a distinct form of immune evasion.

## Methods

### Parasites and mutants

For all *in vitro* experiments, bloodstream form *T. congolense* IL3000 was used (strain originally isolated at the International Livestock Research Institute, Nairobi, Kenya; a gift of Theo Baltz, University of Bordeaux). Cells were cultured at 34 °C with 5% CO_2_ in modified HMI-93 media^105^, with 10% goat serum (Gibco) and 20 g/L albumax II (Thermo-Fisher) replacing the serum/serum plus of the original. Cells were routinely cultivated in 6 or 24 well plates; adherent cells were flushed from the bottom of the wells using a P1000 pipette tip or 5 ml stripette, as appropriate, prior to passage into a new well, or for haemocytometer counting of cells. For large volumes, horizontal 75 cm^2^ flasks were used, with 10 ml or 25 ml stripettes used to flush the adherent parasites from the bottom surface prior to use. The generation of TcoRAD51^-/-^ H2B2 and TcoBRA2^-/-^ Fx2 clonal lines and the subsequent cloning and serial passaging of TcoIL3000 WT and the two homologous recombination mutant lines is described in (Krasilnikova et al, BioRXiv10.64898/2026.02.19.706783), as is the extraction and sequencing of population-level RNAseq data. *T. brucei* TbCas9 Lister 427^14^ or *T. brucei* EATRO 1.1 90:13 (a gift from Keith Matthews) parasites were also used, as indicated. Both were cultured in HMI-11 media containing 10% FBS at 37 °C and 5% CO_2_, with EATRO 1.1 90:13 cells not permitted to grow above a density of 1 x 10^6^/ml. *T. congolense* strain KTT/MSOROM7C1 was used for mice infections, and the derivation of this clonal strain has been described previously by Masuma et al^106^ and Tihon et al^72^.

### Genomes used for analysis

For *T. congolense*, a haplotype-resolved IL3000 *T. congolense* genome (Krasilnikova et al, BioRXiv10.64898/2026.02.19.706783) was used for all analyses. For *T. brucei* SLbarseq, *T. brucei* Lister 427 2018 genome build 68 was used from TriTrypDB (tritrypdb.org). For *T. brucei* single scRNA-seq, a *de novo* assembled *T. brucei* EATRO1125 (AnTat 1.1) genome was used, which will be described more fully elsewhere but, briefly, verkko (v2.2.1) was used to assemble the genome using HIFI, Nanopore and HI-C data (quality control was performed with QUAST (v5.3.0) and BUSCO (v5.7.0_cv1).

### Population-level RNA-seq

Short-read Illumina bulk RNA-seq data from prolonged *in vitro* growth (WT, RAD51-/-, BRCA2-/- samples) was trimmed using trim galore, and Salmon^47^ was used to quantify all gene and pseudogene transcripts annotated by Companion^46^ in the *T. congolense* IL3000 haplotype-resolved genome assembly (Krasilnikova et al, BioRXiv10.64898/2026.02.19.706783). The same approach was used to analyse the previously published *T. congolense* IL3000 RNA-seq data (accession number PRJEB41578 samples ERR4881956-ERR4881959, and accession number PRJNA472742 samples SRR7207631-SRR7207642)^35^. Separately, the experimental bulk RNA-seq data was also processed using vsgseq2 (https://github.com/goldrieve/vsgseq2)^48^. A *T. congolense VSG* database was used, based on putative *VSG* sequences from the haplotype-resolved *T. congolense* IL3000 genome. To identify which vsgseq2-assembled *VSG* transcripts correspond with which *VSG* genes in the haplotype-resolved IL3000 genome assembly, a blastn search was performed to identify matches to the annotated genes and pseudogenes at 100% identity and minimum query coverage of 95%.

## SL-barseq

For SL-barseq, the protocol from Touray et al^52^ was adapted to *T. congolense*. Firstly, both starting clones (P0) and P22 cultures from the RNA-seq experiment were recovered from stabilates, regrown for one passage and diluted to 0.5 cells/well in up to five 48 well plates for each of the six lines: WT IL3000 clones 1 and 2; TcoRAD51^-/-^ clones 2 and 5; and TcoBRCA2^-/-^ clones 1 and 2. When clones grew on these plates, they were transferred to 24 well plates, to a volume of 2 ml in modified HMI- 93, and RNA extracted using the 96 Well Plate Bacterial Total RNA Miniprep Super Kit, following the manufacturer’s instructions (BioBasic, USA BS585) once the adherent layer of parasites was almost confluent. If required, due to growth differences, pelleted cells were lysed in the collection plate with Rlysis-BG buffer from the kit and frozen at -80 °C, then the rest of the protocol followed after thawing. After harvest, 1 ml of new media was added to each well and, the following day, single stabilates were made of each clone by freezing 750 µl of cells combined with 750 µl of a mixture of 70% modified HMI-93:30% glycerol at -80 °C. *T. brucei* TbCas9 control clones were generated by dilution of a culture to 0.2 cells/well in a 96-well plate. Resulting clones were transferred to a 12 well plate and grown to around 1x10^6^/ml in 3 ml HMI-11; these were pelleted at 1000 *g* for 10 minutes and RNA made using the 96 Well Plate Bacterial Total RNA Miniprep Super Kit as above. DNA contamination was removed using RQ1 DNase treatment (Promega, M6101), following manufacturer’s protocol. cDNA was made from each clone using Superscript IV (Thermo Fisher, 18090200), following the manufacturer’s protocol and using a tag-anchored oligo-dT primer and 8 µl of DNase-treated RNA, in 96 well plates. Residual RNA was removed by adding a mix of 1U Ribonuclease H (Thermo Fisher, 18021071) and 7U Ribonuclease A (Qiagen, 19101), incubated for 30 mins at 37 °C. The cDNA was cleaned using Ampure RNA Clean XP beads (Beckman Coulter), with elution of the cDNA in 32 µl nuclease-free water. SLseq PCR was completed using this cDNA, in 96 well plates, with Phusion Polymerase (Thermo Fisher, F530L), the ONT adapted-forward anchored oligo-dT primer (JM125) for all, and the barcoded, ONT adapted-reverse SLprimer for each individual well (as listed in Supplementary Table 1). 30 PCR cycles were completed with a 50 °C annealing temperature and an extension time of 1 min 15s. The resulting PCR products were pooled in groups of up to 22 clones; Ampure XP bead-purified and barcoded libraries were then made from each pool, using the ONT SQK-LSK114 ligation sequencing kit with EXP_PBC001 PCR Barcoding Expansion kit, following the manufacturer’s instructions. The libraries were sequenced on a GridION Mk1 device using R10.4.1 MinION flow cells.

For long-read ONT SL-barseq data, an adapted version of the published VSG-Bar-seq processing and quantification script (https://github.com/cestari-lab/VSG-Bar-seq)^52^ was used, with modifications focused on adaptation for use with multiplexed ONT sequencing, genome annotation-specific features, as well producing summary output (scripts available on Zenodo). Briefly, the bash script split base-called fastq files by sample using the custom barcodes (or barcodes and ONT barcodes for multiplexed sequencing), mapped each sample separately to the genome using minimap2, filtered alignments by minimum mapping quality 1, sorted and indexed bam files using samtools, counted reads per gene using featureCounts (-L -M --fraction --nonSplitOnly) and, finally, combined the output across all samples into a single matrix of counts. Only samples that contained a minimum total of 1000 *VSG* reads/sample were retained for downstream assessment.

### Proteomics

Two clones from the SL-barseq experiment were chosen for proteomic analysis, based on a prediction of having several expressed VSG transcripts above 10% of total, and the predicted peptides of the resulting proteins being distinct enough to be distinguished by mass spectrometry. The selected single stabilates made after harvesting of the SL-barseq experiment were regrown in a 6-well plate to a density of around 1 x10^6^/ml, then re-stabilated. One stabilate per line was thawed into 200 ml of modified HMI-93 and the parasites grown to almost confluence in four horizontal 75 cm flasks per clone. For clone C10 ∼1 x10^9^ total cells were isolated; for clone G03 ∼9.2 x 10^8^ cells total. For both, a cell scraper was used to gently remove the adherent cells from the flask bottoms.

A soluble VSG-release protocol was followed, based on the full protocol in Cross (1984)^56^. Briefly, cells were washed twice in PSG buffer (in 1L water: 8.088g Na_2_HPO_4_, 0.468 g NaH_2_PO_4_.2H_2_O, 2.55 g NaCl, 10g glucose), resuspended in 1 ml 0.1 mM TLCK (Merck, T7254) in H_2_O, and incubated on ice for 5 min. This was then spun at 3000 *g* for 10 min at 4 °C and supernatant removed (=lysis supernatant). The pellets were resuspended in 1 ml 10 mM Na_3_PO_4_ pH8 + 0.1 mM TLCK and incubated at 37 °C for 5 min before chilling on ice, then centrifuged at 12000 *g* for 10 min at 4°C. Removal of the supernatant gave the ‘sVSG’ fraction; the resuspended pellet (in 1 ml 10 mM Na_3_PO_4_) provided the membrane pellet fraction. A 20 ul sample of each of the three fractions per clone was run on an SDS-PAGE gel to assess purity (Thermo Fisher, NP0321BOX; Fig. S8), before the sVSG fraction was analysed by the Proteomics facility at MVLS Shared Research Facilities (Glasgow University).

Protein samples were processed by filter aided sample preparation (FASP), by loading onto a 30k filter, washing twice with 200 µl of 8 M urea in 0.1M Tris/HCl pH8.5 (UA buffer), incubating with 100 µl 0.05M iodoacetamide in UA buffer for 20 min, washing three times with 100 µl UA buffer, then three times with 100 µl 0.05 M NH_4_HCO_3_ in water, and finally trypsinised overnight at 37 °C, with 120 µl of 20 µg/ml trypsin in 0.05 M NH_4_HCO_3_ and an enzyme to protein ratio of 1:100. The samples were eluted with 50 µl 10% Acetonitrile in water, acidified with trifluoroacetic acid, then dried by vacuum centrifugation. Dry peptide residues were solubilised in 20 µl 5% (v/v) acetonitrile with 0.5% (v/v) formic acid using a nanoflow uHPLC system (Thermo Scientific RSLCnano); online detection of peptide ions was by electrospray ionisation mass spectrometry MS/MS with an Orbitrap Elite MS (Thermo Scientific). Ionisation of LC eluent was performed by interfacing the LC coupling device to a NanoMate Triversa (Advion Bioscience) with an electrospray voltage of 1.7 kV. An injection volume of 5 µl of reconstituted peptides were desalted and concentrated for 10 min on trap column (0.3 × 5 mm) using a flow rate of 25 µl / min with 1 % acetonitrile and 0.1 % formic acid. Peptide separation was completed using a Pepmap C18 reversed phase column (3 μm, 100 Å, 75 μm × 50 cm, Thermo Scientific). Samples were fractionated with solvent A (0.1% (v/v) formic acid in water) and B (0.08% (v/v) formic acid in 80% (v/v) acetonitrile in water). The peptide separation was performed at a fixed solvent flow rate of 0.3 μl / min, with a solvent gradient of 4 % B for 1.5 min, 4 to 60 % for 100.5 min, 60 to 99 % for 14 min, held at 99 % for 5 min. A further 9 minutes at initial conditions for column re- equilibration was used before the next injection. The Orbitrap EliteTM MS acquired full-scan spectra in the mass range of m/z 300–2000 *m/z* for a high-resolution precursor scan at 120 000 resolving power (RP, at 200 *m*/*z*), while simultaneously subjecting up to the top 15 precursors to collision- induced dissociation (35% NCE) in the linear ion trap using rapid scan mode. Singly charged ions were excluded from selection, while selected precursors were added to a dynamic exclusion list for 30 seconds. Protein identifications were assigned using the Mascot search engine (v2.6.2, Matrix Science), interrogating protein sequences using a de novo assemble of the *Trypanosoma congolense* IL3000 genome (Krasilnikova et al, BioRXiv10.64898/2026.02.19.706783). A mass tolerance of 10 ppm was allowed for the precursor and 0.6 Da for MS/MS matching. The list of detected proteins was filtered to exclude non-VSG proteins, based on PFAM domains.

### Single-cell RNA-seq sample preparation, library preparation and sequencing

For the *in vitro* samples, single-cell RNA-seq was conducted as described in Briggs *et al* ^67^ with minor modifications. Briefly, fresh parasite cultures were used: *T. congolense* IL3000 clone 1 passage 4 cells grown in 6 well plates, and *T. brucei* EATRO 1.1 90:13 in 25 cm flasks. 1 x 10 ^6^ parasites per species were transferred to 15 ml tubes and centrifuged at 800 *g* for 10 min at RT; these were washed with prewarmed media, centrifuged again at 4 °C, and the pellets resuspended in 1 ml of PSG buffer (see above) supplemented with 0.04% BSA by gentle pipetting with wide-bore tips. Cells were kept on ice from this point onwards and were centrifuged and resuspended carefully in PSG supplemented with 0.04% BSA twice more, giving a final density of ∼2000 cells/µl. Cells were strained through a 40 μm filter into a new 1.5 ml tube before centrifuging again at 800 *g* for 10 min at RT, with the supernatant removed with a pipette. Cells were re-suspended in 150 µl PSG supplemented with 0.04% BSA, counted and a predicted total of 16 000 cells transferred to new 1.5 ml tubes and stored on ice. In parallel, the same protocol was applied to two samples of *Leishmania major* strain Friedlin promastigote cells (grown in 25 cm^2^ sealed flasks at 27°C), with the exception that the centrifugation steps were performed at 400 *g*, and only 8000 cells were collected for the downstream steps. For the Chromium (10x Genomics) library preparation step, two samples were prepared: *T. congolense* 16000 cells plus 8000 *L. major* cells, and *T. brucei* 16000 cells plus 8000 cells of *L. major*. Each sample was loaded onto the Chromium Control, and library preparation was performed with the Chromium Single Cell 3′ chemistry v4 kit. Libraries were sequenced using an Illumina NextSeq 2000, generating 28 × 190 bp paired reads to a mean reads per cell depth of 41,367 (*T. congolense*) and 42,271 (*T. brucei*). Library preparation and sequencing were performed by the Molecular Analysis facility at MVLS Shared Research Facilities (Glasgow University).

For analysis of cells isolated from infected mice, published data was used for *T. brucei* EATRO 1.1 90:13 (day 7 minus sample – files d7mS1 and d7mS3)^60^, while two independent *T. congolense* strains were used. For *T. congolense* KTT/MSOROM7C1, bloodstream forms were collected from infected murine hosts 12 days after tsetse bite, corresponding to an estimated parasitaemia of approximately 10^7.2^ trypanosomes/mL, whereas *T. congolense* IL3000 cells were collected at three distinct days (6, 8 and 26) after infection by injection. For strain KTT/MSOROM7C1, two independent biological replicates were generated from separate infected mice. Cells were DEAE-cellulose purified from infected blood and processed independently for single-cell RNA sequencing. For Chromium (10x Genomics) library preparation, approximately 10,000 cells from each replicate or sample were loaded onto a Chromium Controller, and libraries were prepared using the Chromium Single Cell 3′ Gene Expression v4 kit according to the manufacturer’s instructions. KTT/MSOROM7C1 were sequenced on an Illumina NovaSeq platform, generating paired-end reads of 28 bp (Read 1) and 91 bp (Read 2), to mean sequencing depths of 89,049 and 43,796 reads per cell for replicates 1 and 2, respectively. Sequencing was performed by Novogene (Germany). IL3000 libraries were prepared and sequenced as described for the *in vitro* samples.

### Single-cell RNA-seq general data analysis

Analysis of the scRNA-seq data was performed as described in Briggs *et al* ^67^ with minor modifications. Akin to Briggs *et al* ^67^, the reference genomes for *T. congolense* IL3000, *T. brucei* EATRO1125, and *L. major* Friedlin were modified so that to the end of each annotated coding region in the .gtf file, 2,500 bp were added; if the annotation overlapped with neighbouring genomic features, these extra regions were only extended to the base before the next annotated feature. As both *Trypanosoma* genome assemblies used in scRNA-seq analysis are haplotype-resolved and therefore contain duplicate sequences, to avoid issues with multimapping, annotations only for a single set of chromosomes (‘haplotype’) for each genome was used as a core set of annotations (haplotype A for *T. congolense,* and haplotype1 for *T. brucei*). Next, any putative *VSG* genes on the other haplotype contigs that were not present in the core set (based on all-vs-all blastp search of translated gene and pseudogene annotations, using the following criteria for ‘identical genes’: ≥95% identity, allowing for up to 1% difference in length), were added to the annotation list, as these are likely distinct *VSG* sequences and should be considered for mapping. This collapsed set of annotations was used for all scRNAseq analysis. Two genome files were then compiled using 10x Genomics Cell Ranger (version cellranger-9.0.1) mkref function (default settings): one containing the *T. congolense* and *L. major* genomes, and another containing the *T. brucei* and *L. major* genomes. Reads were then mapped to the corresponding genomes using Cell Ranger count function (default settings), generating a counts matrix for each. In the case of the *in vivo* samples, where *L. major* cells were not added, these were still aligned to the *Trypanosome/Leishmania* genome to allow for comparison with the *in vitro* data. The resulting Cell Ranger matrixes were imported into R (v4.4.1) using Seurat (v5.3.1), where all the analysis was then performed. As in Briggs *et al* ^67^, for each matrix a Seurat object was created using the min.cells = 5 and min.features = 3 settings, meaning genes were kept if detected in at least 5 cells, and only cells where 3 genes were detected were kept. In each sample, cells were then categorised as one of the three species or multiplets, based both on the classification given by Cell Ranger and the percentage of transcripts from each species (> 95% as *T. congolense*, > 95% as *T. brucei*, and > 90% *L. major* but < 5% *T. congolense/T. brucei* transcripts as *L. major* cell; the remaining cells were considered multiplets). *L. major-*labelled cells and multiplets were then excluded from the Seurat objects, leaving 19,889 *T. congolense* cells and 17,684 *T. brucei* cells from the *in vitro* samples, and 29,280 *T. congolense* IL3000 cells and 52,365 *T. brucei* cells from the *in vivo* samples for the downstream analysis. The cells were further filtered based on the following quality control (QC) parameters: number of transcripts per cell (> -1.5x Median Absolute deviation (MAD) and < 3.5x MAD), number of genes per cell (> -1.5x MAD and < 3.5x MAD), percentage of kDNA RNA (> -1.5x MAD and < 3.5x MAD) and percentage of rRNA RNA (> -1.5x MAD and < 3.5x MAD). In the case of the *in vitro* samples, a cutoff based on the percentage of *VSG* RNA was also used (> -1x MAD and < 3x MAD for *T. congolense* and > -1.5x MAD and < 3x MAD for *T. brucei*). Post-QC, 15,127 *T. congolense* cells and 13,160 *T. brucei* cells from the *in vitro* samples, and 26,887 *T. congolense* IL3000 cells and 45,412 *T. brucei* cells from the *in vivo* samples remained. The samples were then normalised and log2- transformed using Scran (v1.32.0)^59^. With *VSG*s excluded, a list of highly variable genes for each sample (950 genes for *T. congolense* and 1,387 *T. brucei* genes for the *in vitro* samples, 1,438 for *T. congolense* IL3000 and 1,062 for *T. brucei in vivo* samples) was obtained by intersecting the top 3000 variable genes resulting from two independent methods: Scran and Scater (uses log2-transformed counts) and Seurat (uses the raw counts). These were added to the Seurat objects and used for scaling (linear transformation) the data and running linear dimension reduction (PCA) using Seurat. The dimensionality of the Seurat objects was then assessed based on the obtained PCA scores, and the number of top principal components (PCs, dimensions) containing most of the data was inferred by analysis of the output from the ElbowPlot function (Seurat). Next, Seurat’s FindNeighbors and FindClusters were used, together with the visualisation package clustree (v0.5.1), to infer which resolution to use, in combination with the dimensions (PCs) defined above, for the non-linear dimensional reduction (UMAP) using Seurat. For the *in vitro* samples, cluster quality was checked by extracting the unique top 5 genes per cluster and plotting their average expression across clusters using the DotPlot and HeatMap functions from Seurat.

### Single-cell RNA-seq *VSG* data analysis

To assess *VSG* expression per cell, two layers of analysis were performed in R. First, we assessed the levels of expression for each *VSG* annotated in the genomes of *T. brucei* and *T. congolense*. To this end, the full lists of *VSG* genes for each species were imported and crossed-checked with the Seurat objects obtained in the previous section (note, even if only one transcript was detected in the sample for a *VSG* this was added to the number of *VSGs* detected in the sample). These were then plotted side-by-side by level of expression (log2-transformed, see previous section) as violin plots.

Each cell was then categorised by the number of *VSG* genes with an expression level > 2 (*in vitro* samples) or >1 (*in vivo* samples), and the expressed genes per cell were identified. Both sets of information were then added to the respective Seurat objects and used for multiple downstream data analyses (number of cells expressing each *VSG*; number of cells expressing multiple *VSG*s; number of *VSG* combinations in the sample; percentage of each expressed *VSG* per cell) and visualisation. Second, the percentage of each *VSG* in the cell was calculated. Only the expressed *VSG*s (expression level >2 *in vitro* and >1 *in vivo*, as above) were considered for calculation of the total percentage of *VSG* transcripts present in each individual cell; that being set as 100%, the percentage of each expressed VSG was then calculated and analysed.

Previous scRNA-seq analysis of VSG expression^29^ used the SoupX tool to predict and remove ambient RNA from the samples prior to the analysis described above. To test the impact of this tool, SoupX (v1.6.2) was used in all samples referred in this study with three rho values: 0.05, 0.1 and 0.15, predicting f5, 10 and 15% ambient RNA contamination. The results were relatively stable between the three values, and the overall conclusions of our study were not changed using SoupX (FigS13, compare with Fig.S12).

The following R packages (excluding the ones mentioned earlier) were used throughout the analyses: dplyr (v1.1.4), patchwork (v1.3.2), ggplot2 (v4.0.0), forcats (v1.0.1), stringr (v1.6.0), gridExtra (v2.3), viridis (v0.6.5), gghalves (v0.1.4), stats (v4.4.1), scater (v1.32.1), clustree (v0.5.1), BBmisc (v1.13), geneHapR (v1.2.4), tibble (v3.3.0), DoubletFinder (v2.0.6), tidytable (v0.11.2), celda (v 1.20.0), scuttle (v1.14.0), Matrix (v1.7.4), scCustomize (v3.2.2), UpSetR (v1.4.0), ggh4x (v0.3.1), ggforce (v0.5.0), sparceMatrixStats (v1.16.0), and data.table (v1.17.8).

### Gene content analysis surrounding expressed *VSG*s

Gene content surrounding the top expressed *T. congolense VSG* genes based on the scRNAseq data was assessed based on Pfam domain presence. Pfam_scan (https://github.com/aziele/pfam_scan) was used to detect Pfam domains across all annotated gene and pseudogene sequences in the *T. congolense* genome; this data was used to count the number of all detectable Pfam domains in the +-15 upstream and downstream genes of the *VSG*s of interest.

### Use of Artificial Intelligence

Throughout the scRNA-seq analysis, the Large Language Models (LLM) Gemini 1.5 Fast/Pro (Google) and Microsoft 365 Copilot (Microsoft), based on the GPT-5 reasoning model, were used to assist in troubleshooting code in R for both complex data analyses and plotting. The AI-suggested code was tested and validated by the authors. The LLMs were not used for the primary data analysis, plotting, or for interpretation of the results. Detailed AI-assisted contributions are documented within the source code made available by the authors.

## Supporting information

All supplementary Figures and a Supp Table, plus legends

## Data availability

Nanopore and Illumina reads for in vitro scRNA-seq and SLbar-seq analyses have been deposited in the NCBI Sequence Read Archive (SRA) under accession number PRJNA1435861; population-level in vitro RNA-seq data are available under accession number PRJNA1432479. ScRNA-seq *in vivo* data for KTT/MSOROM7C1 are available at the NCBI Gene Expression Omnibus (GEO), accession number GSE342023, while *in vivo* IL3000 scRNA-seq data are available at the NCBI SRA, accession number PRJNA1505143. Mass spectrometry proteomics data have been deposited in the ProteomeXchange Consortium via the PRIDE partner repository with the dataset identifier PXD075451 and 10.6019/PXD075451. Single-cell RNA-seq source code, .rds objects and accompanying tables are available in Zenodo: 10.5281/zenodo.21359916.

## Acknowledgments

We thank the Molecular Analysis and Proteomics Facilities of the MVLS Shared Research Facilities for their support and assistance with this work, and Monica Mugnier for comments on the manuscript. This work was supported by the Wellcome Trust (224501/Z/21/Z to RM, 221717/Z/20/Z to KM, 206815/Z/17/Z to KM, LM and RM), and the BBSRC (BB/W001101/1 to RM); the Roslin Institute (LM) is supported through core funding from the BBSRC (BS/E/D/20002173; BBS/E/RL/230002C).

## Author contributions

Experiments: JM, CM, SL, CL, KK, PM. Data analysis: JM, MK, CM, SL, PM, GO, EB. Project conceptualisation: JM, CM, SL, JVDA, LJM, KRM, RM. Manuscript writing and editing: JM, MK, CM, SL, EB, PM, JVDA, LJM, KRM, RM.

## Supplementary figure legends

**Supplementary Figure 1. Diversity of *VSG* transcripts expressed over time in culture by wild type *T. congolense* or null mutants of RAD51 or BRCA2.** Bulk RNA-seq data from WT, RAD51^-/-^ and BRCA2^-/-^ cells mapped to the *T. congolense* transcriptome, showing the relative proportion of all detected *VSG* transcripts per sample, with each colour representing a distinct *VSG* gene. P0, P11 and P22 refer to the passage number of the sample, in duplicate or triplicate. Data show an independent growth experiment, from independent starting clones, to those presented in Figure 1.

**Supplementary Figure 2. *VSG* transcript diversity in wild type *T. congolense* evaluated VSGseq2.** The *de novo VSG* transcript assembly and quantification pipeline vsgseq2 was used to analyse bulk RNA-seq data from WT *T. congolense* serial passage samples; the left-hand panels of the figure show the resulting assembled *VSG*s relative to their expression, with each colour/shade representing a distinct *VSG* gene. On the right-hand side, the analysis of the same data is shown by Salmon, using the *T. congolense* IL3000 transcriptome of the haplotype-resolved genome, for comparison (the same data are shown in Fig. 1 and Fig. S1). Colours of the transcripts detected by vsgseq2 that pass the following blastn search thresholds are matched to the *T. congolense* IL3000 transcriptome colours: 100% sequence identity, ≥95% query coverage. Those vsgseq2-detected and quantified *VSG* transcripts that did not pass the above thresholds are plotted in shades of grey.

**Supplementary Figure 3. Expressed VSG diversity in *T. congolense* persists regardless of defining criteria or inclusion TPM thresholds.** Bulk RNA-seq data (as analysed by salmon) of one replicate each of WT clone 1 P0, P11 and P22 samples, re-analysed using varying minimum TPM values (A) or *VSG* inclusion criteria (B). Numbers of *VSG*s detected is shown below each stacked bar.

**Supplementary Figure 4. Published IL3000 bulk RNA-seq data shows similar *VSG* diversity.** Previously published (ENA accession number PRJEB41578)^35^ *T. congolense* IL3000 bulk RNA-seq data is shown mapped to the *T. congolense* transcriptome, with sequencing run accession numbers for the samples shown below the plot. **A.** Panel shows the relative proportion of all detected *VSG*s per sample, with each colour representing a distinct *VSG* gene. **B.** Expression of all annotated genes and pseudogenes, with *VSG*s in red and non-*VSG*s in grey.

**Supplementary Figure 5. Enrichment of VSG from two clones of wild type *T. congolense*.** An SDS- PAGE gel of protein samples used for characterisation of VSGs expressed in two *T. congolense* WT clonal populations (C10, G03). The soluble VSG fractions (sVSG) were examined by mass spectrometry.

**Supplementary Figure 6. Quality controls for single cell transcriptomics of *T. brucei* and *T. congolense in vitro* samples. A.** and **B.** Plots showing the cutoffs (dashed lines) used to exclude cells with either low or high number of transcripts, number of genes, percentage of kDNA (mitochondrial) transcripts, or rRNA genes (top panels). Middle panels show the sample pre- and post-QC. Bottom panels show the cutoffs used to exclude cells that had a low percentage of *VSG* transcripts (of total transcripts) as dashed lines. The last four plots show the cells pre- and post-QC coloured by the percentage of *VSG* per cell as shown in the legend’s gradient. (A) shows the data for *T. brucei*, while (B) shows the data for *T. congolense*.

**Supplementary Figure 7. Single cell transcriptomics clusters.** Top panels show the Unifold manifold approximation and projection (UMAP) plots for *T. congolense* (left) and *T. brucei* (right), where cells are coloured by cluster as shown in Fig. 4A. Middle panels show dot plots of the top 5 genes identified for each cluster, while the bottom panels show the same information but as heatmaps; both indicate the clusters are relatively well-defined.

**Supplementary Figure 8. VSG expression levels in *T. brucei* and *T. congolense.* A.** Median expression level vs maximum expression level for every gene in *T. brucei* (left) and *T. congolense* (right); *VSG*s are coloured either blue (*T. brucei*) or purple (*T. congolense*). In *T. brucei* the gene with highest mean and maximum expression is *VSG* Tbr-000111900/AnTat1.1; in *T. congolense* the gene with highest mean and maximum expression is Tco-IL3000-001881300, an rRNA gene that was annotated as hypothetical in the genome; VSG Tco-IL3000-000997800 is also highlighted. **B.** Violin plots showing the expression levels of the top 50 *VSG*s (ordered by the number of cells with expression level >4) for *T. brucei* (top) and *T. congolense* (bottom).

**Supplementary Figure 9. Percentage of cells expressing different numbers of VSGs and the number of cells expressing specific combinations of VSGs. –A.** Stacked bar graph showing the percentage of cells expressing different numbers of *VSG* in the *in vitro* samples. **B.** Data relating to the *T. congolense in vitro* sample, where the number of cells expressing a specific combination of *VSG*s (indicated by the dot and connecting line graph) is shown; asterisks highlight the combinations shown in Fig.5D.

**Supplementary Figure 10. Expression of minichromosome-resident *T. congolense* VSGs in the scRNA-seq data.** Locations of all *VSG*s is shown on the *T. congolense* minichromosomes (<100 kb in size); those coloured dark green indicate expressed *VSG*s (expression >2), while those in light green are not detectably expressed. The location of telomeric repeats (TTAGGG) is indicated in grey.

**Supplementary Figure 11. Localisation of *VSG*s expressed in the scRNA-seq data on the larger chromosomes of *T. congolense*. A.** Locations of all *VSG*s is shown on the larger *T. congolense* chromosomes (the >800 kb megabase chromosomes, and two intermediate chromosomes, Tc_intermediate_02); those coloured green indicate expressed *VSG*s (expression >2), while those in black are not detectably expressed. B. –Distribution of genes around expressed *VSG*s with predicted functions based on Pfam domains.

**Supplementary Figure 12. *VSG* expression differs in culture and in mice**. **A.** *T. congolense* IL3000 *in vitro* (top, purple) and *in vivo* (bottom, salmon) data shown as: left panel, percentage of total *VSG* transcripts in the sample (left violin plot); middle panel, dot plot showing the expression level (y- axis) of a *VSG* in a given cell and how many reads (UMIs, x-axis) does that expression level correspond to (note: a cell is represented multiple times if expressing more than one VSG); right panel, sensitivity curve showing the number of cells called as expressing more than one *VSG* when one *VSG* is considered expressed if one, two or more UMIs are detected in the cell. **B.** Same as in (A) but for *T. brucei* samples. **C.** Percentage of cells expressing more than one *VSG* for all the samples when the expression threshold employed to call a *VSG* as expressed is set to 1, 1.5 or 2.

**Supplementary Figure 13. VSG co-expression sensitivity curves post SoupX.** Sensitivity curves are shown for four samples: *T. congolense* IL3000 *in vitro* (purple) and *in vivo* (salmon; only the day 6 sample was used), and *T. brucei in vitro* (blue) and *in vivo* (yellow; only the d7mS1 sample was used), after using SoupX to clean ambient RNA contamination using rho values of 0.05, 0.1 or 0.15 (corresponding to 5%, 10% and 15% contamination, respectively).

**Supplementary Table 1.**
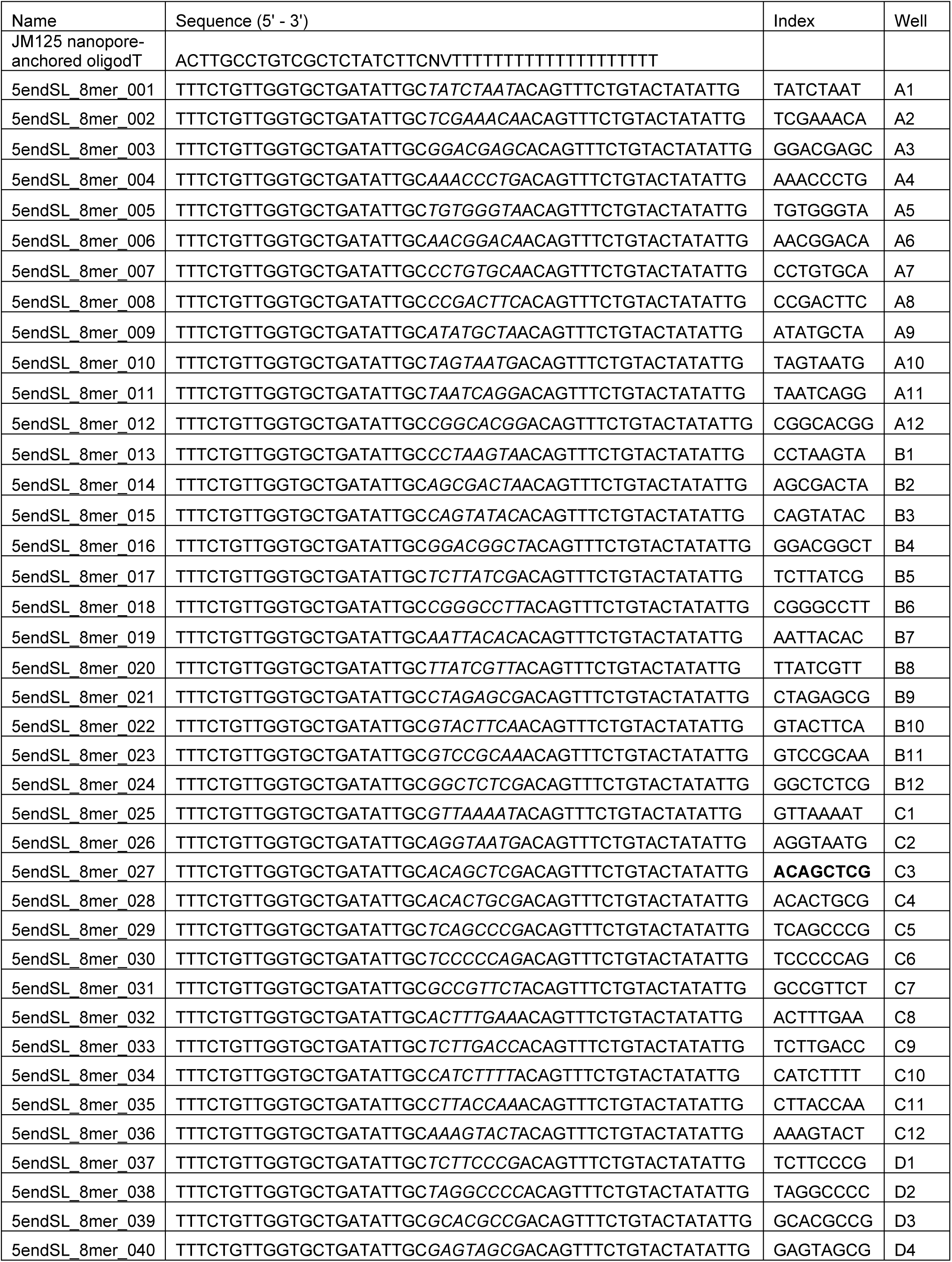
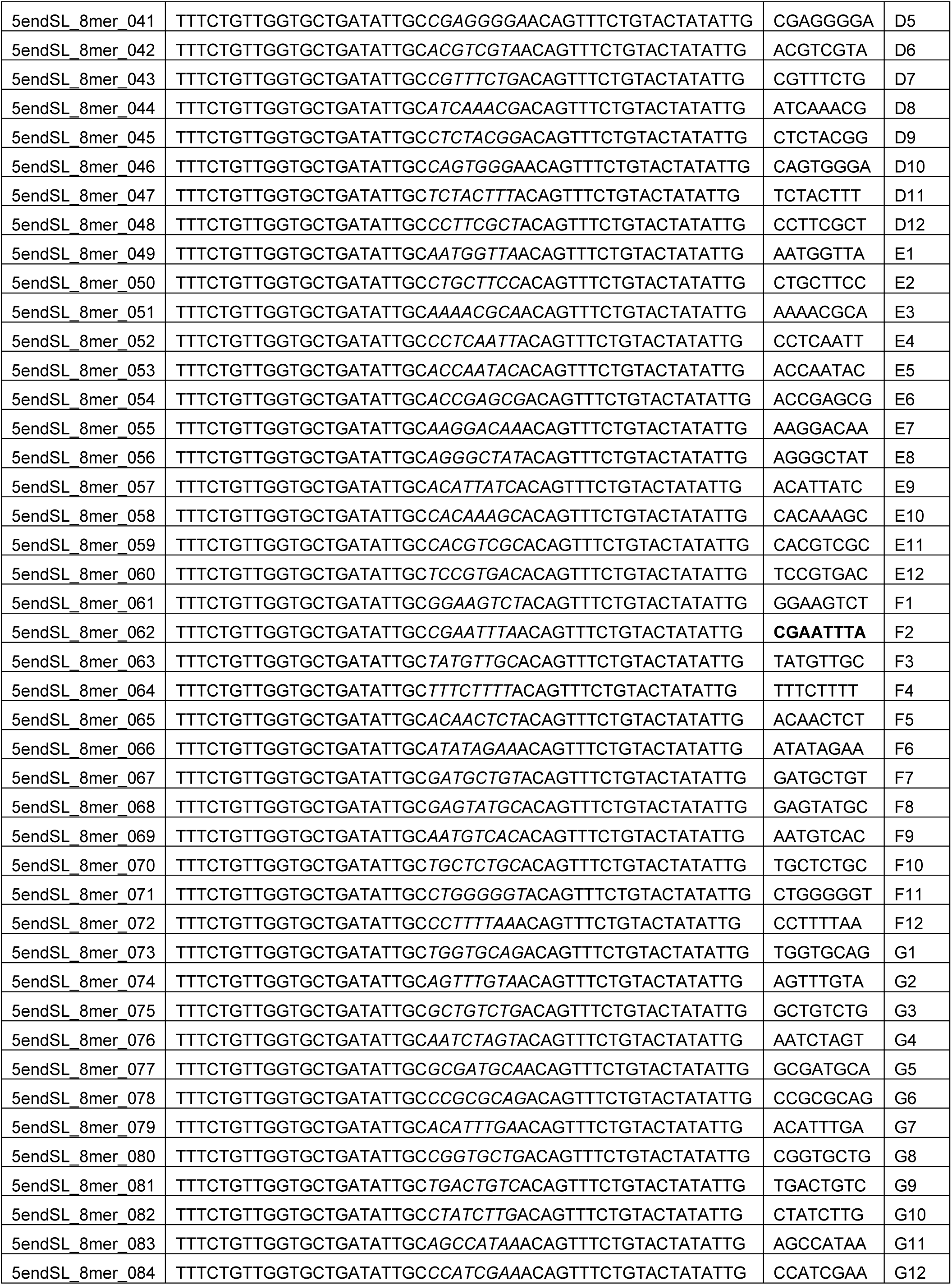
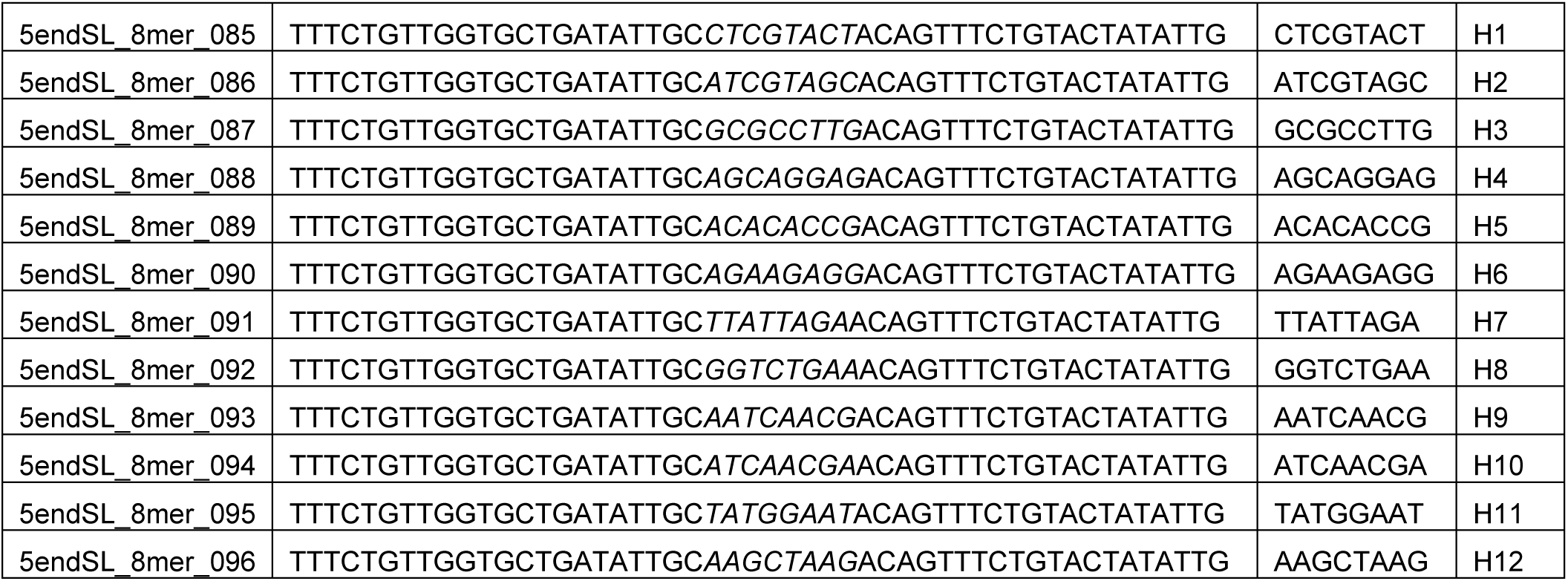
Primers used for SL-barseq.

## References

1 Deitsch, K. W., Lukehart, S. A. & Stringer, J. R. Common strategies for antigenic variation by bacterial, fungal and protozoan pathogens. Nat.Rev.Microbiol. 7, 493–503 (2009).

2 Barcons-Simon, A., Carrington, M. & Siegel, T. N. Decoding the impact of nuclear organization on antigenic variation in parasites. Nat Microbiol 8, 1408–1418 (2023). 10.1038/s41564-023-01424-9

3 Borst, P. Antigenic variation and allelic exclusion. Cell 109, 5–8 (2002). 10.1016/s0092-8674(02)00711-0

4 Florini, F., Visone, J. E. & Deitsch, K. W. Shared Mechanisms for Mutually Exclusive Expression and Antigenic Variation by Protozoan Parasites. Front Cell Dev Biol 10, 852239 (2022). 10.3389/fcell.2022.852239

5 Palmer, G. H., Bankhead, T. & Seifert, H. S. Antigenic Variation in Bacterial Pathogens.Microbiology spectrum 4 (2016). 10.1128/microbiolspec.VMBF-0005-2015

6 Diffendall, G. & Scherf, A. Deciphering the Plasmodium falciparum perinuclear var gene expression site. Trends in parasitology 40, 707–716 (2024). 10.1016/j.pt.2024.06.002

7 Hadjimichael, E. & Deitsch, K. W. Variable surface antigen expression, virulence, and persistent infection by Plasmodium falciparum malaria parasites. Microbiology and molecular biology reviews : MMBR 89, e0011423 (2025). 10.1128/mmbr.00114-23

8 Akpo, Y., Tonouhewa, B. N. A., Farougou, S., Alkoiret, T. & Kpodekon, M. African animal trypanosomosis among cattle in West Africa: A meta-analysis of epidemiological record (2000-2025). Vet Parasitol Reg Stud Reports 69, 101451 (2026). 10.1016/j.vprsr.2026.101451

9 Morrison, L. J., Steketee, P. C., Tettey, M. D. & Matthews, K. R. Pathogenicity and virulence of African trypanosomes: From laboratory models to clinically relevant hosts. Virulence 14, 2150445 (2023). 10.1080/21505594.2022.2150445

10 Berriman, M. et al. The genome of the African trypanosome Trypanosoma brucei. Science 309, 416–422 (2005).

11 Marcello, L. & Barry, J. D. Analysis of the VSG gene silent archive in Trypanosoma brucei reveals that mosaic gene expression is prominent in antigenic variation and is favored by archive substructure. Genome Res. 17, 1344–1352 (2007).

12 Cross, G. A., Kim, H. S. & Wickstead, B. Capturing the variant surface glycoprotein repertoire (the VSGnome) of Trypanosoma brucei Lister 427. Molecular and biochemical parasitology 195, 59–73 (2014). 10.1016/j.molbiopara.2014.06.004

13 Hertz-Fowler, C. et al. Telomeric expression sites are highly conserved in Trypanosoma brucei. PLoS ONE. 3, e3527 (2008).

14 Girasol, M. J. et al. RAD51-mediated R-loop formation acts to repair transcription-associated DNA breaks driving antigenic variation in Trypanosoma brucei. Proceedings of the National Academy of Sciences of the United States of America 120, e2309306120 (2023). 10.1073/pnas.2309306120

15 McCulloch, R. & Barry, J. D. A role for RAD51 and homologous recombination in Trypanosoma brucei antigenic variation. Genes & development 13, 2875–2888 (1999).

16 Hartley, C. L. & McCulloch, R. Trypanosoma brucei BRCA2 acts in antigenic variation and has undergone a recent expansion in BRC repeat number that is important during homologous recombination. Mol.Microbiol. 68, 1237–1251 (2008).

17 Smith, J. E. et al. DNA damage drives antigen diversification in Trypanosoma brucei. Nature (2026). 10.1038/s41586-026-10337-6

18 Hall, J. P., Wang, H. & Barry, J. D. Mosaic VSGs and the scale of Trypanosoma brucei antigenic variation. PLoS pathogens 9, e1003502 (2013). 10.1371/journal.ppat.1003502

19 Mugnier, M. R., Cross, G. A. & Papavasiliou, F. N. The in vivo dynamics of antigenic variation in Trypanosoma brucei. Science 347, 1470–1473 (2015). 10.1126/science.aaa4502

20 Jayaraman, S. et al. Application of long read sequencing to determine expressed antigen diversity in Trypanosoma brucei infections. PLoS neglected tropical diseases 13, e0007262 (2019). 10.1371/journal.pntd.0007262

21 Borst, P. & Ulbert, S. Control of VSG gene expression sites. Molecular and biochemical parasitology 114, 17–27 (2001). 10.1016/s0166-6851(01)00243-2

22 Chaves, I., Rudenko, G., Dirks-Mulder, A., Cross, M. & Borst, P. Control of variant surface glycoprotein gene-expression sites in Trypanosoma brucei. The EMBO journal 18, 4846–4855 (1999). 10.1093/emboj/18.17.4846

23 Ulbert, S., Chaves, I. & Borst, P. Expression site activation in Trypanosoma brucei with three marked variant surface glycoprotein gene expression sites. Mol.Biochem.Parasitol. 120, 225– 235 (2002).

24 Budzak, J. et al. Dynamic colocalization of 2 simultaneously active VSG expression sites within a single expression-site body in Trypanosoma brucei. Proceedings of the National Academy of Sciences of the United States of America 116, 16561–16570 (2019). 10.1073/pnas.1905552116

25 Faria, J., Briggs, E. M., Black, J. A. & McCulloch, R. Emergence and adaptation of the cellular machinery directing antigenic variation in the African trypanosome. Current opinion in microbiology 70, 102209 (2022). 10.1016/j.mib.2022.102209

26 Glover, L., Hutchinson, S., Alsford, S. & Horn, D. VEX1 controls the allelic exclusion required for antigenic variation in trypanosomes. Proceedings of the National Academy of Sciences of the United States of America 113, 7225–7230 (2016). 10.1073/pnas.1600344113

27 Faria, J. et al. Monoallelic expression and epigenetic inheritance sustained by a Trypanosoma brucei variant surface glycoprotein exclusion complex. Nature communications 10, 3023 (2019). 10.1038/s41467-019-10823-8

28 Faria, J. et al. Spatial integration of transcription and splicing in a dedicated compartment sustains monogenic antigen expression in African trypanosomes. Nat Microbiol 6, 289–300 (2021). 10.1038/s41564-020-00833-4

29 Faria, J. R. C. et al. An allele-selective inter-chromosomal protein bridge supports monogenic antigen expression in the African trypanosome. Nature communications 14, 8200 (2023). 10.1038/s41467-023-44043-y

30 Navarro, M. & Gull, K. A pol I transcriptional body associated with VSG mono-allelic expression in Trypanosoma brucei. Nature 414, 759–763 (2001).

31 Lopez-Escobar, L. et al. Stage-specific transcription activator ESB1 regulates monoallelic antigen expression in Trypanosoma brucei. Nat Microbiol 7, 1280–1290 (2022). 10.1038/s41564-022-01175-z

32 Berazategui, M. A. et al. A factor integrating transcription and repression of surface antigen genes in African trypanosomes. Proceedings of the National Academy of Sciences of the United States of America 123, e2531377123 (2026). 10.1073/pnas.2531377123

33 Lansink, L. I. M. et al. Specialized RNA decay fine-tunes monogenic antigen expression in Trypanosoma brucei. Nat Microbiol 11, 1080–1099 (2026). 10.1038/s41564-026-02289-4

34 Steketee, P. C. et al. Divergent metabolism between Trypanosoma congolense and Trypanosoma brucei results in differential sensitivity to metabolic inhibition. PLoS pathogens 17, e1009734 (2021). 10.1371/journal.ppat.1009734

35 Awuah-Mensah, G. et al. Reliable, scalable functional genetics in bloodstream-form Trypanosoma congolense in vitro and in vivo. PLoS pathogens 17, e1009224 (2021). 10.1371/journal.ppat.1009224

36 Jackson, A. P. et al. Antigenic diversity is generated by distinct evolutionary mechanisms in African trypanosome species. Proc.Natl.Acad.Sci.U.S.A 109, 3416–3421 (2012).

37 Silva Pereira, S., Jackson, A. P. & Figueiredo, L. M. Evolution of the variant surface glycoprotein family in African trypanosomes. Trends in parasitology 38, 23–36 (2022). 10.1016/j.pt.2021.07.012

38 Otesile, E. B. & Tabel, H. Enhanced resistance of highly susceptible Balb/c mice to infection with Trypanosoma congolense after infection and cure. J Parasitol 73, 947–953 (1987).

39 Sutherland, D. V., Ross, C. A. & Luckins, A. G. Trypanosoma congolense: re-expression of a deleted metacyclic variable antigen type in vivo and in vitro. Acta tropica 49, 193–199 (1991). 10.1016/0001-706x(91)90038-l

40 Majiwa, P. A., Matthyssens, G., Williams, R. O. & Hamers, R. Cloning and analysis of Trypanosoma (Nannomonas) congolense ILNat 2.1 VSG gene. Molecular and biochemical parasitology 16, 97–108 (1985). 10.1016/0166-6851(85)90052-0

41 Carrington, M. et al. Variant specific glycoprotein of Trypanosoma brucei consists of two domains each having an independently conserved pattern of cysteine residues. J.Mol.Biol. 221, 823–835 (1991).

42 Silva Pereira, S., et al. Variant antigen repertoires in Trypanosoma congolense populations and experimental infections can be profiled from deep sequence data using universal protein motifs. Genome research 28, 1383–1394 (2018). 10.1101/gr.234146.118

43 Abbas, A. H. et al. The Structure of a Conserved Telomeric Region Associated with Variant Antigen Loci in the Blood Parasite Trypanosoma congolense. Genome biology and evolution 10, 2458–2473 (2018). 10.1093/gbe/evy186

44 Majiwa, P. A., Young, J. R., Hamers, R. & Matthyssens, G. Minichromosomal variable surface glycoprotein genes and molecular karyotypes of Trypanosoma (Nannomonas) congolense. Gene 41, 183–192 (1986). 10.1016/0378-1119(86)90097-1

45 Wickstead, B., Ersfeld, K. & Gull, K. The small chromosomes of Trypanosoma brucei involved in antigenic variation are constructed around repetitive palindromes. Genome research 14, 1014–1024 (2004). 10.1101/gr.2227704

46 Haese-Hill, W., Crouch, K. & Otto, T. D. Annotation and visualization of parasite, fungi and arthropod genomes with Companion. Nucleic Acids Res 52, W39–W44 (2024). 10.1093/nar/gkae378

47 Patro, R., Duggal, G., Love, M. I., Irizarry, R. A. & Kingsford, C. Salmon provides fast and bias- aware quantification of transcript expression. Nature methods 14, 417–419 (2017). 10.1038/nmeth.4197

48 Oldrieve, G., Larcombe, S., Krasilnikova, M., Mugnier, M. & Matthews, K. vsgseq2: an updated pipeline for analysis of the diversity and abundance of population-wide Trypanosoma brucei VSG expression. Wellcome Open Res 10, 672 (2025). 10.12688/wellcomeopenres.24932.1

49 Silva Pereira, S., Heap, J., Jones, A. R. & Jackson, A. P. VAPPER: High-throughput variant antigen profiling in African trypanosomes of livestock. Gigascience 8 (2019). 10.1093/gigascience/giz091

50 Silvester, E., Ivens, A. & Matthews, K. R. A gene expression comparison of Trypanosoma brucei and Trypanosoma congolense in the bloodstream of the mammalian host reveals species-specific adaptations to density-dependent development. PLoS neglected tropical diseases 12, e0006863 (2018). 10.1371/journal.pntd.0006863

51 Silva Pereira, S., Mathenge, K., Masiga, D. & Jackson, A. Transcriptomic profiling of Trypanosoma congolense mouthpart parasites from naturally infected flies. Parasit Vectors 15, 152 (2022). 10.1186/s13071-022-05258-y

52 Touray, A. O., Sternlieb, T., Isebe, T. & Cestari, I. Identifying Antigenic Switching by Clonal Cell Barcoding and Nanopore Sequencing in Trypanosoma brucei. Bio Protoc 13, e4904 (2023). 10.21769/BioProtoc.4904

53 Devlin, R. et al. Mapping replication dynamics in Trypanosoma brucei reveals a link with telomere transcription and antigenic variation. eLife 5, e12765 (2016). 10.7554/eLife.12765

54 Krasilnikova, M. et al. Nanopore sequencing reveals that DNA replication compartmentalisation dictates genome stability and instability in Trypanosoma brucei. Nature communications 16, 751 (2025). 10.1038/s41467-025-56087-3

55 Briggs, E., Crouch, K., Lemgruber, L., Lapsley, C. & McCulloch, R. Ribonuclease H1-targeted R- loops in surface antigen gene expression sites can direct trypanosome immune evasion. PLoS genetics 14, e1007729 (2018). 10.1371/journal.pgen.1007729

56 Cross, G. A. Release and purification of Trypanosoma brucei variant surface glycoprotein. Journal of cellular biochemistry 24, 79–90 (1984). 10.1002/jcb.240240107

57 Ishihama, Y. et al. Exponentially modified protein abundance index (emPAI) for estimation of absolute protein amount in proteomics by the number of sequenced peptides per protein. Molecular & cellular proteomics : MCP 4, 1265–1272 (2005). 10.1074/mcp.M500061-MCP200

58 Chavez, S. et al. Extensive Translational Regulation through the Proliferative Transition of Trypanosoma cruzi Revealed by Multi-Omics. mSphere 6, e0036621 (2021). 10.1128/mSphere.00366-21

59 Briggs, E. M., Rojas, F., McCulloch, R., Matthews, K. R. & Otto, T. D. Single-cell transcriptomic analysis of bloodstream Trypanosoma brucei reconstructs cell cycle progression and developmental quorum sensing. Nature communications 12, 5268 (2021). 10.1038/s41467-021-25607-2

60 Larcombe, S. D., Briggs, E. M., Savill, N., Szoor, B. & Matthews, K. R. The developmental hierarchy and scarcity of replicative slender trypanosomes in blood challenges their role in infection maintenance. Proceedings of the National Academy of Sciences of the United States of America 120, e2306848120 (2023). 10.1073/pnas.2306848120

61 Larcombe, S. D. et al. Trypanosoma brucei cattle infections contain cryptic transmission- adapted bloodstream forms at low parasitaemia. Nature communications 16, 9776 (2025). 10.1038/s41467-025-64750-y

62 Muller, L. S. M. et al. Genome organization and DNA accessibility control antigenic variation in trypanosomes. Nature 563, 121–125 (2018). 10.1038/s41586-018-0619-8

63 Keneskhanova, Z. et al. Genomic determinants of antigen expression hierarchy in African trypanosomes. Nature (2025). 10.1038/s41586-025-08720-w

64 Mony, B. M. et al. Genome-wide dissection of the quorum sensing signalling pathway in Trypanosoma brucei. Nature 505, 681–685 (2014). 10.1038/nature12864

65 McWilliam, K. R., Ivens, A., Morrison, L. J., Mugnier, M. R. & Matthews, K. R. Developmental competence and antigen switch frequency can be uncoupled in Trypanosoma brucei. Proceedings of the National Academy of Sciences of the United States of America 116, 22774–22782 (2019). 10.1073/pnas.1912711116

66 Silvester, E. et al. A conserved trypanosomatid differentiation regulator controls substrate attachment and morphological development in Trypanosoma congolense. PLoS pathogens 20, e1011889 (2024). 10.1371/journal.ppat.1011889

67 Briggs, E. M. et al. Profiling the bloodstream form and procyclic form Trypanosoma brucei cell cycle using single-cell transcriptomics. eLife 12 (2023). 10.7554/eLife.86325

68 Beaver, A. K. et al. Tissue spaces are reservoirs of antigenic diversity for Trypanosoma brucei. Nature (2024). 10.1038/s41586-024-08151-z

69 Amiguet-Vercher, A. et al. Loss of the mono-allelic control of the VSG expression sites during the development of Trypanosoma brucei in the bloodstream. Molecular microbiology 51, 1577–1588 (2004). 10.1111/j.1365-2958.2003.03937.x

70 Macgregor, P., Savill, N. J., Hall, D. & Matthews, K. R. Transmission stages dominate trypanosome within-host dynamics during chronic infections. Cell Host.Microbe 9, 310–318 (2011).

71 Silvester, E., Young, J., Ivens, A. & Matthews, K. R. Interspecies quorum sensing in co- infections can manipulate trypanosome transmission potential. Nat Microbiol 2, 1471–1479 (2017). 10.1038/s41564-017-0014-5

72 Tihon, E. et al. Genomic analysis of Isometamidium Chloride resistance in Trypanosoma congolense. Int J Parasitol Drugs Drug Resist 7, 350–361 (2017). 10.1016/j.ijpddr.2017.10.002

73 Kassem, A., Pays, E. & Vanhamme, L. Transcription is initiated on silent variant surface glycoprotein expression sites despite monoallelic expression in Trypanosoma brucei. Proceedings of the National Academy of Sciences of the United States of America 111, 8943– 8948 (2014). 10.1073/pnas.1404873111

74 Berberof, M. et al. The 3’-terminal region of the mRNAs for VSG and procyclin can confer stage specificity to gene expression in Trypanosoma brucei. EMBO J. 14, 2925–2934 (1995).

75 Ridewood, S. et al. The role of genomic location and flanking 3’UTR in the generation of functional levels of variant surface glycoprotein in Trypanosoma brucei. Molecular microbiology 106, 614–634 (2017). 10.1111/mmi.13838

76 Melo do Nascimento, L., et al. Functional insights from a surface antigen mRNA-bound proteome. eLife 10 (2021). 10.7554/eLife.68136

77 Viegas, I. J. et al. N(6)-methyladenosine in poly(A) tails stabilize VSG transcripts. Nature 604, 362–370 (2022). 10.1038/s41586-022-04544-0

78 Bakari-Soale, M., Batram, C., Zimmermann, H., Jones, N. G. & Engstler, M. The Dual Role of the 16mer Motif Within the 3’ Untranslated Region of the Variant Surface Glycoprotein of Trypanosoma brucei. Molecular microbiology 125, 67–79 (2026). 10.1111/mmi.70031

79 Escrivani, D. O. et al. A non-coding role for trypanosome VSG transcripts in allelic exclusion. Nucleic Acids Res 53 (2025). 10.1093/nar/gkaf1011

80 McCulloch, R. & Field, M. C. Quantitative sequencing confirms VSG diversity as central to immune evasion by Trypanosoma brucei. Trends in parasitology 31, 346–349 (2015). 10.1016/j.pt.2015.05.001

81 Aresta-Branco, F., Sanches-Vaz, M., Bento, F., Rodrigues, J. A. & Figueiredo, L. M. African trypanosomes expressing multiple VSGs are rapidly eliminated by the host immune system. Proceedings of the National Academy of Sciences of the United States of America 116, 20725–20735 (2019). 10.1073/pnas.1905120116

82 Goossens, B., Osaer, S., Kora, S. & Ndao, M. Haematological changes and antibody response in trypanotolerant sheep and goats following experimental Trypanosoma congolense infection. Vet Parasitol 79, 283–297 (1998). 10.1016/s0304-4017(98)00171-x

83 Rurangirwa, F. R., Musoke, A. J., Nantulya, V. M. & Tabel, H. Immune depression in bovine trypanosomiasis: effects of acute and chronic Trypanosoma congolense and chronic Trypanosoma vivax infections on antibody response to Brucella abortus vaccine. Parasite Immunol 5, 267–276 (1983). 10.1111/j.1365-3024.1983.tb00743.x

84 Smith, J. E. et al. DNA damage drives antigen diversification in Trypanosoma brucei. Nature 654, 219–228 (2026). 10.1038/s41586-026-10337-6

85 Silva Pereira, S., et al. Variant antigen diversity in Trypanosoma vivax is not driven by recombination. Nature communications 11, 844 (2020). 10.1038/s41467-020-14575-8

86 Sheader, K., Berberof, M., Isobe, T., Borst, P. & Rudenko, G. Delineation of the regulated Variant Surface Glycoprotein gene expression site domain of Trypanosoma brucei. Molecular and biochemical parasitology 128, 147–156 (2003). 10.1016/s0166-6851(03)00056-2

87 Liu, A. Y., Van der Ploeg, L. H., Rijsewijk, F. A. & Borst, P. The transposition unit of variant surface glycoprotein gene 118 of Trypanosoma brucei. Presence of repeated elements at its border and absence of promoter-associated sequences. Journal of molecular biology 167, 57–75 (1983). 10.1016/s0022-2836(83)80034-5

88 McCulloch, R., Rudenko, G. & Borst, P. Gene conversions mediating antigenic variation in Trypanosoma brucei can occur in variant surface glycoprotein expression sites lacking 70- base-pair repeat sequences. Mol.Cell Biol. 17, 833–843 (1997).

89 Kieft, R. et al. Mono-allelic epigenetic regulation of polycistronic transcription initiation by RNA polymerase II in Trypanosoma brucei. mBio 16, e0232824 (2025). 10.1128/mbio.02328-24

90 Ginger, M. L. et al. Ex Vivo and In Vitro Identification of a Consensus Promoter for VSG Genes Expressed by Metacyclic-Stage Trypanosomes in the Tsetse Fly. Eukaryot.Cell 1, 1000–1009 (2002).

91 Crowe, J. S., Barry, J. D., Luckins, A. G., Ross, C. A. & Vickerman, K. All metacyclic variable antigen types of Trypanosoma congolense identified using monoclonal antibodies. Nature 306, 389–391 (1983). 10.1038/306389a0

92 Luckins, A. G., Frame, I. A., Gray, M. A., Crowe, J. S. & Ross, C. A. Analysis of trypanosome variable antigen types in cultures of metacyclic and mammalian forms of Trypanosoma congolense. Parasitology 93 **(Pt** **1**), 99–109 (1986). 10.1017/s0031182000049854

93 Barbet, A. F. & Kamper, S. M. The importance of mosaic genes to trypanosome survival. Parasitol.Today 9, 63–66 (1993).

94 Barry, J. D., Hall, J. P. & Plenderleith, L. Genome hyperevolution and the success of a parasite. Ann.N.Y.Acad.Sci. 1267, 11–17 (2012).

95 McCulloch, R. et al. Emerging challenges in understanding trypanosome antigenic variation. Emerg Top Life Sci 1, 585–592 (2017). 10.1042/ETLS20170104

96 Florini, F. et al. scRNA-seq reveals transcriptional plasticity of var gene expression in Plasmodium falciparum for host immune avoidance. Nat Microbiol 10, 1417–1430 (2025). 10.1038/s41564-025-02008-5

97 Hutchinson, S. et al. The establishment of variant surface glycoprotein monoallelic expression revealed by single-cell RNA-seq of Trypanosoma brucei in the tsetse fly salivary glands. PLoS pathogens 17, e1009904 (2021). 10.1371/journal.ppat.1009904

98 Allred, D. R. Variable and Variant Protein Multigene Families in Babesia bovis Persistence. Pathogens 8 (2019). 10.3390/pathogens8020076

99 Wahlgren, M., Goel, S. & Akhouri, R. R. Variant surface antigens of Plasmodium falciparum and their roles in severe malaria. Nature reviews. Microbiology 15, 479–491 (2017). 10.1038/nrmicro.2017.47

100 Schmid-Siegert, E. et al. Mechanisms of Surface Antigenic Variation in the Human Pathogenic Fungus Pneumocystis jirovecii. mBio 8 (2017). 10.1128/mBio.01470-17

101 Inchausti, L. et al. Single-cell RNA-seq reveals trans-sialidase-like superfamily gene expression heterogeneity in Trypanosoma cruzi populations. eLife 14 (2026). 10.7554/eLife.105822

102 Cruz-Saavedra, L., Loock, M., Antunes, L. B. & Cestari, I. Variation in surface protein expression leads to heterogeneous Trypanosoma cruzi populations during host cell infection. Nature communications 16, 9949 (2025). 10.1038/s41467-025-64900-2

103 Dickson, K. P., Costales, J. A., Domagalska, M. A., Vander Veken, F. & Llewellyn, M. S. Innovation through instability? Genome (dis)organisation in Trypanosoma cruzi. Trends in parasitology 41, 449–459 (2025). 10.1016/j.pt.2025.04.008

104 Kelly, S. et al. An Alternative Strategy for Trypanosome Survival in the Mammalian Bloodstream Revealed through Genome and Transcriptome Analysis of the Ubiquitous Bovine Parasite Trypanosoma (Megatrypanum) theileri. Genome biology and evolution 9, 2093–2109 (2017). 10.1093/gbe/evx152

105 Hirumi, H. & Hirumi, K. In vitro cultivation of Trypanosoma congolense bloodstream forms in the absence of feeder cell layers. Parasitology 102 **Pt** **2**, 225–236 (1991). 10.1017/s0031182000062533

106 Masumu, J., Geysen, D. & Van den Bossche, P. Endemic type of animal trypanosomiasis is not associated with lower genotype variability of Trypanosoma congolense isolates circulating in livestock. Res Vet Sci 87, 265–269 (2009). 10.1016/j.rvsc.2009.03.003

