## Supplementary material for "*Trypanosoma congolense* Variant Surface Glycoprotein gene expression occurs in the absence of monoallelic transcription control": All supplementary Figures and a Supp Table, plus legends

**Supplementary Figure 13. VSG co-expression sensitivity curves post SoupX.** Sensitivity curves are shown for four samples: *T. congolense* IL3000 *in vitro* (purple) and *in vivo* (salmon; only the day 6 sample was used), and *T. brucei in vitro* (blue) and *in vivo* (yellow; only the d7mS1 sample was used), after using SoupX to clean ambient RNA contamination using rho values of 0.05, 0.1 or 0.15 (corresponding to 5%, 10% and 15% contamination, respectively).

Supplementary Table 1 – Primers used for SL-barseq

| Name | Sequence (5' - 3') | Index | Well |
|---|---|---|---|
| JM125 nanopore-anchored oligodT | ACTTGCCTGTCGCTCTATCTTCNVTTTTTTTTTTTTTTTTTTTT | | |
| 5endSL_8mer_001 | TTTCTGTTGGTGCTGATATTGC*TATCTAAT*ACAGTTTCTGTACTATATTG | TATCTAAT | A1 |
| 5endSL_8mer_002 | TTTCTGTTGGTGCTGATATTGC*TCGAAACA*ACAGTTTCTGTACTATATTG | TCGAAACA | A2 |
| 5endSL_8mer_003 | TTTCTGTTGGTGCTGATATTGC*GGACGAGC*ACAGTTTCTGTACTATATTG | GGACGAGC | A3 |
| 5endSL_8mer_004 | TTTCTGTTGGTGCTGATATTGC*AAACCCTG*ACAGTTTCTGTACTATATTG | AAACCCTG | A4 |
| 5endSL_8mer_005 | TTTCTGTTGGTGCTGATATTGC*TGTGGGTA*ACAGTTTCTGTACTATATTG | TGTGGGTA | A5 |
| 5endSL_8mer_006 | TTTCTGTTGGTGCTGATATTGC*AACGGACA*ACAGTTTCTGTACTATATTG | AACGGACA | A6 |
| 5endSL_8mer_007 | TTTCTGTTGGTGCTGATATTGC*CCTGTGCA*ACAGTTTCTGTACTATATTG | CCTGTGCA | A7 |
| 5endSL_8mer_008 | TTTCTGTTGGTGCTGATATTGC*CCGACTTC*ACAGTTTCTGTACTATATTG | CCGACTTC | A8 |
| 5endSL_8mer_009 | TTTCTGTTGGTGCTGATATTGC*ATATGCTA*ACAGTTTCTGTACTATATTG | ATATGCTA | A9 |
| 5endSL_8mer_010 | TTTCTGTTGGTGCTGATATTGC*TAGTAATG*ACAGTTTCTGTACTATATTG | TAGTAATG | A10 |
| 5endSL_8mer_011 | TTTCTGTTGGTGCTGATATTGC*TAATCAGG*ACAGTTTCTGTACTATATTG | TAATCAGG | A11 |
| 5endSL_8mer_012 | TTTCTGTTGGTGCTGATATTGC*CGGCACGG*ACAGTTTCTGTACTATATTG | CGGCACGG | A12 |
| 5endSL_8mer_013 | TTTCTGTTGGTGCTGATATTGC*CCTAAGTA*ACAGTTTCTGTACTATATTG | CCTAAGTA | B1 |
| 5endSL_8mer_014 | TTTCTGTTGGTGCTGATATTGC*AGCGACTA*ACAGTTTCTGTACTATATTG | AGCGACTA | B2 |
| 5endSL_8mer_015 | TTTCTGTTGGTGCTGATATTGC*CAGTATAC*ACAGTTTCTGTACTATATTG | CAGTATAC | B3 |
| 5endSL_8mer_016 | TTTCTGTTGGTGCTGATATTGC*GGACGGCT*ACAGTTTCTGTACTATATTG | GGACGGCT | B4 |
| 5endSL_8mer_017 | TTTCTGTTGGTGCTGATATTGC*TCTTATCG*ACAGTTTCTGTACTATATTG | TCTTATCG | B5 |
| 5endSL_8mer_018 | TTTCTGTTGGTGCTGATATTGC*CGGGCCTT*ACAGTTTCTGTACTATATTG | CGGGCCTT | B6 |
| 5endSL_8mer_019 | TTTCTGTTGGTGCTGATATTGC*AATTACAC*ACAGTTTCTGTACTATATTG | AATTACAC | B7 |
| 5endSL_8mer_020 | TTTCTGTTGGTGCTGATATTGC*TTATCGTT*ACAGTTTCTGTACTATATTG | TTATCGTT | B8 |
| 5endSL_8mer_021 | TTTCTGTTGGTGCTGATATTGC*CTAGAGCG*ACAGTTTCTGTACTATATTG | CTAGAGCG | B9 |
| 5endSL_8mer_022 | TTTCTGTTGGTGCTGATATTGC*GTACTTCA*ACAGTTTCTGTACTATATTG | GTACTTCA | B10 |
| 5endSL_8mer_023 | TTTCTGTTGGTGCTGATATTGC*GTCCGCAA*ACAGTTTCTGTACTATATTG | GTCCGCAA | B11 |
| 5endSL_8mer_024 | TTTCTGTTGGTGCTGATATTGC*GGCTCTCG*ACAGTTTCTGTACTATATTG | GGCTCTCG | B12 |
| 5endSL_8mer_025 | TTTCTGTTGGTGCTGATATTGC*GTTAAAAT*ACAGTTTCTGTACTATATTG | GTTAAAAT | C1 |
| 5endSL_8mer_026 | TTTCTGTTGGTGCTGATATTGC*AGGTAATG*ACAGTTTCTGTACTATATTG | AGGTAATG | C2 |
| 5endSL_8mer_027 | TTTCTGTTGGTGCTGATATTGC*ACAGCTCG*ACAGTTTCTGTACTATATTG | **ACAGCTCG** | C3 |
| 5endSL_8mer_028 | TTTCTGTTGGTGCTGATATTGC*ACACTGCG*ACAGTTTCTGTACTATATTG | ACACTGCG | C4 |
| 5endSL_8mer_029 | TTTCTGTTGGTGCTGATATTGC*TCAGCCCG*ACAGTTTCTGTACTATATTG | TCAGCCCG | C5 |
| 5endSL_8mer_030 | TTTCTGTTGGTGCTGATATTGC*TCCCCCAG*ACAGTTTCTGTACTATATTG | TCCCCCAG | C6 |
| 5endSL_8mer_031 | TTTCTGTTGGTGCTGATATTGC*GCCGTTCT*ACAGTTTCTGTACTATATTG | GCCGTTCT | C7 |
| 5endSL_8mer_032 | TTTCTGTTGGTGCTGATATTGC*ACTTTGAA*ACAGTTTCTGTACTATATTG | ACTTTGAA | C8 |
| 5endSL_8mer_033 | TTTCTGTTGGTGCTGATATTGC*TCTTGACC*ACAGTTTCTGTACTATATTG | TCTTGACC | C9 |
| 5endSL_8mer_034 | TTTCTGTTGGTGCTGATATTGC*CATCTTTT*ACAGTTTCTGTACTATATTG | CATCTTTT | C10 |
| 5endSL_8mer_035 | TTTCTGTTGGTGCTGATATTGC*CTTACCAA*ACAGTTTCTGTACTATATTG | CTTACCAA | C11 |
| 5endSL_8mer_036 | TTTCTGTTGGTGCTGATATTGC*AAAGTACT*ACAGTTTCTGTACTATATTG | AAAGTACT | C12 |
| 5endSL_8mer_037 | TTTCTGTTGGTGCTGATATTGC*TCTTCCCG*ACAGTTTCTGTACTATATTG | TCTTCCCG | D1 |
| 5endSL_8mer_038 | TTTCTGTTGGTGCTGATATTGC*TAGGCCCC*ACAGTTTCTGTACTATATTG | TAGGCCCC | D2 |
| 5endSL_8mer_039 | TTTCTGTTGGTGCTGATATTGC*GCACGCCG*ACAGTTTCTGTACTATATTG | GCACGCCG | D3 |
| 5endSL_8mer_040 | TTTCTGTTGGTGCTGATATTGC*GAGTAGCG*ACAGTTTCTGTACTATATTG | GAGTAGCG | D4 |

| | | | |
|---|---|---|---|
| 5endSL_8mer_041 | TTTCTGTTGGTGCTGATATTGC*CGAGGGGA*ACAGTTTCTGTACTATATTG | CGAGGGGA | D5 |
| 5endSL_8mer_042 | TTTCTGTTGGTGCTGATATTGC*ACGTCGTA*ACAGTTTCTGTACTATATTG | ACGTCGTA | D6 |
| 5endSL_8mer_043 | TTTCTGTTGGTGCTGATATTGC*CGTTTCTG*ACAGTTTCTGTACTATATTG | CGTTTCTG | D7 |
| 5endSL_8mer_044 | TTTCTGTTGGTGCTGATATTGC*ATCAAACG*ACAGTTTCTGTACTATATTG | ATCAAACG | D8 |
| 5endSL_8mer_045 | TTTCTGTTGGTGCTGATATTGC*CTCTACGG*ACAGTTTCTGTACTATATTG | CTCTACGG | D9 |
| 5endSL_8mer_046 | TTTCTGTTGGTGCTGATATTGC*CAGTGGGA*ACAGTTTCTGTACTATATTG | CAGTGGGA | D10 |
| 5endSL_8mer_047 | TTTCTGTTGGTGCTGATATTGC*TCTACTTT*ACAGTTTCTGTACTATATTG | TCTACTTT | D11 |
| 5endSL_8mer_048 | TTTCTGTTGGTGCTGATATTGC*CCTTCGCT*ACAGTTTCTGTACTATATTG | CCTTCGCT | D12 |
| 5endSL_8mer_049 | TTTCTGTTGGTGCTGATATTGC*AATGGTTA*ACAGTTTCTGTACTATATTG | AATGGTTA | E1 |
| 5endSL_8mer_050 | TTTCTGTTGGTGCTGATATTGC*CTGCTTCC*ACAGTTTCTGTACTATATTG | CTGCTTCC | E2 |
| 5endSL_8mer_051 | TTTCTGTTGGTGCTGATATTGC*AAAACGCA*ACAGTTTCTGTACTATATTG | AAAACGCA | E3 |
| 5endSL_8mer_052 | TTTCTGTTGGTGCTGATATTGC*CCTCAATT*ACAGTTTCTGTACTATATTG | CCTCAATT | E4 |
| 5endSL_8mer_053 | TTTCTGTTGGTGCTGATATTGC*ACCAATAC*ACAGTTTCTGTACTATATTG | ACCAATAC | E5 |
| 5endSL_8mer_054 | TTTCTGTTGGTGCTGATATTGC*ACCGAGCG*ACAGTTTCTGTACTATATTG | ACCGAGCG | E6 |
| 5endSL_8mer_055 | TTTCTGTTGGTGCTGATATTGC*AAGGACAA*ACAGTTTCTGTACTATATTG | AAGGACAA | E7 |
| 5endSL_8mer_056 | TTTCTGTTGGTGCTGATATTGC*AGGGCTAT*ACAGTTTCTGTACTATATTG | AGGGCTAT | E8 |
| 5endSL_8mer_057 | TTTCTGTTGGTGCTGATATTGC*ACATTATC*ACAGTTTCTGTACTATATTG | ACATTATC | E9 |
| 5endSL_8mer_058 | TTTCTGTTGGTGCTGATATTGC*CACAAAGC*ACAGTTTCTGTACTATATTG | CACAAAGC | E10 |
| 5endSL_8mer_059 | TTTCTGTTGGTGCTGATATTGC*CACGTCGC*ACAGTTTCTGTACTATATTG | CACGTCGC | E11 |
| 5endSL_8mer_060 | TTTCTGTTGGTGCTGATATTGC*TCCGTGAC*ACAGTTTCTGTACTATATTG | TCCGTGAC | E12 |
| 5endSL_8mer_061 | TTTCTGTTGGTGCTGATATTGC*GGAAGTCT*ACAGTTTCTGTACTATATTG | GGAAGTCT | F1 |
| 5endSL_8mer_062 | TTTCTGTTGGTGCTGATATTGC*CGAATTTA*ACAGTTTCTGTACTATATTG | **CGAATTTA** | F2 |
| 5endSL_8mer_063 | TTTCTGTTGGTGCTGATATTGC*TATGTTGC*ACAGTTTCTGTACTATATTG | TATGTTGC | F3 |
| 5endSL_8mer_064 | TTTCTGTTGGTGCTGATATTGC*TTTCTTTT*ACAGTTTCTGTACTATATTG | TTTCTTTT | F4 |
| 5endSL_8mer_065 | TTTCTGTTGGTGCTGATATTGC*ACAACTCT*ACAGTTTCTGTACTATATTG | ACAACTCT | F5 |
| 5endSL_8mer_066 | TTTCTGTTGGTGCTGATATTGC*ATATAGAA*ACAGTTTCTGTACTATATTG | ATATAGAA | F6 |
| 5endSL_8mer_067 | TTTCTGTTGGTGCTGATATTGC*GATGCTGT*ACAGTTTCTGTACTATATTG | GATGCTGT | F7 |
| 5endSL_8mer_068 | TTTCTGTTGGTGCTGATATTGC*GAGTATGC*ACAGTTTCTGTACTATATTG | GAGTATGC | F8 |
| 5endSL_8mer_069 | TTTCTGTTGGTGCTGATATTGC*AATGTCAC*ACAGTTTCTGTACTATATTG | AATGTCAC | F9 |
| 5endSL_8mer_070 | TTTCTGTTGGTGCTGATATTGC*TGCTCTGC*ACAGTTTCTGTACTATATTG | TGCTCTGC | F10 |
| 5endSL_8mer_071 | TTTCTGTTGGTGCTGATATTGC*CTGGGGGT*ACAGTTTCTGTACTATATTG | CTGGGGGT | F11 |
| 5endSL_8mer_072 | TTTCTGTTGGTGCTGATATTGC*CCTTTTAA*ACAGTTTCTGTACTATATTG | CCTTTTAA | F12 |
| 5endSL_8mer_073 | TTTCTGTTGGTGCTGATATTGC*TGGTGCAG*ACAGTTTCTGTACTATATTG | TGGTGCAG | G1 |
| 5endSL_8mer_074 | TTTCTGTTGGTGCTGATATTGC*AGTTTGTA*ACAGTTTCTGTACTATATTG | AGTTTGTA | G2 |
| 5endSL_8mer_075 | TTTCTGTTGGTGCTGATATTGC*GCTGTCTG*ACAGTTTCTGTACTATATTG | GCTGTCTG | G3 |
| 5endSL_8mer_076 | TTTCTGTTGGTGCTGATATTGC*AATCTAGT*ACAGTTTCTGTACTATATTG | AATCTAGT | G4 |
| 5endSL_8mer_077 | TTTCTGTTGGTGCTGATATTGC*GCGATGCA*ACAGTTTCTGTACTATATTG | GCGATGCA | G5 |
| 5endSL_8mer_078 | TTTCTGTTGGTGCTGATATTGC*CCGCGCAG*ACAGTTTCTGTACTATATTG | CCGCGCAG | G6 |
| 5endSL_8mer_079 | TTTCTGTTGGTGCTGATATTGC*ACATTTGA*ACAGTTTCTGTACTATATTG | ACATTTGA | G7 |
| 5endSL_8mer_080 | TTTCTGTTGGTGCTGATATTGC*CGGTGCTG*ACAGTTTCTGTACTATATTG | CGGTGCTG | G8 |
| 5endSL_8mer_081 | TTTCTGTTGGTGCTGATATTGC*TGACTGTC*ACAGTTTCTGTACTATATTG | TGACTGTC | G9 |
| 5endSL_8mer_082 | TTTCTGTTGGTGCTGATATTGC*CTATCTTG*ACAGTTTCTGTACTATATTG | CTATCTTG | G10 |
| 5endSL_8mer_083 | TTTCTGTTGGTGCTGATATTGC*AGCCATAA*ACAGTTTCTGTACTATATTG | AGCCATAA | G11 |
| 5endSL_8mer_084 | TTTCTGTTGGTGCTGATATTGC*CCATCGAA*ACAGTTTCTGTACTATATTG | CCATCGAA | G12 |

| | | | |
|---|---|---|---|
| 5endSL_8mer_085 | TTTCTGTTGGTGCTGATATTGC*CTCGTACT*ACAGTTTCTGTACTATATTG | CTCGTACT | H1 |
| 5endSL_8mer_086 | TTTCTGTTGGTGCTGATATTGC*ATCGTAGC*ACAGTTTCTGTACTATATTG | ATCGTAGC | H2 |
| 5endSL_8mer_087 | TTTCTGTTGGTGCTGATATTGC*GCGCCTTG*ACAGTTTCTGTACTATATTG | GCGCCTTG | H3 |
| 5endSL_8mer_088 | TTTCTGTTGGTGCTGATATTGC*AGCAGGAG*ACAGTTTCTGTACTATATTG | AGCAGGAG | H4 |
| 5endSL_8mer_089 | TTTCTGTTGGTGCTGATATTGC*ACACACCG*ACAGTTTCTGTACTATATTG | ACACACCG | H5 |
| 5endSL_8mer_090 | TTTCTGTTGGTGCTGATATTGC*AGAAGAGG*ACAGTTTCTGTACTATATTG | AGAAGAGG | H6 |
| 5endSL_8mer_091 | TTTCTGTTGGTGCTGATATTGC*TTATTAGA*ACAGTTTCTGTACTATATTG | TTATTAGA | H7 |
| 5endSL_8mer_092 | TTTCTGTTGGTGCTGATATTGC*GGTCTGAA*ACAGTTTCTGTACTATATTG | GGTCTGAA | H8 |
| 5endSL_8mer_093 | TTTCTGTTGGTGCTGATATTGC*AATCAACG*ACAGTTTCTGTACTATATTG | AATCAACG | H9 |
| 5endSL_8mer_094 | TTTCTGTTGGTGCTGATATTGC*ATCAACGA*ACAGTTTCTGTACTATATTG | ATCAACGA | H10 |
| 5endSL_8mer_095 | TTTCTGTTGGTGCTGATATTGC*TATGGAAT*ACAGTTTCTGTACTATATTG | TATGGAAT | H11 |
| 5endSL_8mer_096 | TTTCTGTTGGTGCTGATATTGC*AAGCTAAG*ACAGTTTCTGTACTATATTG | AAGCTAAG | H12 |

Fig.S1

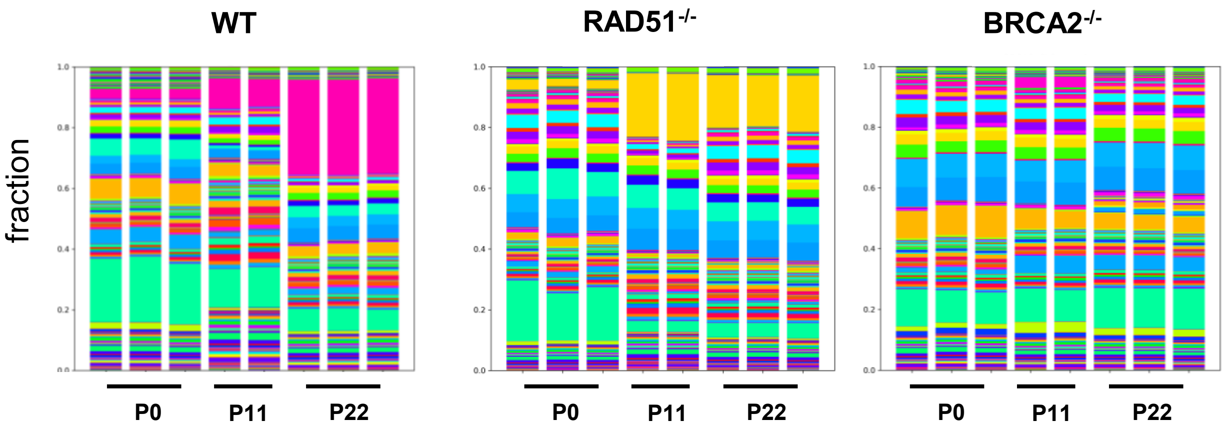

Fig.S2

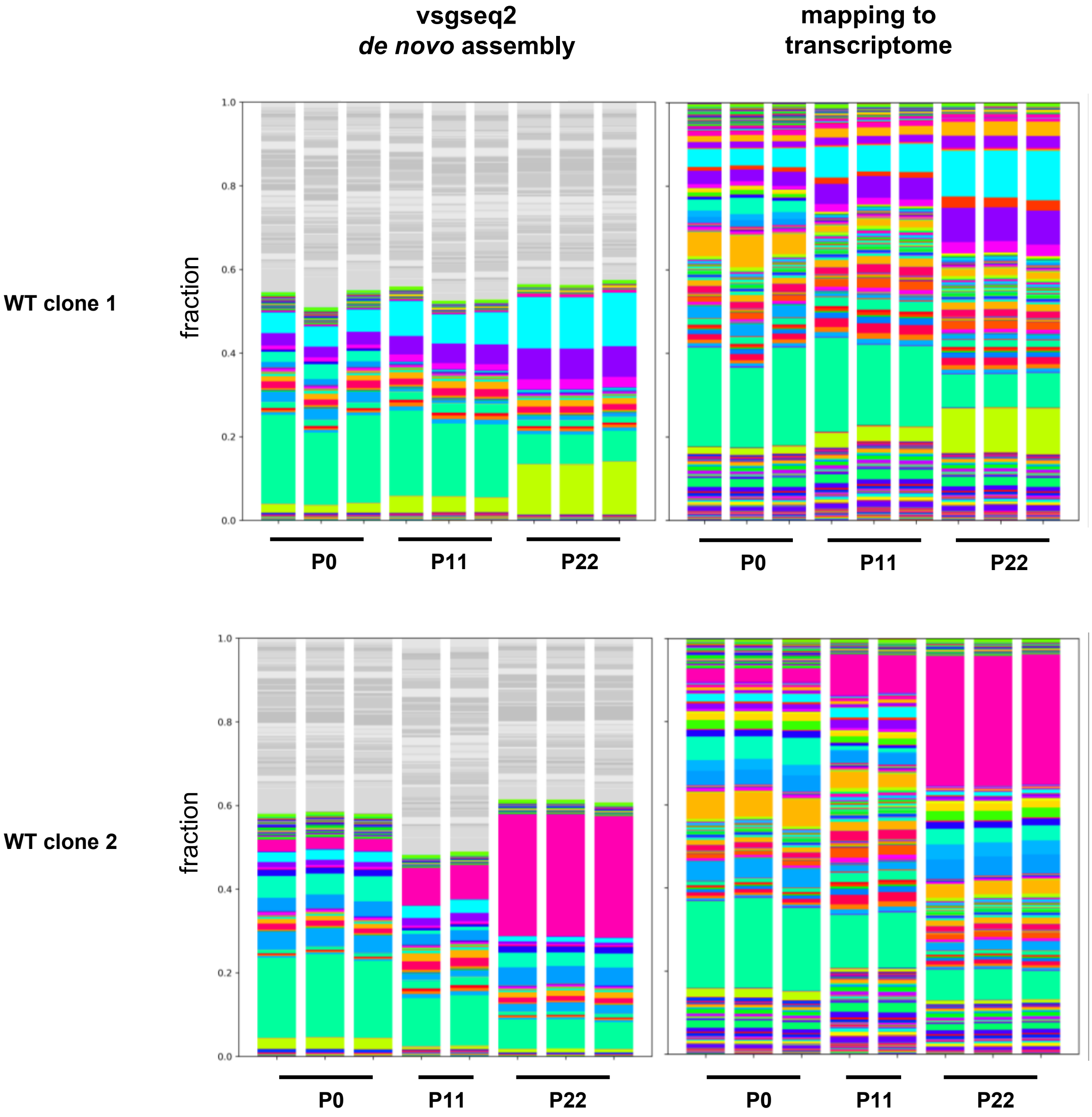
vsgseq2
de novo assembly
mapping to
transcriptome
WT clone 1
fraction
1.0
0.8
0.6
0.4
0.2
0.0
P0
P11
P22
WT clone 2
fraction
1.0
0.8
0.6
0.4
0.2
0.0
P0
P11
P22

Fig.S3

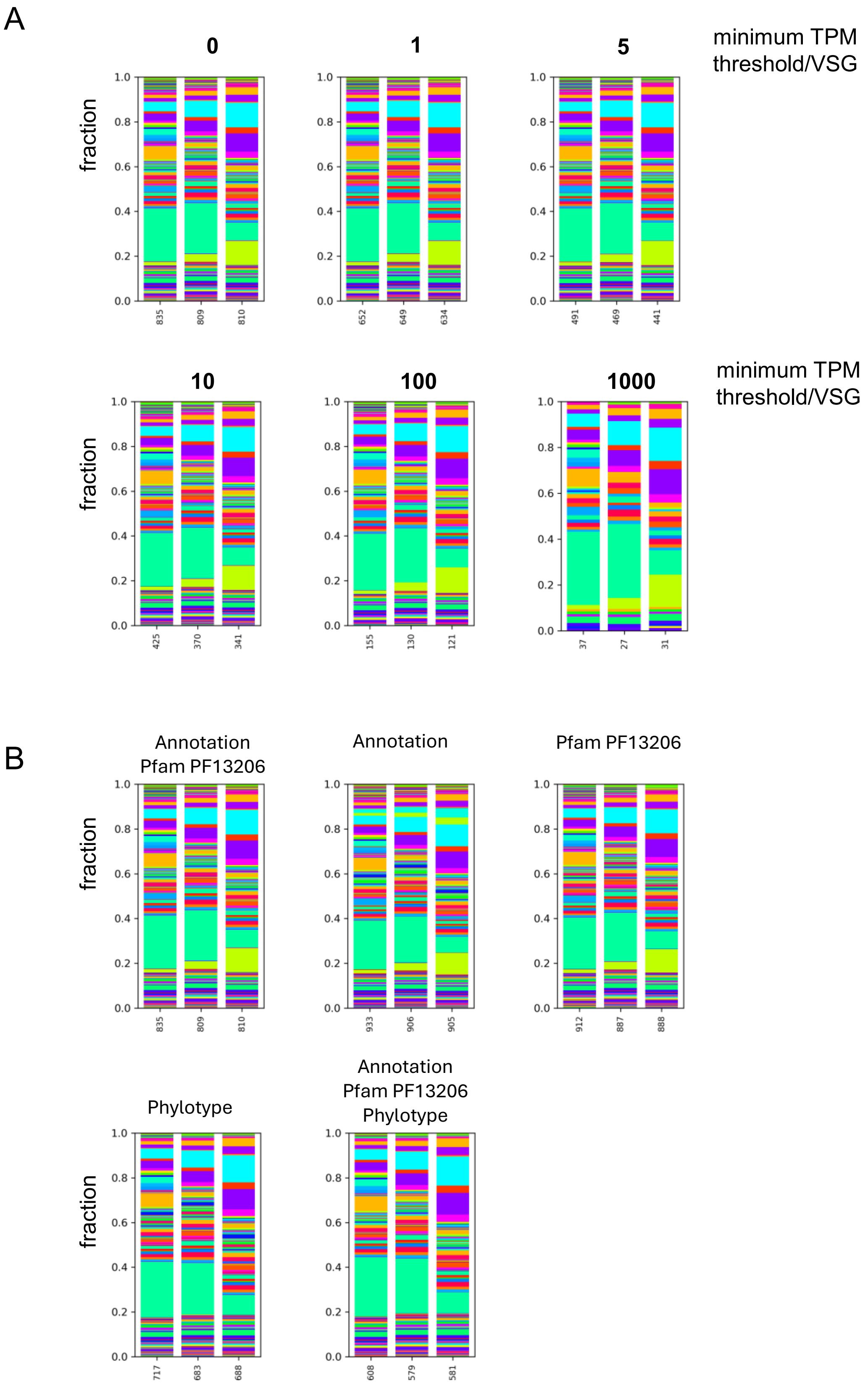
A
minimum TPM threshold/VSG
0
1
5
10
100
1000
fraction
1.0
0.8
0.6
0.4
0.2
0.0
835
809
810
652
649
634
491
469
441
425
370
341
155
130
121
37
27
31
B
Annotation Pfam PF13206
Annotation
Pfam PF13206
Phylotype
Annotation Pfam PF13206 Phylotype
835
809
810
933
906
905
912
887
888
717
683
688
608
579
581

Fig.S4

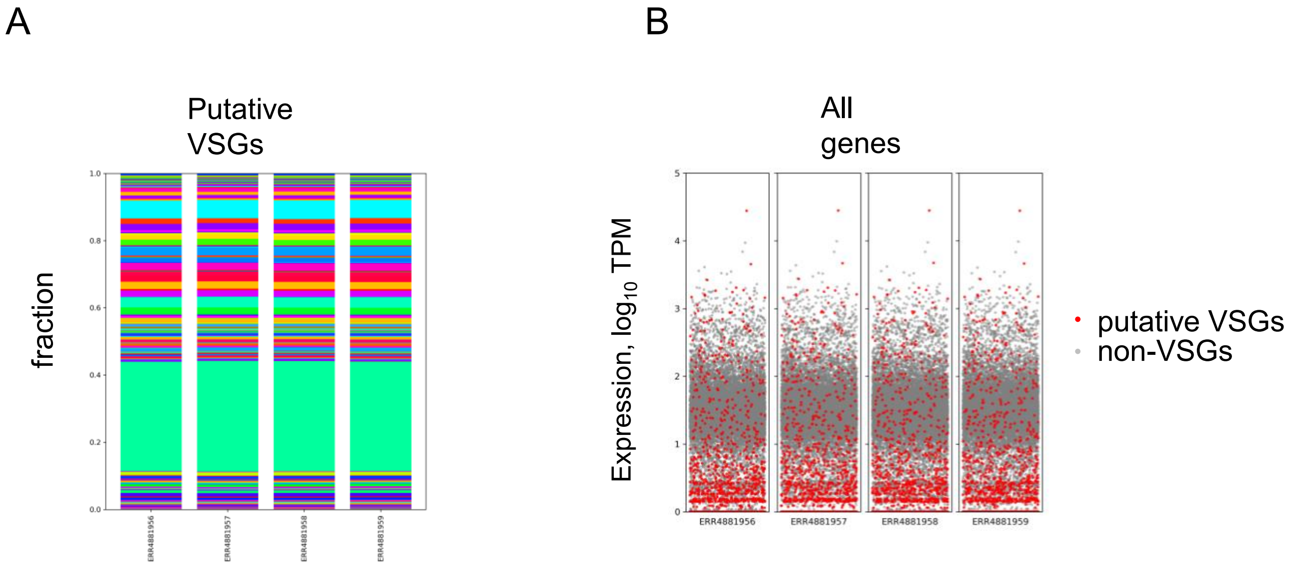
A
Putative
VSGs
fraction
1.0
0.8
0.6
0.4
0.2
0.0
ERR4881956
ERR4881957
ERR4881958
ERR4881959
B
All
genes
Expression, log10 TPM
5
4
3
2
1
0
ERR4881956
ERR4881957
ERR4881958
ERR4881959
putative VSGs
non-VSGs

Fig.S5

**C10** **G03**

Lysed Memb sVSG Lysed Memb sVSG

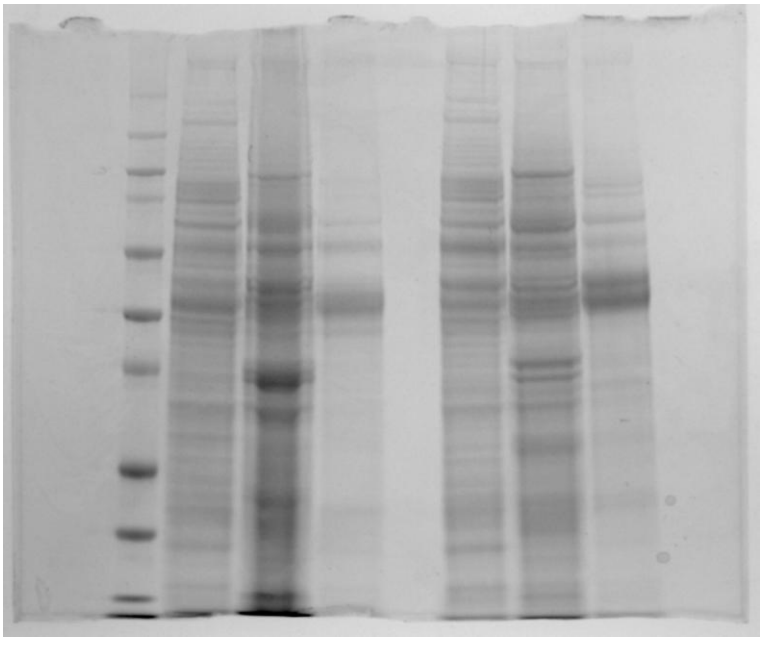

Fig.S6

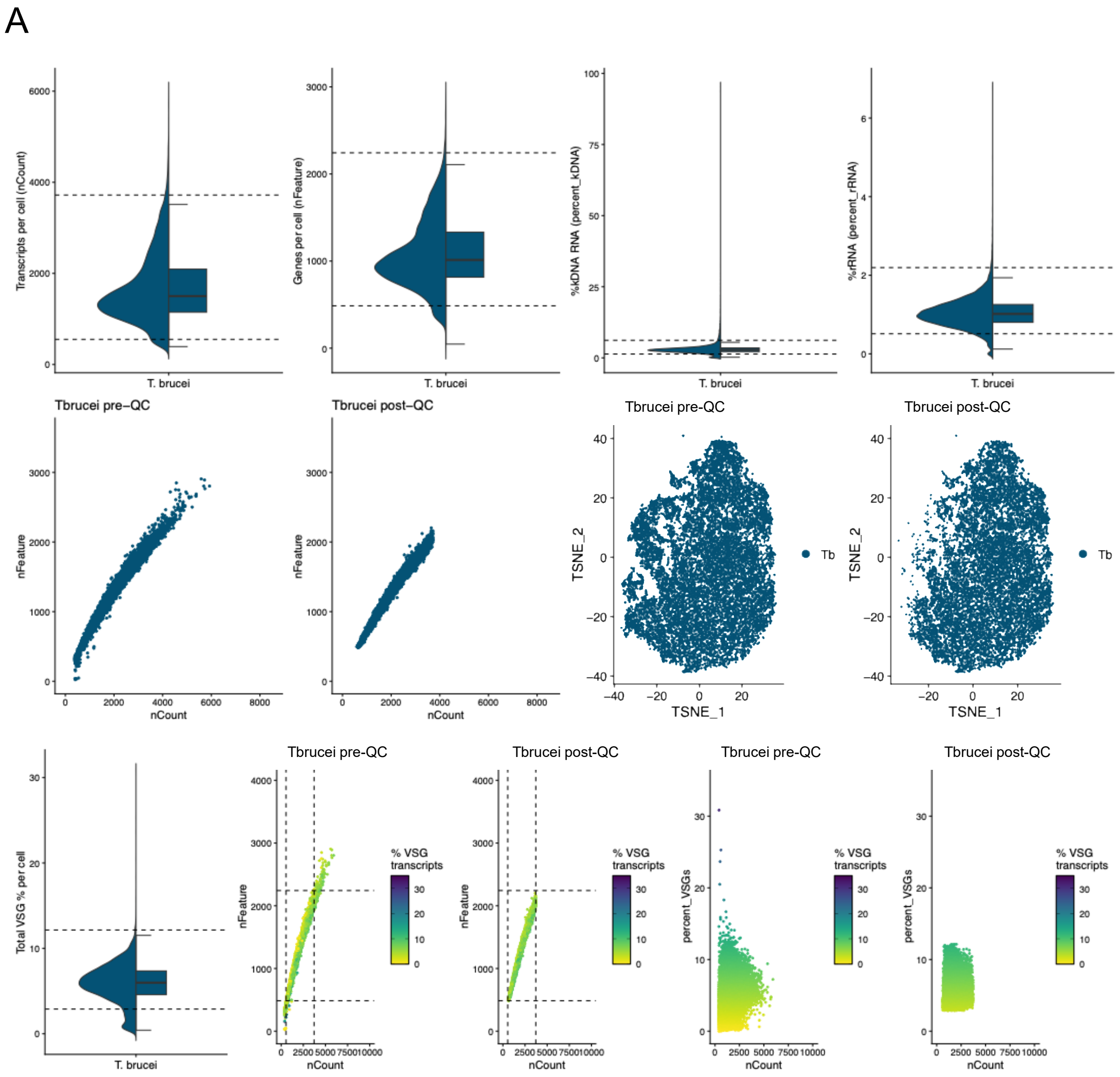
A
Transcripts per cell (nCount)
T. brucei
Genes per cell (nFeature)
T. brucei
%kDNA RNA (percent_kDNA)
T. brucei
%rRNA (percent_rRNA)
T. brucei
Tbrucei pre–QC
nFeature
nCount
Tbrucei post–QC
nFeature
nCount
Tbrucei pre-QC
TSNE_2
TSNE_1
Tb
Tbrucei post-QC
TSNE_2
TSNE_1
Tb
Total VSG % per cell
T. brucei
Tbrucei pre-QC
nFeature
nCount
% VSG transcripts
Tbrucei post-QC
nFeature
nCount
% VSG transcripts
Tbrucei pre-QC
percent_VSGs
nCount
% VSG transcripts
Tbrucei post-QC
percent_VSGs
nCount
% VSG transcripts

Fig.S6 (continue)

B

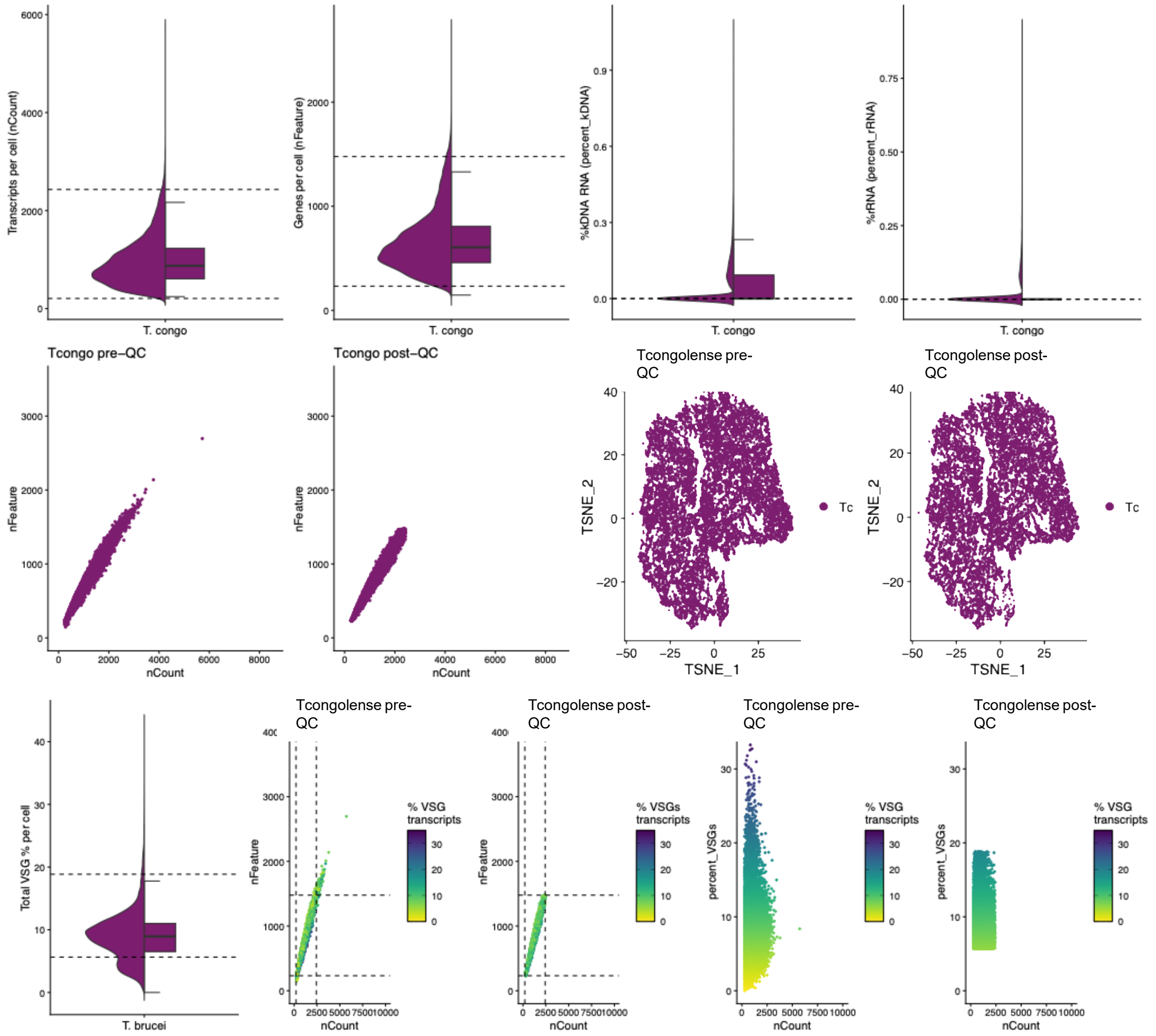

Fig.S7

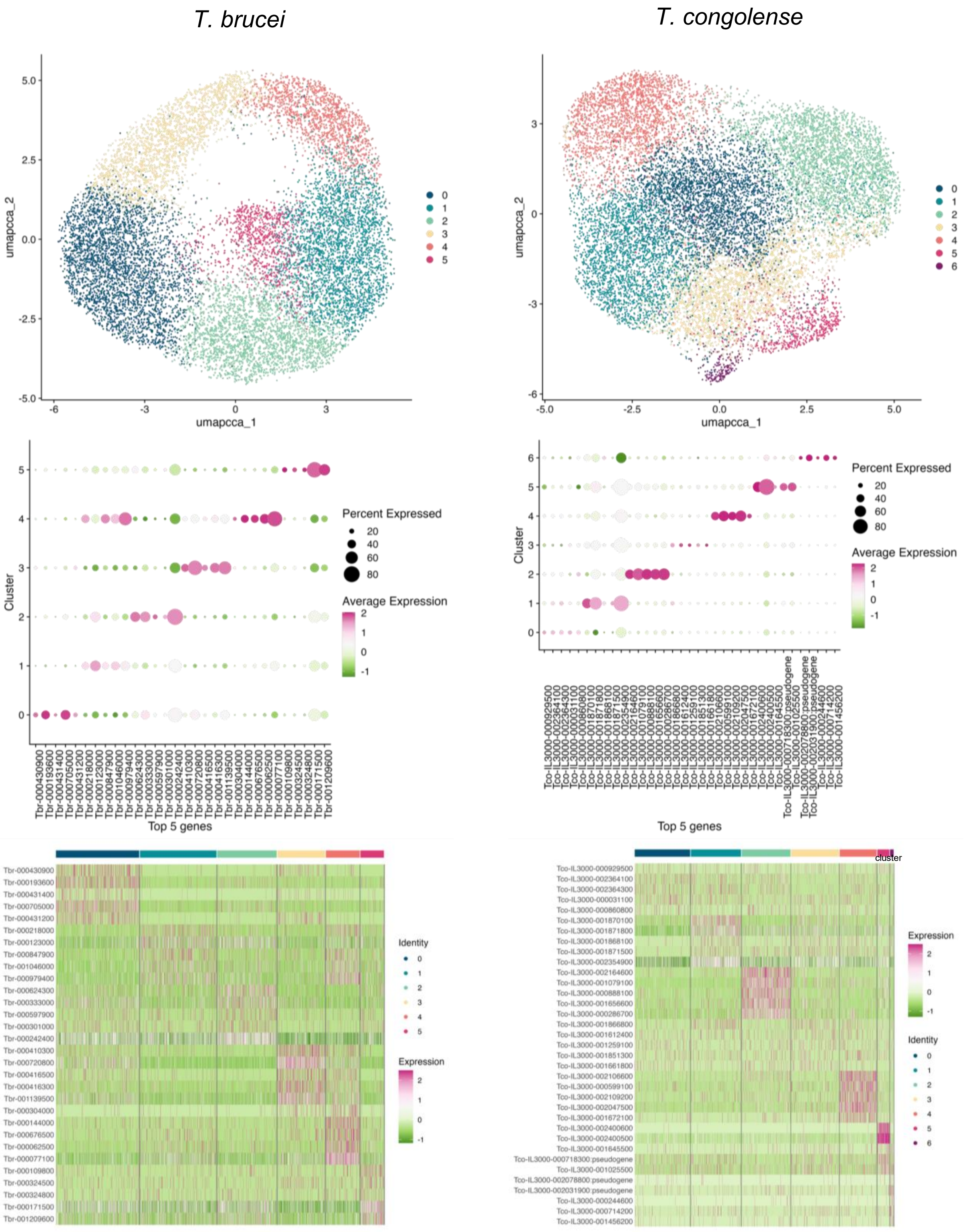
T. brucei
T. congolense
umapcca_1
umapcca_2
Cluster
Top 5 genes
Percent Expressed
20
40
60
80
Average Expression
Identity
Expression
cluster

Fig.S8

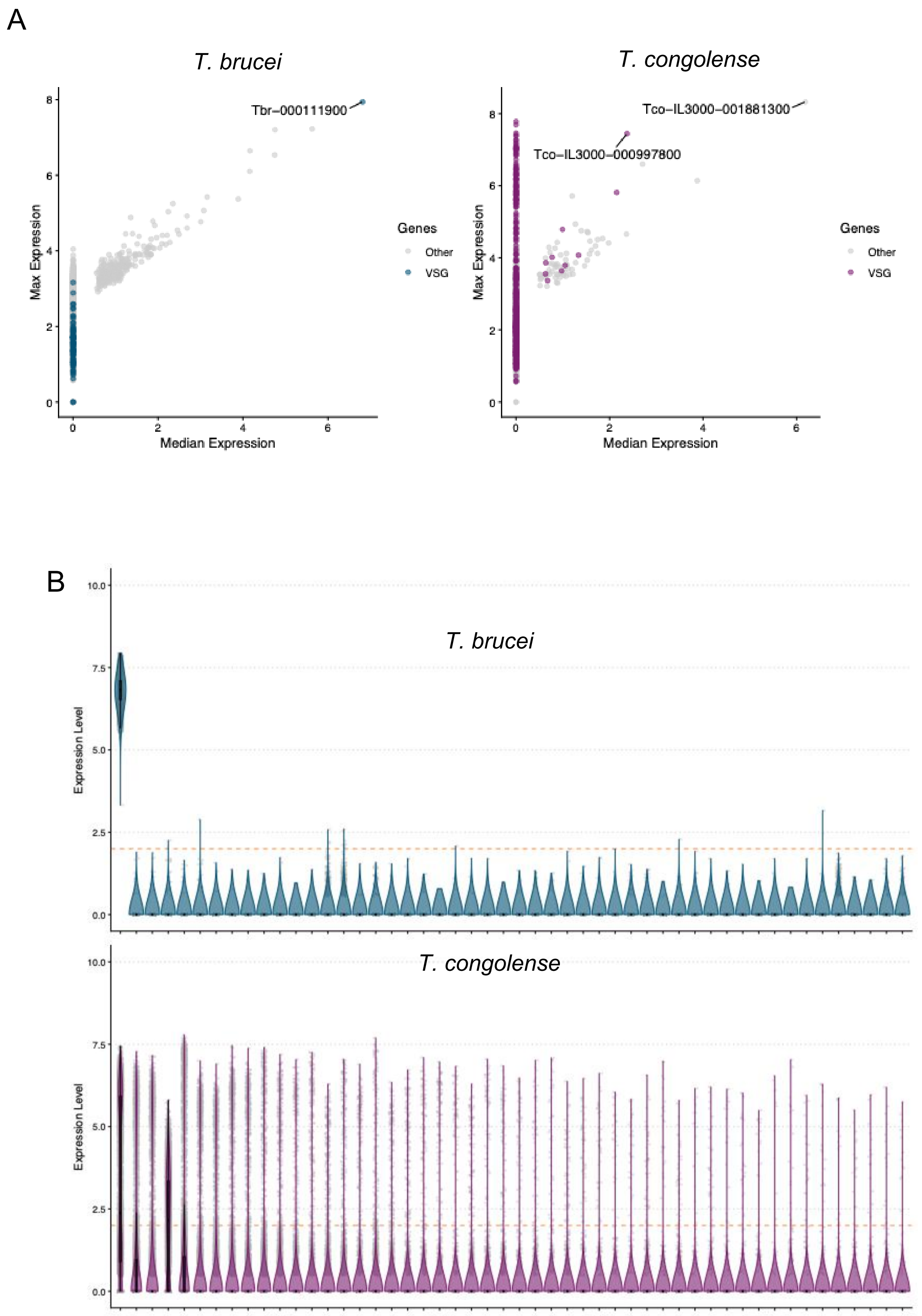

A
T. brucei
Tbr−000111900
Max Expression
Median Expression
Genes
Other
VSG
T. congolense
Tco−IL3000−001881300
Tco−IL3000−000997800
Max Expression
Median Expression
Genes
Other
VSG
B
T. brucei
Expression Level
T. congolense
Expression Level

Fig.S9

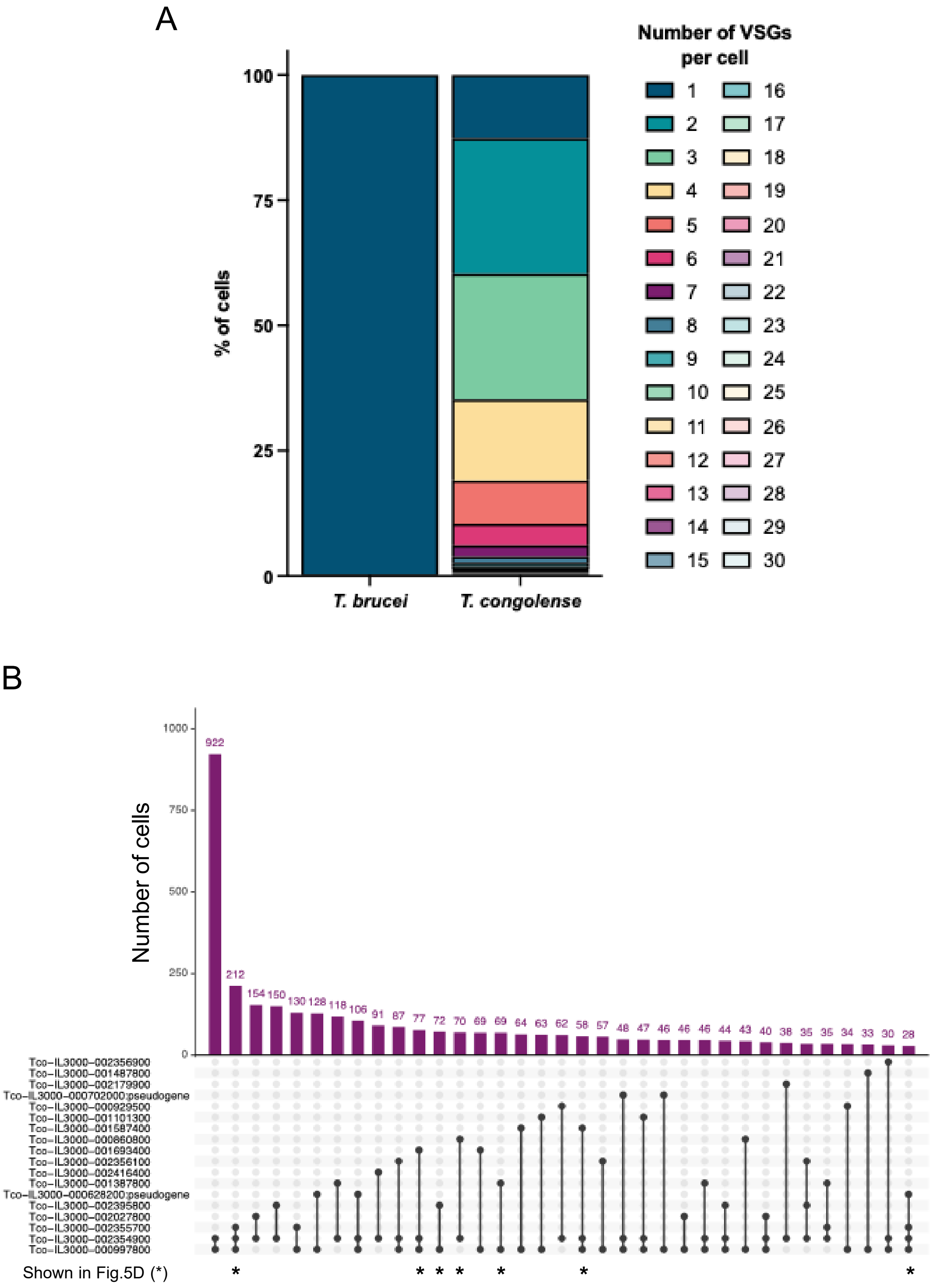
A
Number of VSGs per cell
% of cells
100
75
50
25
0
T. brucei
T. congolense
B
Number of cells
1000
750
500
250
0
922
212
154
150
130
128
118
106
91
87
77
72
70
69
69
64
63
62
58
57
48
47
46
46
46
44
43
40
38
35
35
34
33
30
28
Tco-IL3000-002356900
Tco-IL3000-001487800
Tco-IL3000-002179900
Tco-IL3000-000702000:pseudogene
Tco-IL3000-000929500
Tco-IL3000-001101300
Tco-IL3000-001587400
Tco-IL3000-000860800
Tco-IL3000-001693400
Tco-IL3000-002356100
Tco-IL3000-002416400
Tco-IL3000-001387800
Tco-IL3000-000628200:pseudogene
Tco-IL3000-002395800
Tco-IL3000-002027800
Tco-IL3000-002355700
Tco-IL3000-002354900
Tco-IL3000-000997800
Shown in Fig.5D (*)
* * * * * * *

Fig.S10

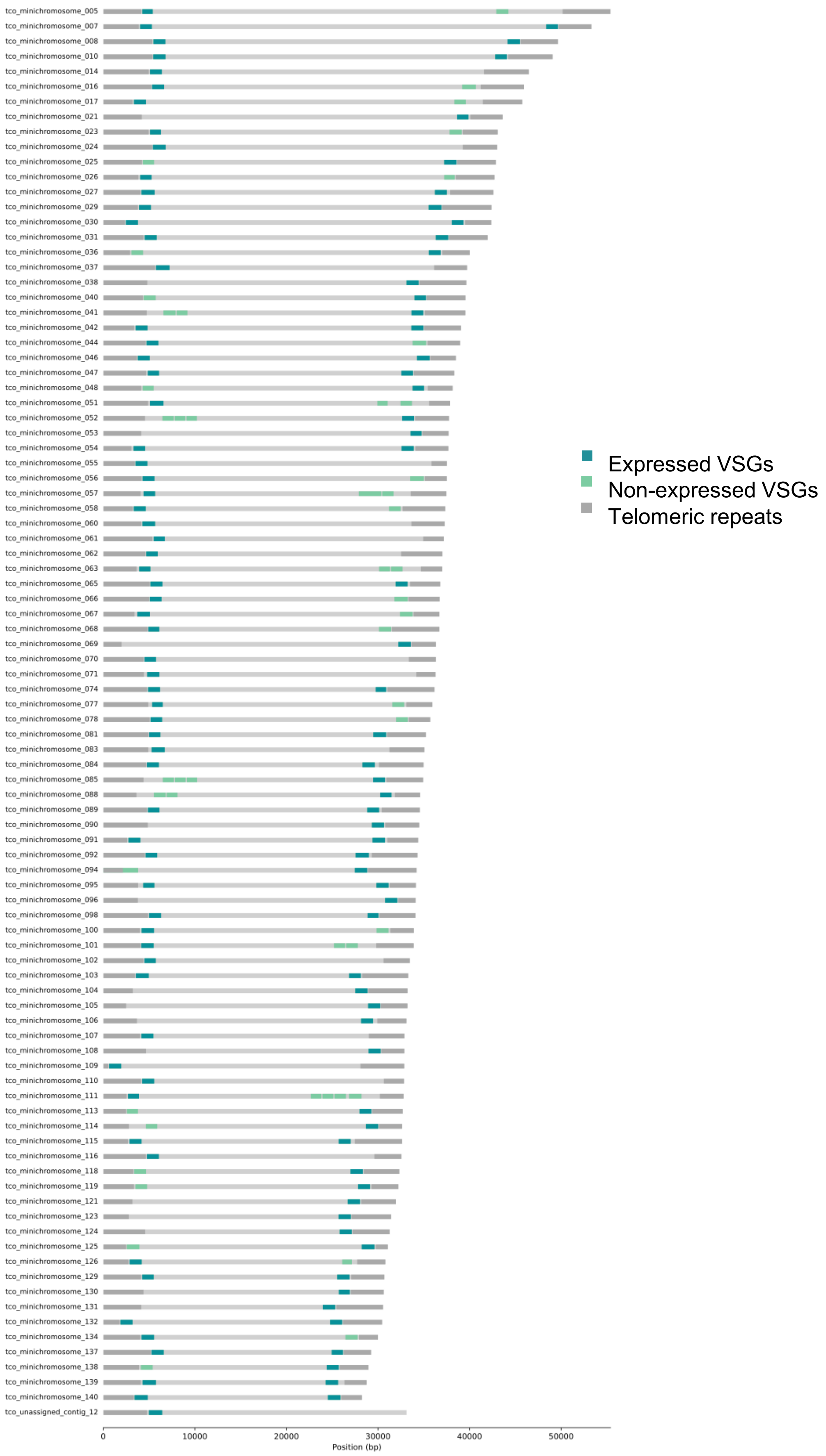
tco_minichromosome_005
tco_minichromosome_007
tco_minichromosome_008
tco_minichromosome_010
tco_minichromosome_014
tco_minichromosome_016
tco_minichromosome_017
tco_minichromosome_021
tco_minichromosome_023
tco_minichromosome_024
tco_minichromosome_025
tco_minichromosome_026
tco_minichromosome_027
tco_minichromosome_029
tco_minichromosome_030
tco_minichromosome_031
tco_minichromosome_036
tco_minichromosome_037
tco_minichromosome_038
tco_minichromosome_040
tco_minichromosome_041
tco_minichromosome_042
tco_minichromosome_044
tco_minichromosome_046
tco_minichromosome_047
tco_minichromosome_048
tco_minichromosome_051
tco_minichromosome_052
tco_minichromosome_053
tco_minichromosome_054
tco_minichromosome_055
tco_minichromosome_056
tco_minichromosome_057
tco_minichromosome_058
tco_minichromosome_060
tco_minichromosome_061
tco_minichromosome_062
tco_minichromosome_063
tco_minichromosome_065
tco_minichromosome_066
tco_minichromosome_067
tco_minichromosome_068
tco_minichromosome_069
tco_minichromosome_070
tco_minichromosome_071
tco_minichromosome_074
tco_minichromosome_077
tco_minichromosome_078
tco_minichromosome_081
tco_minichromosome_083
tco_minichromosome_084
tco_minichromosome_085
tco_minichromosome_088
tco_minichromosome_089
tco_minichromosome_090
tco_minichromosome_091
tco_minichromosome_092
tco_minichromosome_094
tco_minichromosome_095
tco_minichromosome_096
tco_minichromosome_098
tco_minichromosome_100
tco_minichromosome_101
tco_minichromosome_102
tco_minichromosome_103
tco_minichromosome_104
tco_minichromosome_105
tco_minichromosome_106
tco_minichromosome_107
tco_minichromosome_108
tco_minichromosome_109
tco_minichromosome_110
tco_minichromosome_111
tco_minichromosome_113
tco_minichromosome_114
tco_minichromosome_115
tco_minichromosome_116
tco_minichromosome_118
tco_minichromosome_119
tco_minichromosome_121
tco_minichromosome_123
tco_minichromosome_124
tco_minichromosome_125
tco_minichromosome_126
tco_minichromosome_129
tco_minichromosome_130
tco_minichromosome_131
tco_minichromosome_132
tco_minichromosome_134
tco_minichromosome_137
tco_minichromosome_138
tco_minichromosome_139
tco_minichromosome_140
tco_unassigned_contig_12
0
10000
20000
30000
40000
50000
Position (bp)
Expressed VSGs
Non-expressed VSGs
Telomeric repeats

Fig.S11

A

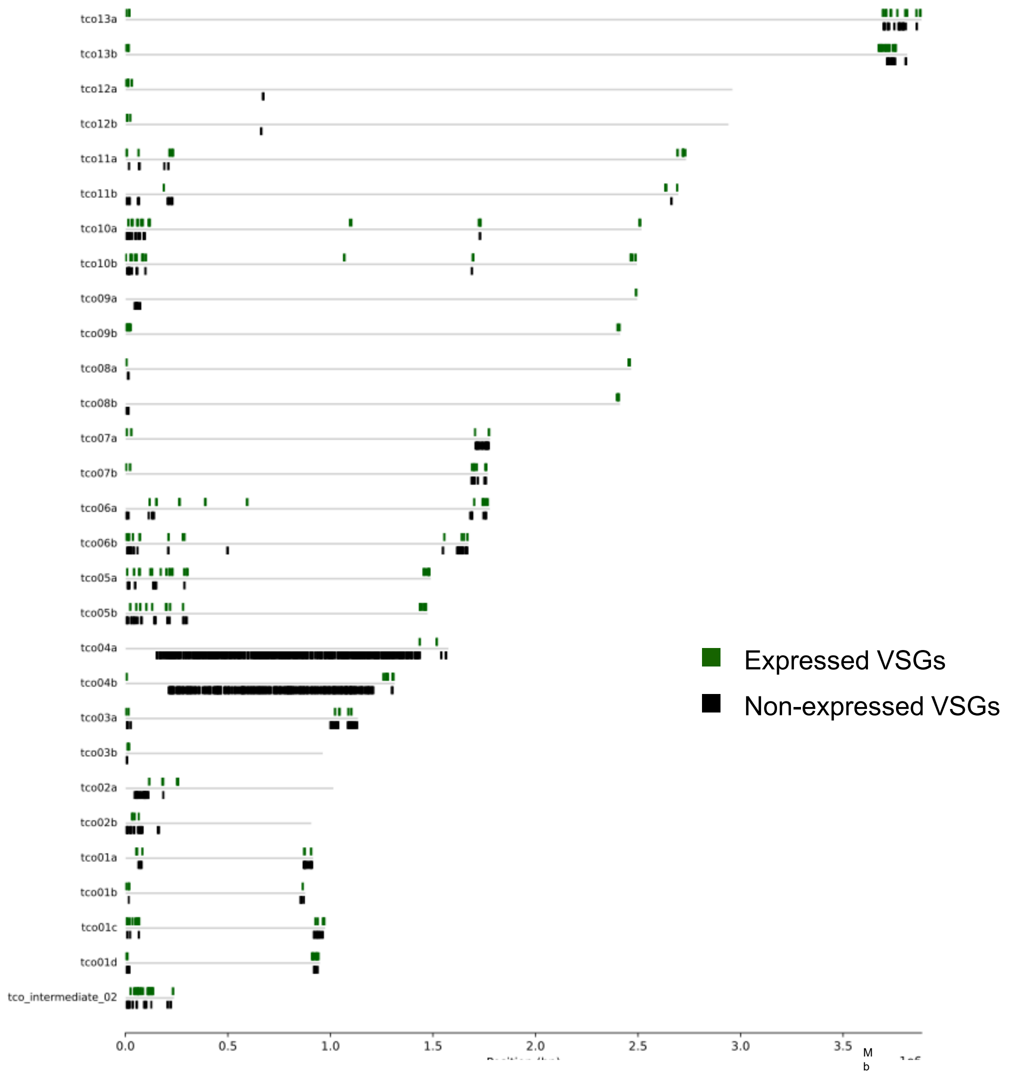

tco13a
tco13b
tco12a
tco12b
tco11a
tco11b
tco10a
tco10b
tco09a
tco09b
tco08a
tco08b
tco07a
tco07b
tco06a
tco06b
tco05a
tco05b
tco04a
tco04b
tco03a
tco03b
tco02a
tco02b
tco01a
tco01b
tco01c
tco01d
tco_intermediate_02
0.0
0.5
1.0
1.5
2.0
2.5
3.0
3.5
Mb
Expressed VSGs
Non-expressed VSGs

B

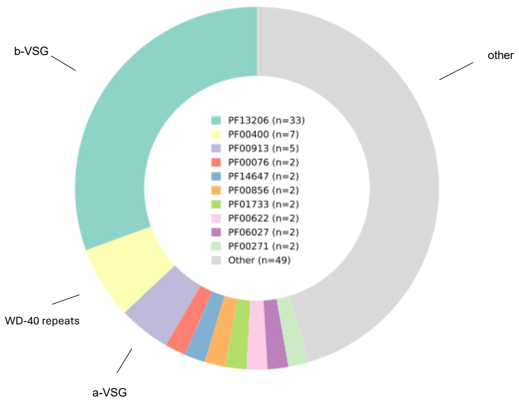

b-VSG
other
PF13206 (n=33)
PF00400 (n=7)
PF00913 (n=5)
PF00076 (n=2)
PF14647 (n=2)
PF00856 (n=2)
PF01733 (n=2)
PF00622 (n=2)
PF06027 (n=2)
PF00271 (n=2)
Other (n=49)
WD-40 repeats
a-VSG

Fig.S12

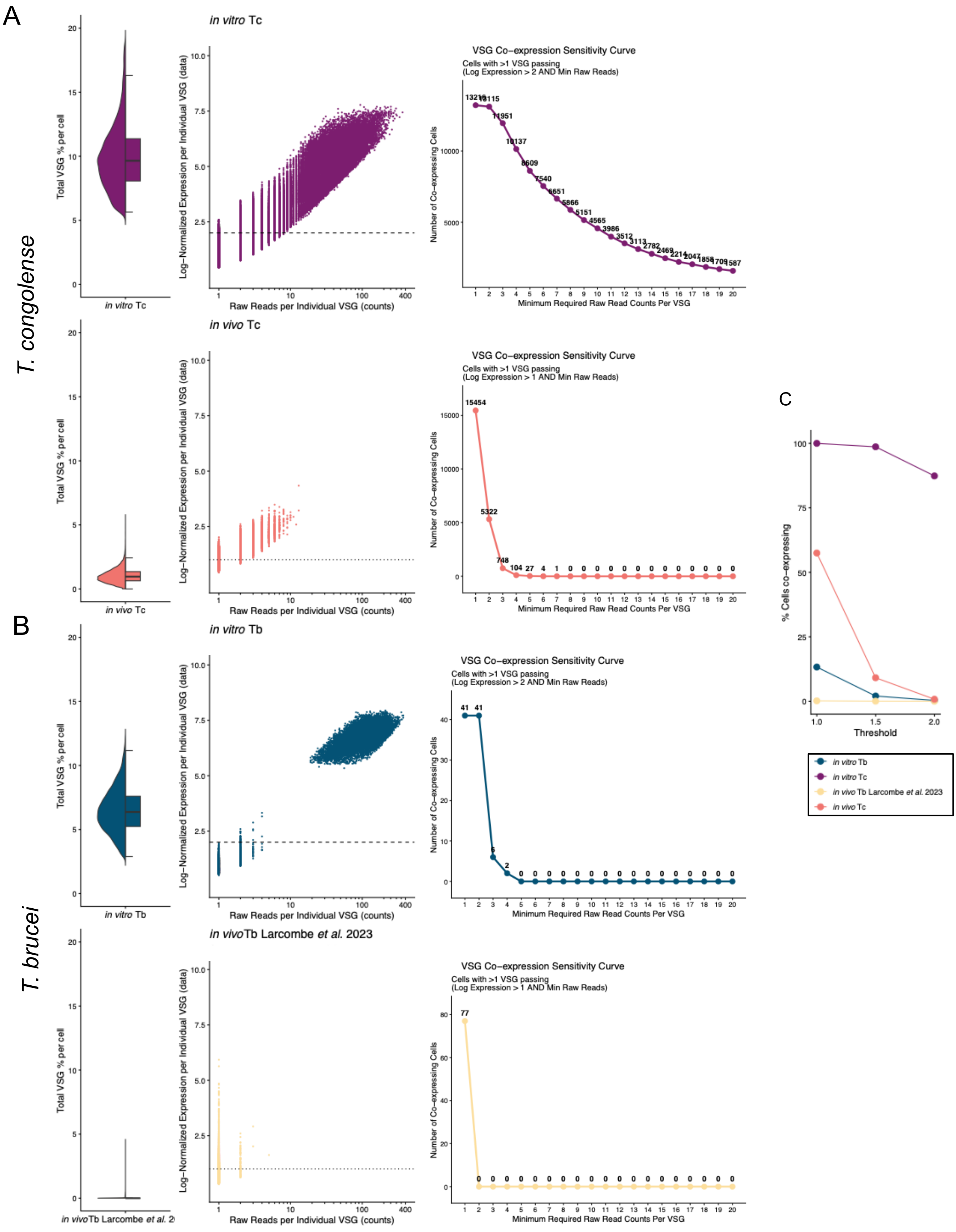
A
B
C
T. congolense
T. brucei
in vitro Tc
in vivo Tc
in vitro Tb
in vivo Tb Larcombe et al. 2023
Total VSG % per cell
Log–Normalized Expression per Individual VSG (data)
Raw Reads per Individual VSG (counts)
VSG Co–expression Sensitivity Curve
Cells with >1 VSG passing
(Log Expression > 2 AND Min Raw Reads)
(Log Expression > 1 AND Min Raw Reads)
Number of Co–expressing Cells
Minimum Required Raw Read Counts Per VSG
% Cells co–expressing
Threshold

Fig.S13

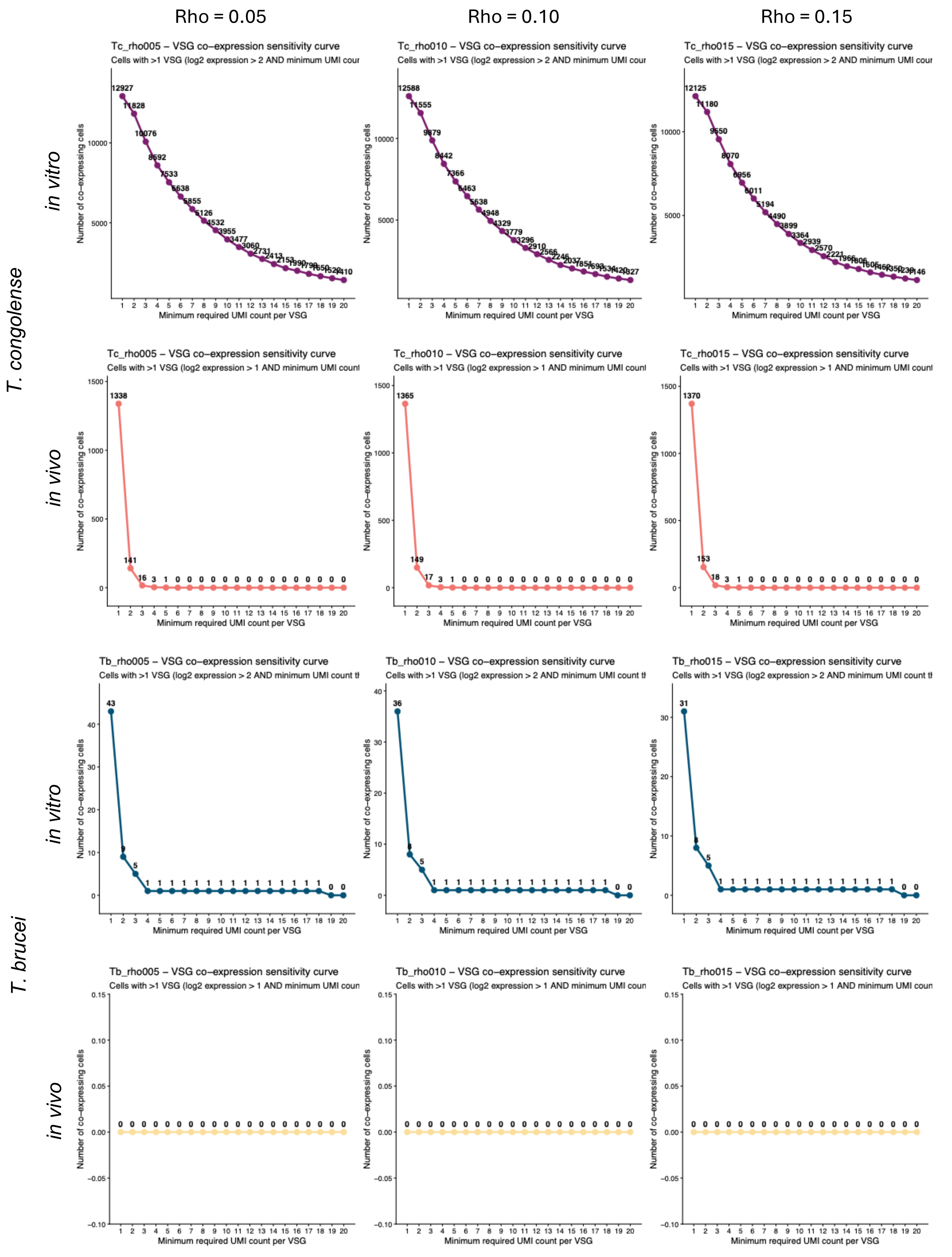
Rho = 0.05
Rho = 0.10
Rho = 0.15
T. congolense
T. brucei
in vitro
in vivo
in vitro
in vivo
Tc_rho005 – VSG co–expression sensitivity curve
Cells with >1 VSG (log2 expression > 2 AND minimum UMI cour
Tc_rho010 – VSG co–expression sensitivity curve
Cells with >1 VSG (log2 expression > 2 AND minimum UMI cour
Tc_rho015 – VSG co–expression sensitivity curve
Cells with >1 VSG (log2 expression > 2 AND minimum UMI cour
Tc_rho005 – VSG co–expression sensitivity curve
Cells with >1 VSG (log2 expression > 1 AND minimum UMI count
Tc_rho010 – VSG co–expression sensitivity curve
Cells with >1 VSG (log2 expression > 1 AND minimum UMI count
Tc_rho015 – VSG co–expression sensitivity curve
Cells with >1 VSG (log2 expression > 1 AND minimum UMI count
Tb_rho005 – VSG co–expression sensitivity curve
Cells with >1 VSG (log2 expression > 2 AND minimum UMI count th
Tb_rho010 – VSG co–expression sensitivity curve
Cells with >1 VSG (log2 expression > 2 AND minimum UMI count th
Tb_rho015 – VSG co–expression sensitivity curve
Cells with >1 VSG (log2 expression > 2 AND minimum UMI count th
Tb_rho005 – VSG co–expression sensitivity curve
Cells with >1 VSG (log2 expression > 1 AND minimum UMI coun
Tb_rho010 – VSG co–expression sensitivity curve
Cells with >1 VSG (log2 expression > 1 AND minimum UMI coun
Tb_rho015 – VSG co–expression sensitivity curve
Cells with >1 VSG (log2 expression > 1 AND minimum UMI coun
Number of co–expressing cells
Minimum required UMI count per VSG
